# Engineering SIRPα conformational plasticity to reveal a cryptic pocket suitable for structure-based drug design

**DOI:** 10.64898/2025.12.10.693509

**Authors:** M Storder, S Barelier, F Cordier, T Yacoub, L Ilari, K Barral, S Mahmoodi, M Saez-Ayala, S Combes, S Betzi, C Derviaux, A Ulliana, F Torres, J Rubin, P Roche, X Morelli, ED Garcin, TW Miller

## Abstract

The protein-protein interaction between Signal Regulatory Protein alpha (SIRPα) and CD47 is a critical immune checkpoint that enables tumor immune escape, making it a key target for cancer immunotherapy. While antibody-based therapies exist, the development of small-molecule inhibitors has been hindered by the flat, featureless binding interface. Here, we report the discovery of a novel, druggable cryptic pocket within the SIRPα D1 domain (the WYF pocket), revealed through a structure-based fragment screening campaign using x-ray crystallography. This pocket, defined by residues Trp38, Tyr50, and Phe74, is only accessible in a conformation that is incompatible with CD47 binding, making it a candidate for structure-based drug design and immune checkpoint inhibitor development.

Through a combination of NMR spectroscopy, molecular dynamics simulations, and biophysical assays, we demonstrate that access to this cryptic site is dynamically controlled by a single “gatekeeper” residue, Gln52. The rotameric state of Gln52 dictates a conformational equilibrium between a “closed,” state and a ligand-accessible “open” state. We validated this mechanism by engineering SIRPα mutants to bias this equilibrium. A Q52F mutation locked the pocket in a closed state, abolishing both CD47 and fragment binding, while Q52A and Q52R mutations biased the protein toward an open state. These “open-biased” mutants not only exhibited decreased affinity for CD47 but also significantly improved binding to small-molecule fragments that inhibit the SIRPα-CD47 interaction.

This work reveals the intrinsic conformational plasticity of SIRPα and establishes a validated structure-based roadmap for a new class of allosteric inhibitors. This ‘flexibility-for-inhibition’ strategy functions by trapping a non-binding conformation and represents a broadly applicable framework for targeting this and other challenging immune checkpoints.

## INTRODUCTION

The protein-protein complex comprised of Signal Regulatory Protein alpha (SIRPα) and cluster of differentiation-47 (CD47) is a critical immune checkpoint implicated in tumor immune escape and microenvironment remodeling. SIRPα is a transmembrane protein consisting of three extracellular immunoglobulin superfamily (IgSF) domains (D1, D2, D3), a transmembrane segment, and a cytoplasmic tail containing immunoreceptor tyrosine-based inhibition motifs (ITIMs) that recruit the phosphatases SHP-1 and SHP-2 upon ligand binding (**Figure 1A**). The N-terminal Ig-like D1 domain (SIRPα-D1) binds CD47 via four loops that form the binding interface: the BC (Thr28-Pro35), C’D (Asn51-Phe57), DE (Ser64-Asn71), and FG (Lys96-Asp100) loops (**Figure 1B**)[1]. Within these loops, 13 residues directly engage CD47 via 29 hydrogen bonds and 10 salt bridges[2]. Among these interactions, SIRPα DE loop Ser66 hydrogen bonds to CD47 pyroglutamate residue, and C’D loop Gln52 side chain hydrogen bonds to CD47 Glu104 main chain nitrogen. SIRPα-D1 exhibits significant polymorphism although this does not significantly alter overall CD47 binding affinity[1,3]. The two predominant human allelic variants, SIRPαV1 (NCBI XP_047295872.1) and SIRPαV2 (NCBI XP_054179025.1)[4], share approximately 97% sequence identity across the full-length protein as well as the residues of the CD47 interface (**Figure 1C**).

**Figure 1.**
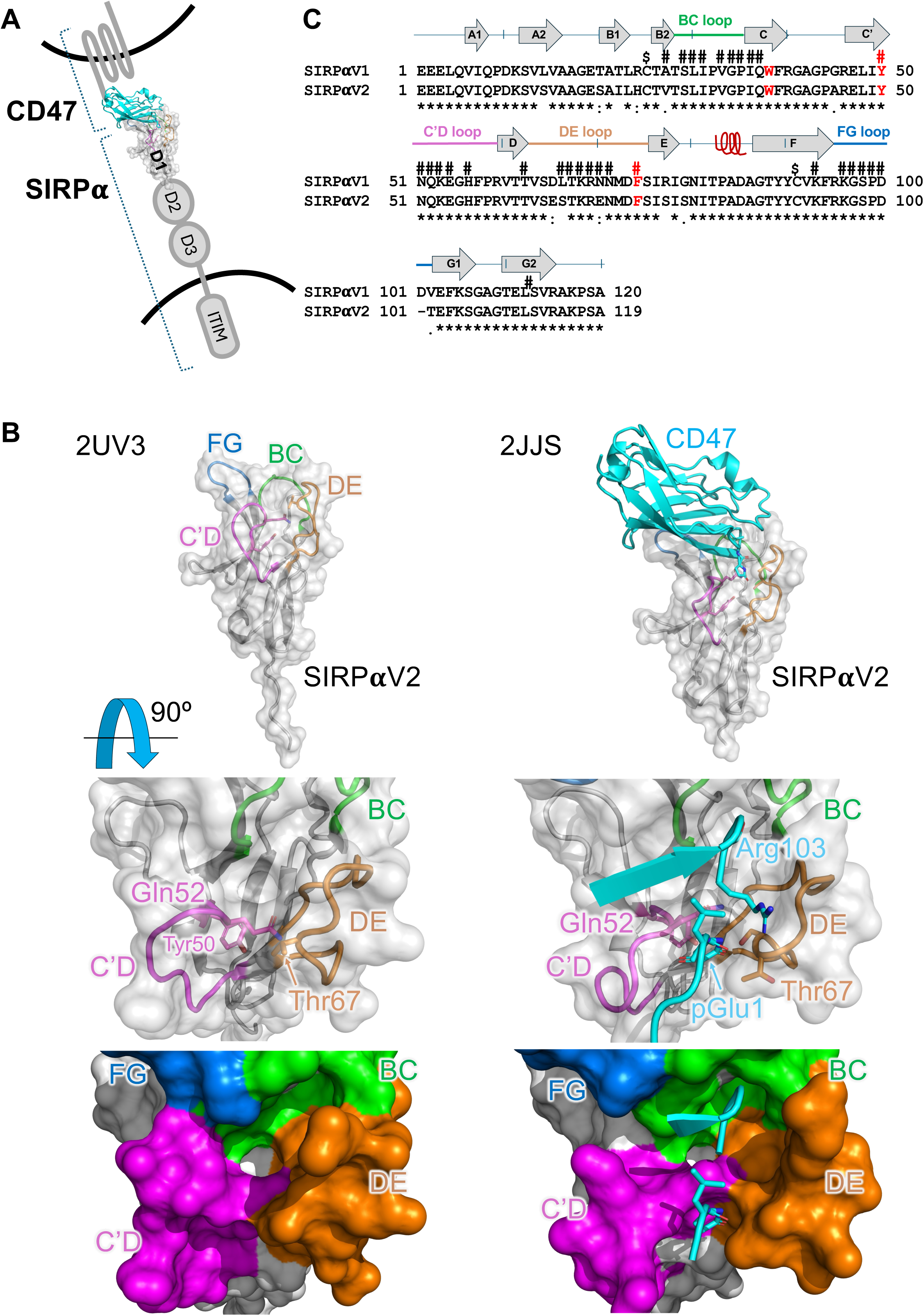
The CD47-SIRPα immune checkpoint. **(A)** Schematic of SIRPα-CD47 interaction. The interacting extracellular CD47 (cyan cartoon) and the D1 domain of SIRPαV2 (white surface) are represented (from PDB entry 2JJS). Cell membranes are represented by curved black lines. **(B)** Structural comparison between SIRPα alone (PDB entry 2UV3) and in complex with CD47 (PDB entry 2JJS). Top: overall view of the 3D structure. Middle view: zoom on the regions involved in CD47 interaction. Bottom: accessible surface highlighting the conformational changes in the C’D and DE loops upon CD47 binding. Loops involved in the interaction with CD47 are colored as in (**C**). Figures were made with Pymol. **(C)** Amino acid sequence alignment of the D1 domains for SIRPαV1 and SIRPαV2 variants. Secondary structures (PDB entry 2UV3) are shown above the sequence. Residues involved in the interaction with CD47 are highlighted (#). The disulfide bridge between Cys25 and 91 is shown ($). Aromatic amino acids forming the cryptic WYF pocket are highlighted in red. The sequence alignment was done with CLUSTAL Omega.

Considerable research efforts by academic and pharmaceutical groups are aimed at discovering inhibitors targeting this interaction and yielded anti-CD47 antibodies, which have progressed to phase 3 clinical trials[5]. However, these therapeutic agents are associated with numerous adverse side effects attributable to the ubiquitous expression of CD47, including on erythrocytes and leukocytes[6–8]. In contrast, agents targeting SIRPα may present a safer therapeutic strategy, as SIRPα expression is predominantly restricted to myeloid cells. Previous investigations involving anti-SIRPα antibodies have demonstrated comparatively lower toxicity profiles relative to anti-CD47 agents[8,9]. Moreover, recent research has underscored the prognostic significance of SIRPα-positive tumor-associated macrophages (TAMs) as indicators of unfavorable outcomes and therapeutic resistance, independent of tumor CD47 expression[10]. Consequently, direct targeting of SIRPα is gaining traction as a potentially safer and more efficacious immunotherapeutic strategy[11–13].

To date, preclinically validated small molecule inhibitors that directly block the SIRPα-CD47 interaction do not exist despite the accumulation of detailed structural information, the existence of high-throughput screening assays[14], and several chemical screening campaigns[15]. Importantly, this strategy has many potential advantages compared to antibodies, including more flexible pharmacokinetics to minimize on-target toxicity, the potential to create therapeutic combinations, and enhanced solid tumor penetration (reviewed in [16]). Here, we describe the first phase of a rational approach to generate SIRPα-targeted inhibitors of this immune checkpoint using structure-based drug design. We have discovered a cryptic druggable pocket in a CD47-non-binding conformation and described the dynamics of its formation. Furthermore, we reveal validated small molecule fragment ligands discovered by x-ray crystallography that bind to the SIRPα cryptic pocket and block CD47 interaction as starting points for chemical optimization. Lastly, we demonstrate that a key Gln residue acts as the gatekeeper of the cryptic pocket, and that opening of the pocket can be modulated by a single-point mutation.

## RESULTS

### Fragment Screening Reveals a Cryptic Pocket in SIRPαV2 and Identifies Initial Ligands

To identify chemical starting points for creating high-affinity ligands that modulate SIRPα function, we initiated a structure-based fragment screening campaign. Due to the extensive homology of the CD47 binding interface in SIRPαV1 and SIRPαV2 variants, as well as difficulties crystallizing SIRPαV1 without CD47, we focused our initial studies on the SIRPαV2 variant. Robust crystallization conditions were first established for the D1 domain of human SIRPαV2 (SIRPαV2-D1), differing from those used in previously reported structures. These conditions allowed for rapid and reproducible protein crystallization (2-3 days) that resulted in high-quality diffraction (<1.8Å).

The structure of SIRPαV2-D1 alone was solved at 1.6Å resolution (**Table S1**). The final model contained 3 mol/ASU and included 323 water molecules, and 7 zinc atoms from the crystallization condition. The three molecules displayed overall similar structures (RMSD ranging from 0.25 to 0.48 Å), with the largest deviations observed in the C’D, DE and FG loops (RMSD ranging from 0.7 to 2.5 Å). Additionally, the largest difference with the previously solved structure of SIRPαV2-D1 (2UV3) were located in the C’D, DE, and FG loops (RMSD ranging from 1 to 3.7Å).

We initiated a structure-based drug design program by screening a combined library of over 800 low-molecular-weight fragments (< 300 Da) sourced from the DSI-poised library[17] using x-ray crystallography via the high-throughput XChem platform (Diamond Light Source)[19]. A total of 697 x-ray datasets were collected from crystals soaked with fragments (typically at 20 mM), including 619 unique compounds and 78 DMSO/control datasets. The control datasets were combined into a composite “ground state” electron density map to allow detection of bound fragments with low occupancy with PANDDA[20]. Analysis revealed 27 structures containing bound fragments (’hits’) that were further refined. Notably, 19 of these 27 hits clustered within a hydrophobic pocket delineated by residues Trp38, Tyr50, and Phe74 (hereafter referred to as the WYF pocket) in only one of the three SIRPα molecules in the asymmetric unit (**Suppl. Fig. S1, Figure 2A-D**). Among the hits with high occupancy was compound x0408, an indole derivative, exemplifying the shared structural motif that consisted of an aromatic ring bound in the WYF pocket in an edge-to-face π interaction[21] with Tyr50 and Phe74 (**Figure 2A-C**). Superimposition of the structure bound to compound x0408 on the structure of SIRPαV2 alone yielded an overall rmsd of 0.43 Å (**Figure 2D**). The largest differences were located in regions 41-46, 51-56 (C’D loop), 66-70 (DE loop), and 96-99 (FG loop). In addition, the side chain of Gln52 adopted a different rotameric conformation, where it no longer hydrogen bonded to the Ile36 main chain (“Gln52-in”), but instead hydrogen bonded to Tyr50 hydroxyl (“Gln52-out”).

**Figure 2.**
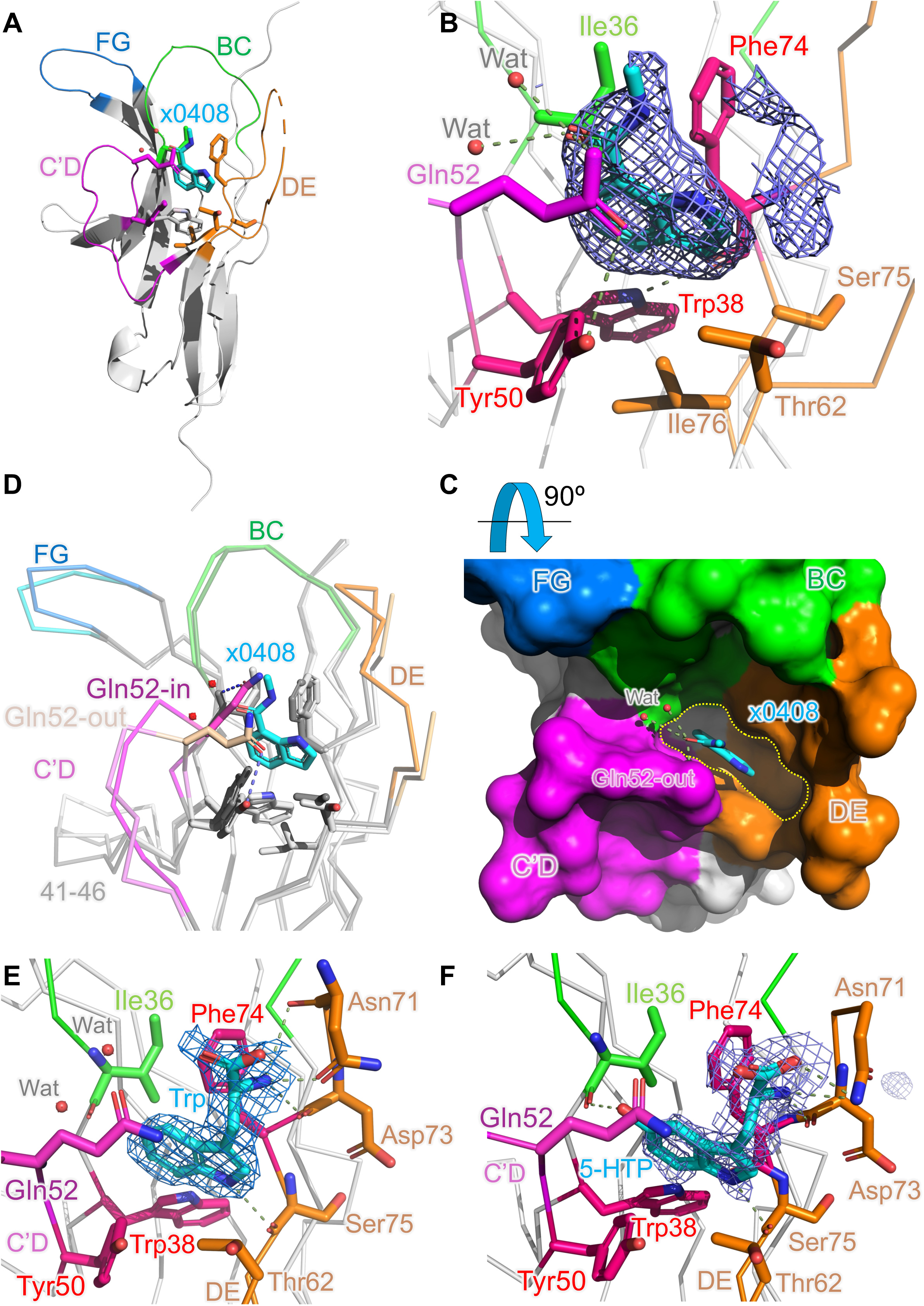
Fragment screening reveals the cryptic WYF pocket. (A) Overall x-ray structure of SIRPαV2 with fragment x0408 (cyan sticks), shown in the same orientation and same coloring as in Fig. 1. **(B)** Zoom on the WYF pocket with bound x0408 fragment (cyan) and event map contoured at 1 sigma (blue mesh). Interactions between x0408 and the protein and solvent (Wat) are represented in green dashed lines. **(C)** Rotated view showing the WYF pocket (yellow dotted outline) with bound x0408 compound (cyan) with SIRPαV2 represented as a surface (colors as in (**A**)). Hydrogen bonds from x0408 to water molecules (Wat) or Gln52 side chain are represented in green dashed lines. **(D)** Superimposition of empty SIRPαV2 structure (dark color) and SIRPαV2 bound to fragment x0408 (light color). View and colors are the same as in (**A**). In the empty SIRPαV2 structure, the Gln52 side chain hydrogen bonds to Ile36 main chain amide (“Gln52-in”). In the SIRPαV2-x0408 complex structure, the Gln52 side chain swings out and hydrogen bonds to the Tyr50 hydroxyl (“Gln52-out”). **(E)** X-ray structure of SIRPαV2 bound to Trp (cyan sticks) with 2Fo-Fc electron density map contoured at 1 sigma in the same orientation as (**B**). **(F)** X-ray structure of SIRPαV2 bound to 5-HTP (cyan sticks) with 2Fo-Fc electron density map contoured at 0.5 sigma in the same orientation as (**B**). Interactions between ligands and protein residues are indicated by dotted lines.

Screening of the iSCB fragment library[18] via crystal soaking yielded additional hits, including the metabolites L-Tryptophan (Trp) and 5-hydroxy-tryptophan (5-HTP), which shared the bicyclic indole core with x0408 and occupied the same binding pocket (**Figure 2E,F**). Identification of multiple fragments sharing similar indole-like pharmacophoric features and binding to the WYF pocket strongly suggested the presence of a druggable site not readily apparent in previously described SIRPα structures.

These crystallographic hits were characterized and validated through functional and biophysical assays. To assess their potential to disrupt the SIRPα-CD47 interaction, we employed our previously validated AlphaLISA assay[14]. Seventeen distinct hits, repurchased from validated commercial sources, were tested at concentrations up to 10 mM. Thirteen of these fragments demonstrated greater than 50% inhibition of the SIRPα-CD47 interaction (**Table S2, Suppl. Fig. S2**). Trp and 5-HTP inhibited the interaction with IC_50_ values of 6.0 ± 1.3 mM and 2.0 ± 0.3 mM, respectively, while x0408 was the most potent of the indole analogs at 0.5 ± 0.1 mM (**Table 1**).

**Table 1.** IC_50_ and K_D_ for WT and mutant SIRPα proteins.

|  | Trp (mM) | 5-HTP (mM) | x0408 (mM) | CD47 (μM) |
| --- | --- | --- | --- | --- |
| **IC<sub>50</sub> AlphaLISA** |  |  |  |  |
| SIRPαV2-CD47 | 6.0 ± 1.3 | 2.0 ± 0.3 | 0.5 ± 0.1 |  |
| **K<sub>D</sub> SPR** |  |  |  |  |
| SIRPαV2 | 13.5 ± 1.2 | 7.8 ± 0.8 | NA | 0.4 ± 0.0 |
| SIRPαV1 | 45 ± 19 | 18.6 ± 2.8 | NA | 0.3 ± 0.0 |
| **K<sub>D</sub> ITC** |  |  |  |  |
| SIRPαV2 WT | 8.1 ± 0.3 | 8.8 ± 0.9 | NA | 0.4 ± 0.1 |
| SIRPαV2 Q52A | NA | 4.6 ± 0.2 | NA | 5.4 ± 1.3 |
| SIRPαV2 Q52R | 3.0 ± NA | 1.9 ± 0.4 | NA | 3.8 ± 0.8 |
| SIRPαV2 Q52K | 3.8 ± NA | 2.3 ± NA | NA | 80.0 ± 40.5 |
| SIRPαV2 Q52F | No binding | No binding | NA | No binding |
| **K<sub>D</sub> CIDNP NMR** |  |  |  |  |
| SIRPαV2 WT |  | - |  |  |
| SIRPαV2 Q52A |  | >10 |  |  |
| SIRPαV2 Q52R |  | 2.3 ± 0.7 |  |  |
| **HSQC NMR (CSP)** |  |  |  |  |
| SIRPαV2 | Positive | Positive | Positive |  |
| SIRPαV1 | Negative | Negative | Negative |  |

| IC<sub>50</sub> SIRPα competition WT vs Q52A (AlphaLISA) | SIRPαV2 WT (μM) | SIRPαV2 Q52A (μM) | SIRPαV2 Q52R (μM) |
| --- | --- | --- | --- |
| SIRPαV2 WT-CD47 | 0.7 ± 0.1 | 9.3 ± 1.4 | 5.0 ± 0.5 |
| SIRPαV2 Q52A-CD47 | 0.1 ± 0.0 | 2.8 ± 0.1 | 2.8 ± 0.2 |
| SIRPαV2 Q52R-CD47 | 0.5 ± 0.1 | 2.9 ± 0.2 | 3.8 ± 0.2 |
| IC<sub>50</sub> SIRPα competition WT vs Q52A (HTRF) | SIRPαV2 WT (μM) | SIRPαV2 Q52A (μM) | SIRPαV2 Q52R (μM) |
| SIRPαV2 WT-CD47 | 1.9 ± 0.2 | 26.3 ± 4.1 | 31.2 ± 7.5 |
| SIRPαV2 Q52A-CD47 | 0.3 ± 0.0 | 2.4 ± 0.3 | 3.9 ± 0.5 |
| SIRPαV2 Q52R-CD47 | 0.4 ± 0.1 | 3.1 ± 0.3 | 3.3 ± 0.5 |

Given the low affinities of the fragment hits and to confirm the activity of weak hits using an orthogonal biophysical assay, we used Nuclear Magnetic Resonance (NMR) spectroscopy. This technique allowed us to confirm direct binding of the fragments to SIRPα in solution and verify their binding sites. As a prerequisite, we successfully obtained the first reported backbone chemical shift assignments for the ^15^N-^13^C-labeled D1 domains of both human SIRPαV1 and SIRPαV2 (**Suppl. Fig. S3A,B**). The largest amide chemical shift differences (Δδ) between the two isoforms were localized in the vicinity of the amino acid variants and covered most of the DE loop (**Suppl. Fig. S3C,D**). Initial ^1^H−^15^N Heteronuclear Single Quantum Coherence (HSQC) spectra were recorded on ^15^N−labeled SIRPαV2 in the presence of 50-fold molar excess of fragments. For the indole-derivatives x0408, Trp, and 5−HTP, significant ^1^H/^15^N Chemical Shift Perturbations (CSPs) were measured in the region of the WYF pocket (particularly Ile36, Trp38, Tyr50, Gln52, Ser64, Phe74), indicative of direct binding (**Table S2**, **Suppl. Fig. S4)**. Other fragments showed no significant CSPs under these conditions, likely reflecting very weak affinities. Similarly, no significant CSPs were measured in HSQC spectra recorded on ^15^N−labeled SIRPαV1 with 50-fold molar excess of Trp and 5-HTP, suggesting weak affinity for this variant in these conditions (data not shown).

Binding of Trp and 5-HTP to SIRPαV2 and SIRPαV1 was further validated using surface plasmon resonance (SPR) spectroscopy (**Table 1, Suppl. Fig. S5**). For SIRPαV2, binding affinities were in general agreement with their IC_50_. For SIRPαV1, these data confirmed the existence of the WYF pocket, with an approximately 2 to 3-fold lower affinity for the ligands, confirming the NMR data. The x0408 compound was not tested due to incompatibility of the experimental system with DMSO.

We subsequently used Isothermal Titration Calorimetry (ITC) to quantify the binding affinity (K_D_) of Trp and 5-HTP to WT SIRPαV2. We showed that Trp and 5-HTP bound weakly to WT SIRPαV2 (K_D_ of 8.1 ± 0.3 mM and 8.8 ± 0.9 mM, respectively; **Table 1, Suppl. Fig. S6)**. Both results were in general agreement with the measured IC_50_ values and SPR-based affinity measurements.

### Intrinsic Loop Dynamics Drive an Induced-Fit Mechanism for Ligand Recognition in the SIRPα WYF Pocket

To characterize the molecular determinants of WYF-pocket formation and estimate its occurrence in SIRPα intrinsic dynamics, we used a combination of NMR spectroscopy and computational molecular dynamics simulations.

As revealed above, the WYF pocket-bearing conformation required concerted movement of the C’D and DE loops and a specific rotameric state of Gln52 (Gln52-out conformation). We analyzed backbone dynamics of SIRPαV1 and V2 variants on the ps-ns (order parameter S^2^) and the μs-ms (exchange contribution R_ex_) timescales by ^15^N relaxation measurements. We found significant differences in the dynamics of the C’D and DE loops between the two variants. For SIRPαV2, the region around DE loop residue Ser66 displayed a low order parameter (S^2^ < 0.6), indicative of high flexibility, whereas the backbone region near C’D loop residue Gln52 displayed high order parameter (S^2^ >0.8), indicative of low flexibility (**Figure 3A-C, Suppl. Fig. S7A**). These results agree with our crystal structure (PDB entry 9TF5), where Gln52 hydrogen bonds to Ile36, thereby restricting WYF pocket access, while the DE loop adopts a more open conformation (**Figure 2D**). For SIRPαV1, regions around Leu66 and Gln52 both showed higher order parameters (S^2^ > 0.8) and no conformational exchange (R_ex_ ∼ 0), both indicative of low flexibility (**Suppl. Fig. S7A-E)**. This observation agrees with the crystal structure of SIRPαV1 (PDB entry 6NMU), in which Leu66 is oriented towards the WYF pocket and engages in hydrophobic interactions with the Gln52 side chain, thereby stabilizing a closed DE loop. Additionally, prediction of backbone dynamics from the experimental backbone chemical shifts using TALOS-N[22] was in agreement with the experimental data above, with significantly higher predicted flexibility for the SIRPαV2 DE loop compared to SIRPαV1 (order parameters S^2,pred^ < 0.55, and ∼0.75-0.8, respectively; **Suppl. Fig. S7F-H**). Thus, NMR data revealed that the SIRPαV2 DE loop was more dynamic on the fast timescale than SIRPαV1. This could explain the 3-fold increase in 5-HTP binding affinity in SIRPαV2 as measured by SPR.

**Figure 3:**
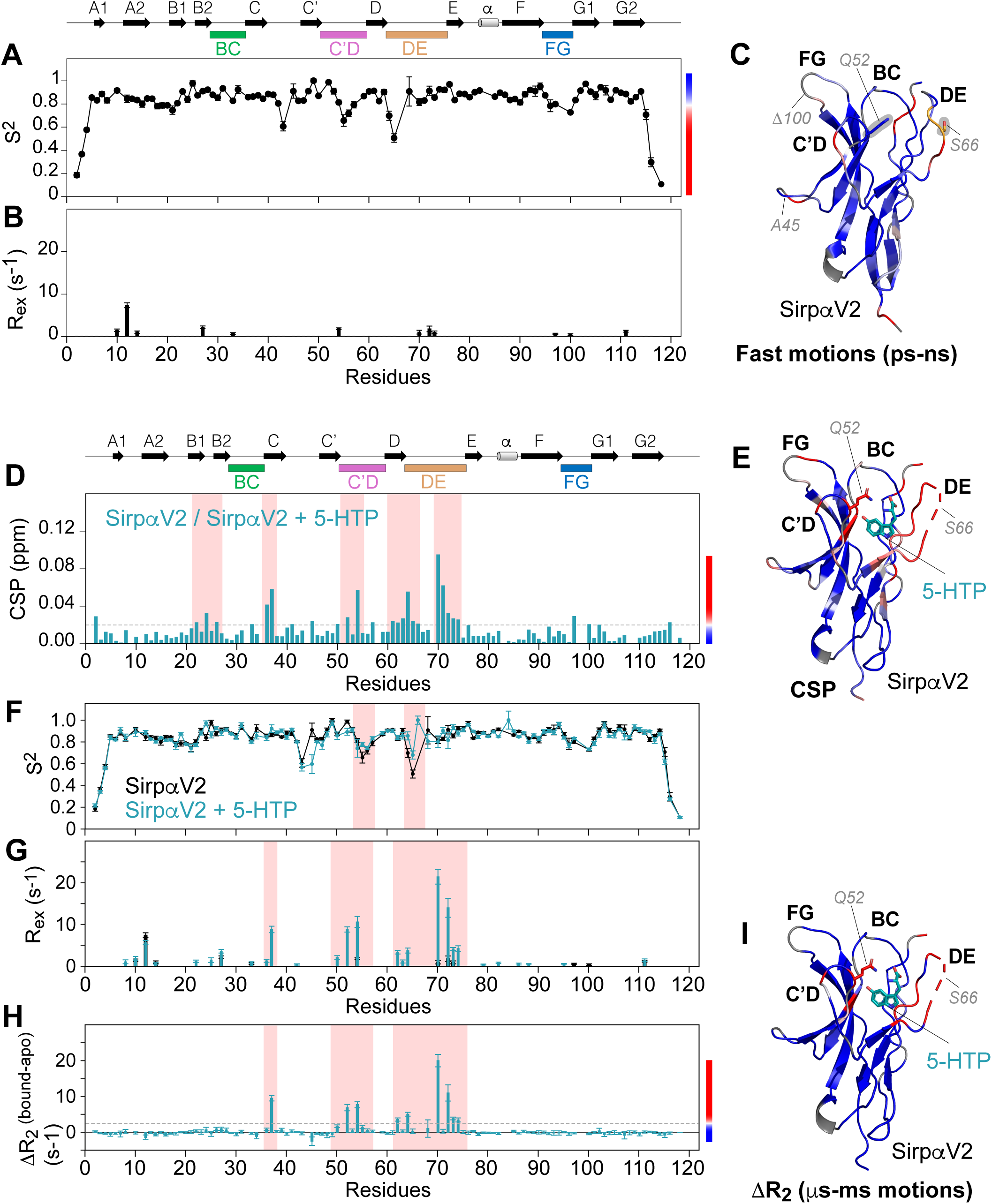
Experimental dynamics of SIRPαV2 and impact of 5-HTP ligand binding on SIRPαV2. (A, **B)** Backbone dynamics parameters of SIRPαV2, extracted from ^15^N relaxation data using the Lipari-Szabo model-free approach with an anisotropic global reorientation model: **(A)** order parameter S^2^ probing motions on the ps-ns timescale and **(B)** exchange contribution R_ex_ probing motions on the ms-ms timescale. Secondary structure elements and loops are indicated at the top. **(C)** Fast motions (S^2^) mapped on our SIRPαV2 structure (PDB code 9TF5), with color coding as indicated on panel **A**, i.e., from blue to red for increasing ps-ns dynamics (grey for missing data). Side chains of Gln52 and Ser66 are highlighted as sticks. **(D)** Chemical shift perturbations (CSP) on SIRPαV2 (200 µM) induced by a 5-fold molar excess of compound 5-HTP. Secondary structure elements and loops are indicated at the top. Regions around the WYF pocket are most affected by the binding (light red highlight). **(E)** Mapping of the CSP on our structure of SIRPαV2 bound to 5-HTP (PDB code 9T7F) with color ranging from blue to red for increasing CSP values as indicated in panel (**D**), and grey for missing data. Gln52 is shown as red sticks and 5-HTP as cyan sticks. **(F, G)** Backbone dynamics parameters (S^2^ and R_ex_) for SIRPαV2 (black) and 5-HTP bound SIRPαV2 (cyan). Regions most affected by ligand binding are highlighted in light red. **(H)** Difference of transversal relaxation rates between 5-HTP-bound and apo SIRPαV2 (DR_2_ = 1/T_2_^bound^ - 1/T_2_^apo^) as an indicator of increased conformational exchange in bound SIRPαV2. **(I)** Mapping of DR_2_ on the structure of SIRPαV2 bound to 5-HTP (PDB code 9T7F) with the color ranging from blue to red for increasing ms-ms dynamics in bound SIRPαV2 as indicated in panel (**H**), and grey for missing data. 5-HTP is shown as cyan sticks.

We used 5-HTP as a probe to study the effects of ligand binding on the dynamics of the WYF pocket in SIRPαV2 (**Figure 3D-I**; **Suppl. Fig. S8)**. Under the conditions of the relaxation studies (200 μM SIRPαV2 and 1000 μM 5-HTP) and given the measured K_D_ of ∼8 mM, we expected about 11% of SIRPαV2 bound to 5-HTP. Despite this modest occupancy, we observed small but significant CSPs in the region of the WYF pocket (**Figure 3D,E**). In the presence of the ligand, SIRPαV2 exhibited strong conformational exchange in the region of the WYF pocket (ΔR_2_ > 0 and R_ex_ > 0) as well as less flexibility of the C’D and DE loops (S^2^ > 0.7, **Figure 3F-I**), likely reflecting induced fit conformational changes necessary for 5-HTP to bind the WYF pocket and stabilization of this ligand-bound conformation.

To investigate the overall conformational landscape of SIRPα-D1 and the frequency of the WYF-pocket accessible conformation, we performed extensive 2-µs molecular dynamics simulations (MDS) in explicit water solvent starting from crystallographic structures representative of SIRPαV1 (PDB entry 6NMR, Chain I) and SIRPαV2 (PDB entry 2JJS, Chain A). Root mean square deviation (RMSD) analysis revealed that the SIRPαV2 variant exhibited slightly greater global flexibility (1.72 ± 0.41 Å) compared to SIRPαV1 (1.44 ± 0.47 Å), suggesting a potential increase in conformational plasticity in agreement with the NMR results above (**Suppl. Fig. S9**). Root mean square fluctuation (RMSF) analysis identified the DE (residues 64-71) and C’D (residues 52-57) loops, which define the entrance of the binding pocket, as highly flexible regions both in SIRPαV1 and SIRPαV2 (**Suppl. Fig. S10**). This was further supported by a clustering analysis that identified the 10 major conformations sampled during the MDS trajectories (**Suppl. Fig. S11**). To quantitatively characterize the conformational landscape of the binding pocket, we monitored two Cα–Cα interatomic distances: D1 (between residues Glu54 and Thr67), and D2 (between residues Gln52 and Phe74), which together capture the conformational fluctuation and spatial arrangement of the pocket entrance. Both trajectories sample a broad and continuous conformational space (**Suppl. Fig. S12**). Importantly, the ensemble of conformations observed in the simulations spans the full range of binding pocket geometries found in known x-ray crystallographic structures, suggesting good agreement between MDS trajectories and experimental data. Together, these findings suggest that both variants exhibit extensive and exhaustive sampling of the conformational space relevant to ligand binding.

### Gln52 is a Dynamic Gatekeeper Regulating WYF-Pocket Access

Our x-ray structure data and SIRPα conformational dynamics data from MDS and NMR suggested that Gln52 was a key residue modulating access to the WYF pocket. Located strategically at the pocket rim, the Gln52 side chain exhibited marked conformational variability, predominantly switching between two distinct rotameric states: the Gln52-in state wherein the Gln52 side chain hydrogen bonded to Ile36 main chain in an inward orientation that precluded access to the WYF pocket; and the Gln52-out state, wherein the Gln52 side chain hydrogen bonded to Tyr50 side-chain hydroxyl in an outward orientation that correlated with the WYF-open ligand-accessible state for SIRPαV2 (**Figure 2C, D**). The dynamic behavior of the Gln52 side chain as a key WYF-pocket gatekeeper is crucial for structure-based drug design efforts, and even more so in fragment-based drug design campaigns where initial hits typically display weak affinities.

Interatomic distances between Gln52-OE1 and Ile36-N, as well as between Gln52-OE1 and Tyr50-OH, were monitored throughout the MD simulations to identify potential hydrogen bonds (**Figure 4**). Analysis of the conformational ensemble revealed that Gln52 predominantly adopted the Gln-in orientation, occurring in 53% of the sampled conformations in SIRPαV1 and 41% in SIRPαV2. In contrast, the Gln-out conformation was observed much less frequently, representing only 5% and 8% of the conformations in SIRPαV1 and SIRPαV2, respectively. This distribution suggests a strong preference for the Gln-in state in both variants, with a slightly reduced prevalence in SIRPαV2.

**Figure 4.**
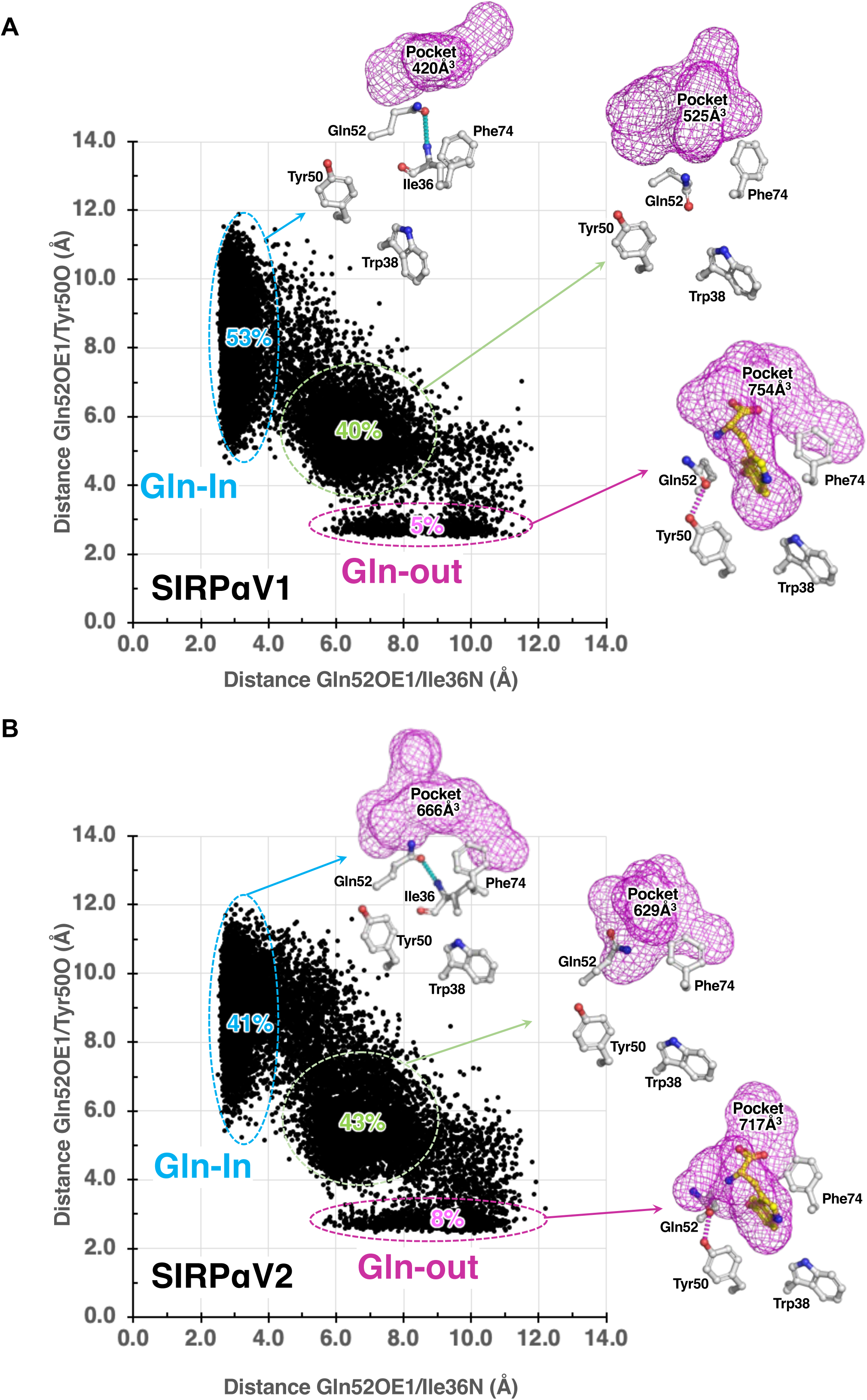
Molecular dynamics simulations reveal the conformational landscape of Gln52. Dynamics of Gln52 were analyzed using MD simulations for **(A)** SIRPαV1, and **(B)** SIRPαV2. To monitor hydrogen bond formation, interatomic distances were measured between Gln52-0E1 and Ile36-N (x-axis), and between Gln52-OE1 and Tyr50-OH (y-axis). These analyses revealed two extreme conformations, the relatively frequent “Gln-in” conformation where Gln52 forms a hydrogen bond with Ile36 and its side chain fills and closes the WYF pocket, and the rare “Gln-out” conformation, where Gln52 forms a hydrogen bond with Tyr50, leaving the WYF pocket open and able to accommodate a small molecule such as 5-HTP.

The frequency of binding pocket formation during MD trajectories was analyzed using MDPocket[23]. For SIRPαV1, 90% of the detected pocket volumes ranged from 220 to 640Å^3^, while the corresponding range was 210 to 644 Å^3^ for SIRPαV2 **(Suppl. Fig. S13)**. In both cases, these volumes corresponded to the presence of one to three subpockets. The binding pocket consisted of a large entrance pocket (up to 500 Å^3^) and the deeper WYF pocket (up to 300 Å^3^). The WYF pocket was occluded in most conformations due to the inward orientation of the Gln52 sidechain (Gln-in, **Figure 4**). The WYF pocket was observed when the Gln52 side chain pointed outward (Gln-out), in a conformation that could accommodate the 5-HTP moiety. In 30% of the frames, the WYF pocket was obstructed by Phe74, which adopted a distinct rotamer conformation involving a 90-degree rotation along the chi1 torsion angle (**Suppl. Fig. S14**). Interestingly, this conformation has never been reported in any x-ray structures.

### Gln52 Mutations Impact Thermal Stability and Protein Conformation

To investigate the role of Gln52 in modulating access to the WYF pocket, we performed a large multi-sequence alignment of SIRPα-D1 domains across species as well as in other SIRP family isoforms. To determine the conservation of key WYF pocket residues, we analyzed over 1410 sequences identified as SIRP in the Clustered-NR database. While the overall sequence identity of the D1 domain ranged from 47 to 99%, the WYF-pocket motif was strikingly well conserved (90%). We found that Trp38 was invariant, Tyr50 was mostly conserved (88%, or replaced by Phe, 10%), and Phe74 was mostly conserved (91% or replaced by Tyr 6%). We analyzed the conservation of the gatekeeper residue (position 52 in the human SIRPαV2 sequence). Gln and Phe were by far the most frequent (43 and 40%, respectively), while Leu, Asp, Pro (3-4%), and Ala, Arg, Lys (0.1-0.2%) were also represented (**Suppl. Fig. S15**). According to these results, we generated SIRPαV2 mutants predicted to bias SIRPαV2 towards the WYF-open (Q52A, Q52K, Q52R) or WYF-closed (Q52F) states. We predicted that mutants biased towards the WYF-open state would display enhanced flexibility, while the Q52F mutant would display increased stability.

We measured the overall stability of the SIRPαV1 and SIRPαV2 D1 domains, as well as the mutant SIRPαV2-D1 domains using nano differential scanning fluorimetry (nanoDSF, **Figure 5A**). In agreement with the dynamics data above, we found that SIRPαV1 was more stable than SIRPαV2 by ∼7°C (V1 Tm = 50.2°C, V2 Tm = 43.5°C). Compared to WT SIRPαV2, mutations predicted to favor the WYF-open state decreased protein stability: Q52A (Tm = 42.4°C), Q52K (Tm = 39.2°C), and Q52R (Tm = 39.2°C). In contrast, the Q52F mutation, predicted to favor the WYF-closed conformation, significantly enhanced thermal stability (Tm = 51.7°C).

**Figure 5.**
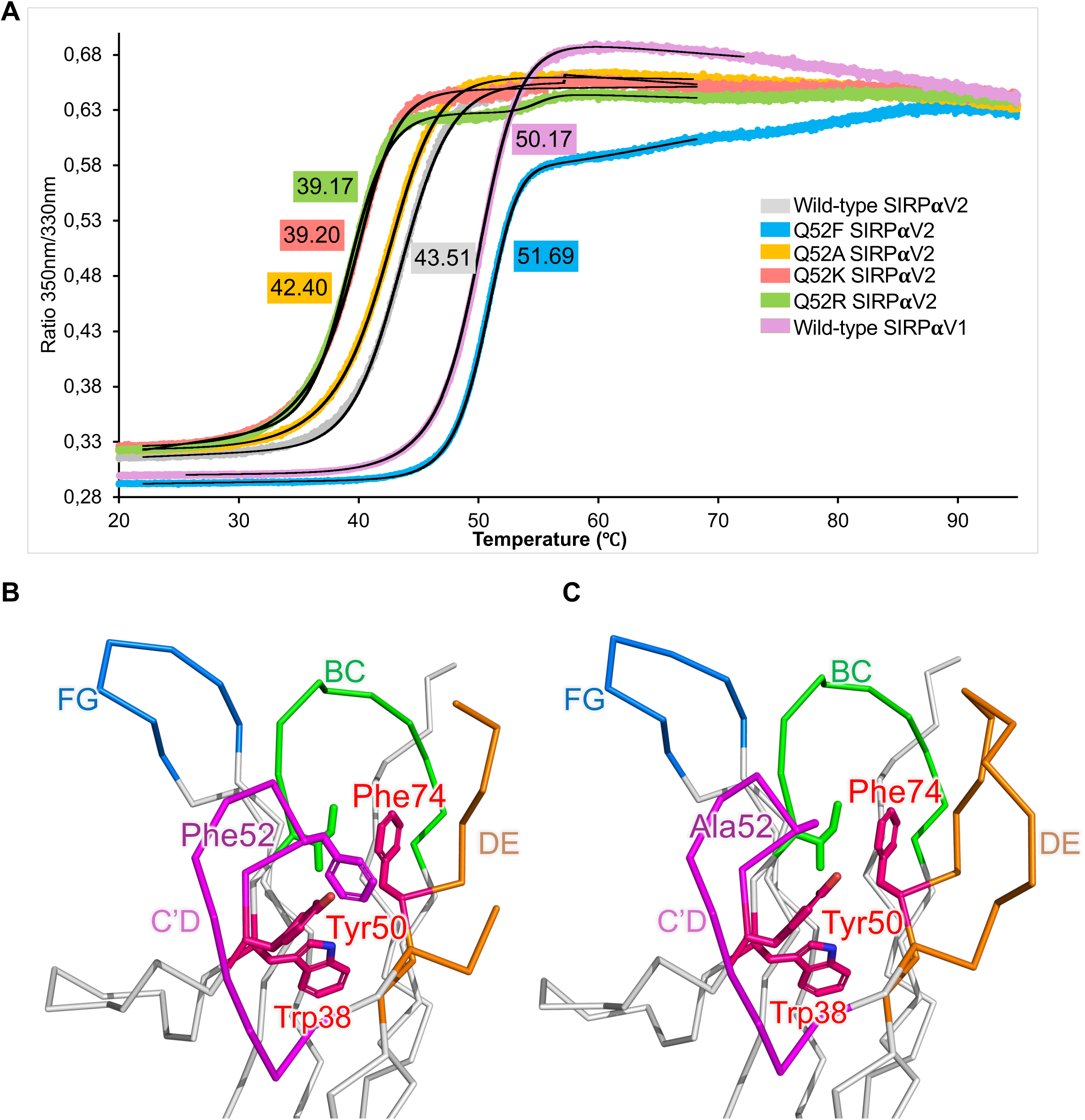
Gln52 acts as a conformational gatekeeper controlling SIRPα stability and accessibility. **(A)** Thermal stability of SIRPα variants measured by nanoDSF. The melting temperature (Tm) was determined by monitoring the intrinsic fluorescence of the proteins during a thermal ramp. Tm values for WT and mutant proteins are shown. Black curves represent the fit to experimental data (colored points). **(B)** Zoom on the WYF pocket showing the orientation of the C’D and DE loops and Q52F mutation (PDB entry 9SID) in the same orientation as in Fig. 2. (**C**) Zoom on the WYF pocket showing the orientation of the C’D and DE loops and Q52A mutation (PDB entry 9SIC) in the same orientation as Fig. 2.

To obtain direct structural evidence of the conformational effects of these mutations, we determined x-ray structures for the Q52A and Q52F SIRPαV2 mutants (**Figure 5B,C**). Crystals were obtained in similar conditions to WT SIRPαV2, and structures were solved at 1.5 and 1.7Å resolution, respectively (**Table S1**). In the Q52F SIRPαV2 mutant structure, the DE loop was poorly defined in the electron density (missing aa 65-68) in both molecules of the asymmetric unit. Phe52 was oriented inside the WYF pocket, stacked between Phe74 and Tyr50, which moved closer to Val60 and Phe57 to avoid a steric clash. Therefore, the bulky Phe52 side chain filled the hydrophobic cavity and formed stabilizing hydrophobic interactions that locked SIRPα in the WYF-closed conformation (**Figure 5B**). The structural and nanoDSF results confirmed our prediction regarding enhanced stability of this mutant, in agreement with previous studies on respiratory syncytial virus vaccine design[24], and recent computational methods that predict the effect of mutations[25].

In the Q52A mutant structure, we observed slightly different conformations of the C’D and DE loops in the three molecules of the asymmetric unit. DE loop residues 65-69 were disordered in two molecules. The distance between C’D loop residue 52 and DE loop residue 70 was 13.9 Å (chain A), 13.5 Å (chain B), and 15.1 Å (chain C). This last distance was comparable to the one observed in the x0408 fragment complex structure (16 Å), with an open WYF pocket. Therefore, this DE loop conformation, combined with the absence of the Gln sidechain, allowed opening of the WYF pocket in the absence of a ligand (**Figure 5C**). This result agrees with the decrease in stability observed by nanoDSF.

Efforts to solve the structure of the Q52R mutant, engineered to favor the WYF-open conformation and displaying an even lower Tm than the Q52A mutant, produced only fragile crystals. All collected datasets exhibited high anisotropy, possibly reflecting increased flexibility within the crystal lattice, and yielded poor diffraction data. Although the data did not allow us to refine a structure with acceptable statistics, we could still detect pronounced disorder in both the C’D and DE loops, supporting the high flexibility of this mutant and overall suggesting a preference for the WYF-open state.

### Gln52 Mutations Impact SIRPa Dynamics

To gain information on the dynamics of the WYF pocket in the SIRPαV2 Gln52 mutants, we performed NMR experiments on Q52F and Q52A SIRPαV2 proteins. Dynamics data were not recorded for Q52R SIRPαV2 as this protein was poorly expressed in ^15^N-enriched minimal media. Amide (^1^H, ^15^N) chemical shift assignment of the two mutants was performed by analogy to WT SIRPαV2 and careful analysis of neighboring residues in case of ambiguity (**Suppl. Fig. S16**). Phe52 induced pronounced amide CSPs within C’D and DE loops residues, the neighboring regions including the β-sheet spanning strands B1 and B2 (aa 24-28), and residues Ile36 and Trp38, which were located inside the WYF pocket (**Figure 6A,B**). These results confirmed that Phe52 significantly impacted the conformation of the pocket as observed in our crystal structure. Mutating Gln52 into an Ala resulted in much smaller perturbations, mostly located on neighboring Ala52 residues of the C’D loop (aa 50-55) and Ile36, whose main chain no longer hydrogen bonded to Gln52 (**Figure 6C,D**).

**Figure 6.**
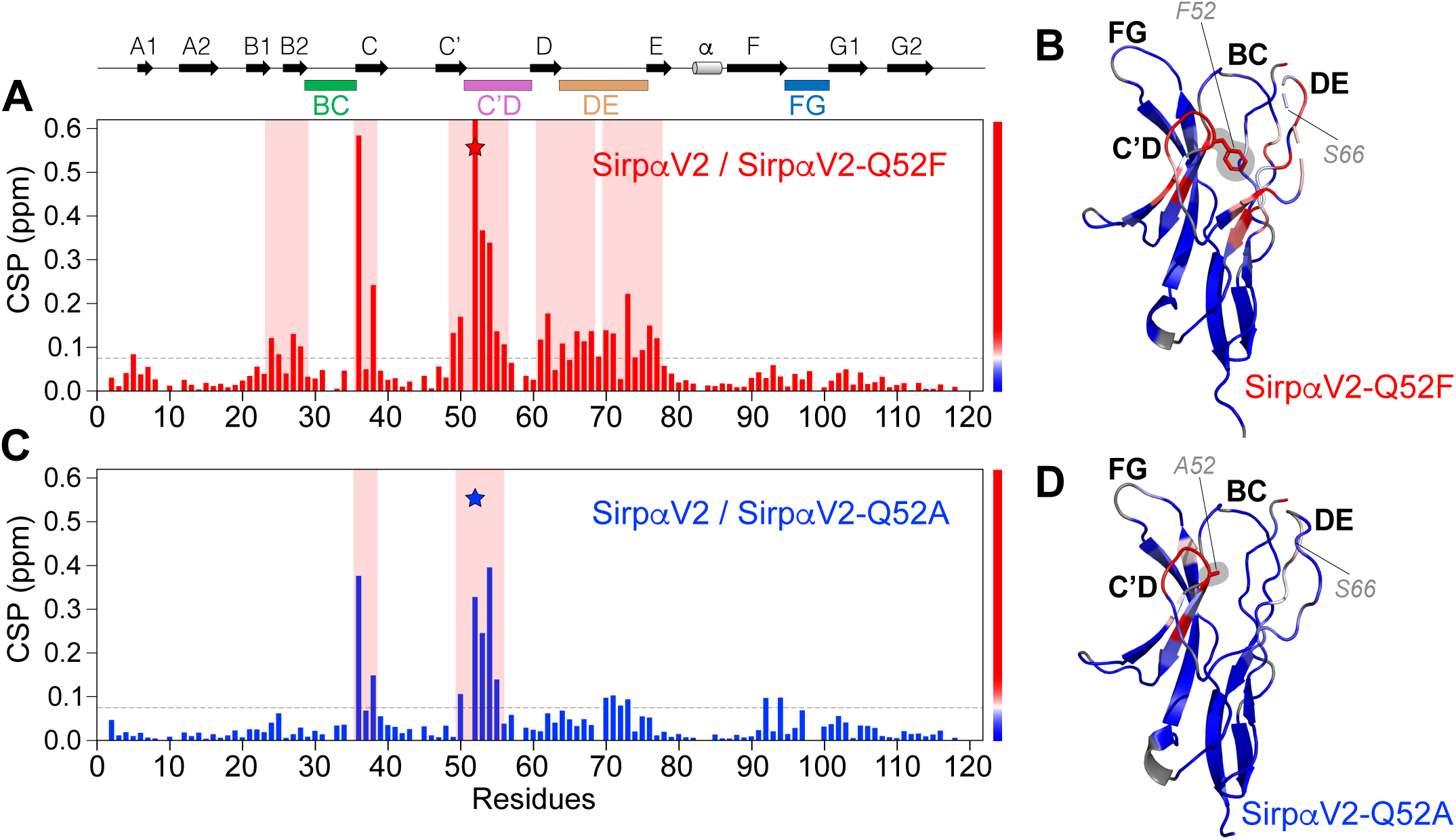
Gln52 mutations induce distinct conformational changes in the WYF pocket. (A,C) Chemical shift perturbations (CSP) in SIRPαV2 induced by the **(A)** Q52F and **(C)** Q52A mutations. Regions most affected by the respective Gln52 mutation (red and blue stars) are highlighted in light red. Secondary structure elements and loops are indicated at the top. **(B,D)** Mapping of the CSP on our structures of **(B)** SIRPαV2-Q52F (PDB entry 9SID) and **(D)** SIRPαV2-Q52A (PDB entry 9SIC), with color ranging from blue to red for increasing CSP values as indicated in panels (**A**,**C**), and grey for missing data. Amino acids PheF52 and Ala52 are shown as sticks.

To further assess pocket dynamics in both SIRPαV2 mutants, we recorded ^15^N relaxation data (**Suppl. Fig. S17**) and analyzed dynamics parameters (**Figure 7**). Compared to WT SIRPαV2, we observed an increase in S^2^ values for the Q52F mutant 62-66 region (ΔS^2^ ∼ +0.1 to +0.2), which suggested decreased flexibility of the DE loop (**Figure 7A,B**). In addition, we observed enhanced conformational exchange on the μs-ms timescale for C’D loop residues 54-57 (R_ex_ and ΔR_2_ ∼ 5-25 s^-1^), reflecting their flexibility at the expense of Phe52 anchoring in the WYF pocket (**Figure 7A,C,D**). In contrast, we observed no conformational exchange for Q52A SIRPαV2 (**Figure 7E,G**), but a modest decrease in S^2^ values (ΔS^2^ ∼ -0.1 to -0.2) in the C’D and DE loops, indicative of increased flexibility of the regions surrounding the entrance of the WYF pocket (**Figure 7E,F,H**).

**Figure 7.**
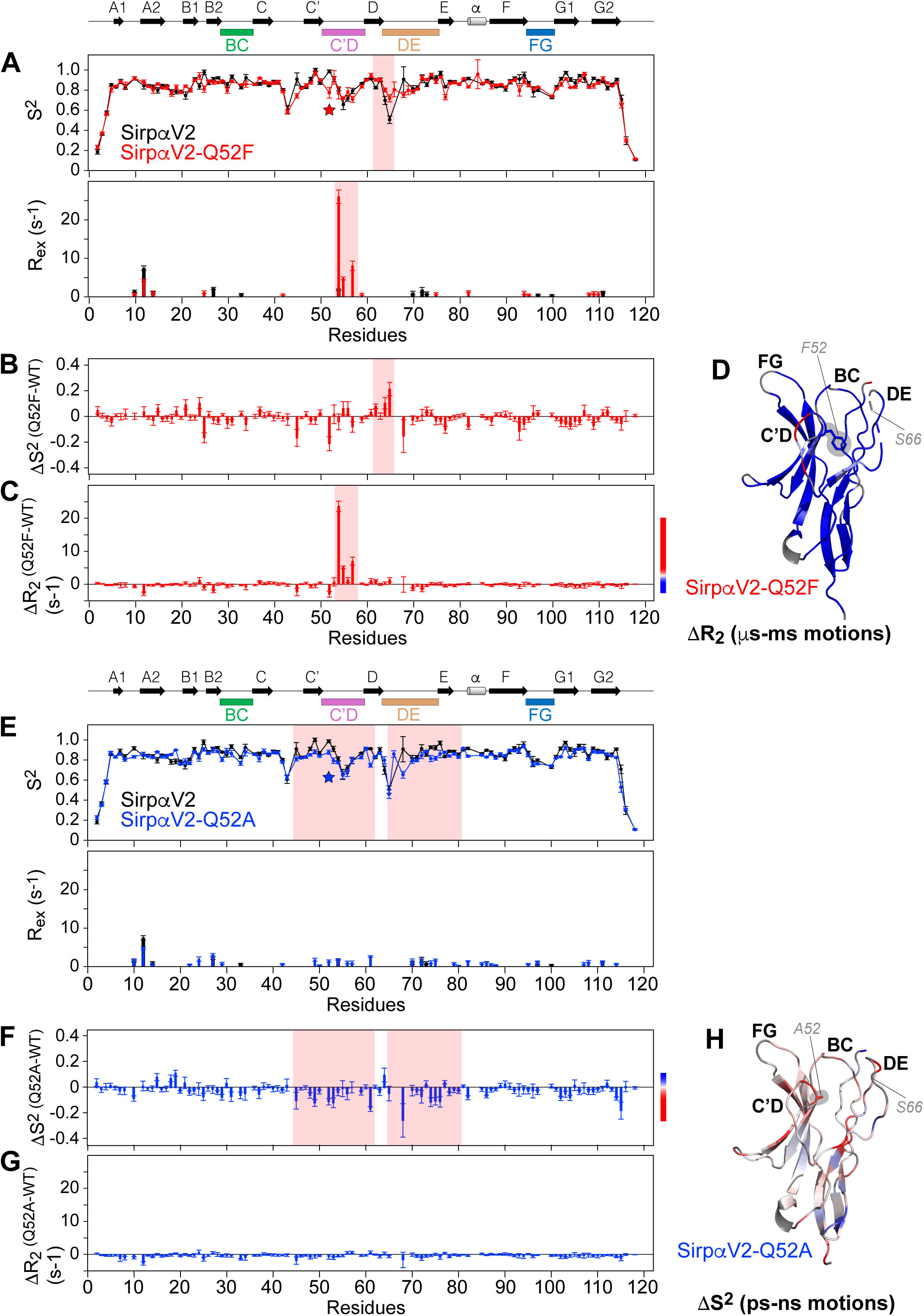
Gln52 mutations differentially impact the backbone dynamics of the WYF pocket. (A,E) Backbone dynamics parameters, S^2^ (top) and R_ex_ (bottom) of SIRPαV2 WT (black) and SIRPαV2-Q52F (red in panel A) and SIRPαV2-Q52A (blue in panel E). Secondary structure elements and loops are indicated at the top, and Q52F and Q52A mutations by the red and blue stars. **(B,F)** Difference of order parameters between the Gln52 mutants and WT SIRPαV2 (DS^2^ = S^2,mutant^ - S^2,WT^). **(C,G)** Difference of transversal relaxation rates between the Gln52 mutants and WT SIRPαV2 (DR_2_ = 1/T_2_^mutant^ - 1/T_2_^WT^) as indicator of conformational exchange. **(D)** Mapping of DR_2_ on our structure of SIRPαV2-Q52F (PDB entry 9SID), with color ranging from blue to red for increasing ms-ms dynamics in the Q52F mutant as indicated in panel (C), and grey for missing data. Phe52 is shown as sticks. **(H)** Mapping of DS^2^ on our structure of SIRPαV2-Q52A (PDB entry 9SIC), with color ranging from blue to red for increasing fast ps-ns dynamics in the Q52A mutant as indicated in panel (F), and grey for missing data. Ala52 is shown as sticks.

We observed a slight divergence between crystal structures and solution data regarding the Q52F mutant. In X-ray structures, the DE loop of Q52F appears disordered, whereas NMR relaxation data indicate the loop is more rigid compared to WT. This likely reflects the limitations of crystallography, where lack of electron density can result from static disorder or multiple discrete conformations, whereas NMR parameters directly probe ps-ns motions in solution. We conclude that the NMR data more accurately reflect the locked nature of the Q52F DE loop in a physiological environment.

In summary, stabilization of the DE loop and insertion of the Phe52 bulky side chain in the pocket of the Q52F mutant likely restricted access to open-like conformations of the WYF pocket, thereby potentially impairing ligand binding compared to WT SIRPαV2. In contrast, the enhanced dynamic environment around the pocket in Q52A SIRPαV2 may facilitate access to the binding site and improve ligand affinity compared to WT SIRPαV2.

To investigate the impact of Gln52 mutations, 50-ns MD simulations in explicit aqueous solvent were performed for WT, Q52A and Q52F SIRPαV2, and pocket analyses were performed with MDPocket as described above. For WT SIRPαV2, the conformational behavior and pocket properties were consistent with those observed in the longer 2-µs MD trajectory (**Suppl. Fig. S18**). As expected, removal of Gln sidechain (Q52A) resulted in a significantly larger average pocket volumes compared to WT (460.4 ± 123.9 Å^3^ vs 429.4 ± 141.7 Å^3^; p<0.0001 one-way ANOVA; **Suppl. Fig. S18 A,B**). In contrast, the Q52F mutant exhibited an average pocket size comparable to WT (425.4 ± 121.7 Å^3^). However, in this mutant, the WYF pocket remained occluded throughout the MD simulation due to the position of the Phe52 side chain inside the pocket, in agreement with the mutant crystal structure (**Suppl. Fig. S18C**). This position of Phe52 remained unchanged, contrary to Gln52 in WT SIRPαV2, where it sampled multiple side-chain conformations allowing for opening and closing of the WYF pocket.

### Gln52 Mutations Impact Binding to CD47

To determine the effect of the Gln52 mutations on SIRPαV2 interaction with CD47, we measured binding affinity and thermodynamic parameters by ITC (**Table 1, Suppl. Fig. 19A,B**). Under our experimental conditions, WT SIRPαV2 bound CD47 with a K_D_ of 0.43 ± 0.1 μM in agreement with previous reports[1,14,26–28]. The Q52F SIRPαV2 mutant showed no detectable binding to CD47. The Q52A and Q52R SIRPαV2 mutants showed significantly reduced affinity for CD47 compared to WT (∼13-fold and ∼6-fold increase in K_D_, respectively). The Q52K mutant displayed measurable but very weak binding to CD47 at the limit of ITC detection (80 ± 40 μM; ∼186-fold lower vs. WT).

These results suggested that while Q52A and Q52R mutations still permitted binding, Gln52 played an important role in optimizing the interaction interface for high-affinity CD47 interaction. These observations were confirmed using a competition binding assay based on our AlphaLISA and HTRF conditions to assess the relative ability of the SIRPαV2 Q52 mutants to bind CD47 and cross compete (**Table 1, Suppl. Fig. 19C-H**). Consistent with its K_D_, untagged WT SIRPαV2 self competed with its biotin-tagged version for CD47 binding with an IC_50_ of 0.7 and 1.7 μM (in AlphaLISA and HRTF assays, respectively). Likewise, Q52A and Q52R SIRPαV2 self competed with their biotin-tagged version displaying weaker IC_50_ values. As expected, WT SIRPαV2 was more active when competing with biotin-tagged Q52A (6.5-fold lower IC_50_; HTRF) and Q52R (4.4-fold lower IC_50_; HTRF). Q52 mutants were also less active when competing against the biotin-tagged WT (Q52A 14-fold lower IC_50_; HTRF and Q52R 16-fold lower IC_50_; HTRF).

In summary, Gln52 mutations in SIRPαV2 significantly impaired its binding to CD47. While some mutations (Q52A, Q52R) showed reduced affinity, others (Q52F) abolished binding entirely, suggesting a crucial role of Gln52 in optimal CD47 interaction.

### Gln52 Mutations Impact Small Molecule Binding to the WYF pocket

To characterize the role of Gln52 in limiting ligand access to the WYF pocket, we measured the binding affinity of Trp and 5-HTP to SIRPαV2 Gln52 mutants using ITC (**Table 1, Suppl. Fig. 6A,B**). As expected, no binding interaction was observed between Trp or 5-HTP and the Q52F mutant, in which Phe52 occupied the WYF pocket. Conversely, replacing Gln52 with residues favoring the WYF-open state (Ala, Arg, Lys) resulted in increased binding affinity of Trp and 5-HTP for the mutants compared to WT SIRPαV2 (∼up to 4.6-fold (5-HTP) and 2.7-fold (Trp) improvement for Q52R). We confirmed the greater affinity of 5-HTP for the WYF-open SIRPα mutants using a recently developed photo Chemically-Induced Dynamic Nuclear Polarization (photo CIDNP) binding assay. In line with ITC results, we showed that 5-HTP had the greatest affinity for Q52R SIRPαV2 mutant (K_D_ = 2.3 ± 0.7 mM) followed by Q52A (K_D_ estimated >10 mM), while binding of 5-HTP to WT SIRPαV2 was too weak to measure using this technique (**Table 1, Suppl. Fig. S20).**

These findings provided strong support for a key role of Gln52 in restricting ligand access to the WYF pocket in WT SIRPα and demonstrated that the WYF-closed conformation prevented ligand binding, while mutations promoting the WYF-open state facilitated ligand engagement with the cryptic pocket. Collectively, these results suggested that enhanced binding affinity of the small-molecule ligands to the WYF-open SIRPαV2 state facilitated by mutations, translated into increased functional inhibition of the SIRPα-CD47 interaction.

## DISCUSSION

The SIRPα-CD47 axis represents a critical immune checkpoint, and inhibiting this interaction holds significant therapeutic promise in oncology. While antibody-based approaches dominate current strategies, the potential for small molecule inhibitors remains largely unexplored. In this study, we combined mutagenesis, x-ray crystallography, NMR spectroscopy, molecular dynamics simulations, and biophysical assays to investigate the conformational dynamics of the SIRPα-D1 domain and identify novel targeting opportunities.

The canonical SIRPα−CD47 complex exhibits a relatively flat protein−protein interface area of approximately 1000Å^2^, which lacks deep pockets suitable for small-molecule binding (**Figure 1B**). Analysis of SIRPα structures alone or in complex (with CD47 or anti-SIRPα antibodies) showed a remarkably similar overall structure, except for conformational variability in the C’D (aa 52-55), the DE (aa 65-68), and the FG loops (aa 97-99) and no apparent ligand binding pocket (**Figure 1B**).

Our findings highlight the significant plasticity of the CD47 binding site, revealing a previously unknown ligand binding pocket (WYF pocket). Pocket formation is characterized by a dynamic equilibrium between a WYF-closed conformation, consistent with previously published structures, and a ligand-stabilized WYF-open conformation. Crucially, this open state exposes a cryptic, yet druggable, binding pocket that is only accessible in a conformation that is incompatible with CD47 binding. Central to this mechanism is the discovery that a single residue, Gln52, functions as a pivotal conformational gatekeeper, which dynamically controls the equilibrium between the WYF-closed state and the pocket-bearing WYF-open state. Accordingly, engineered mutations of this gatekeeper residue perturb the conformational equilibrium and provide strong validation for this model.

By identifying small-molecule fragments that bind to this cryptic site and functionally disrupt the SIRPα-CD47 interaction, this work establishes a validated structure-based roadmap for the development of a new class of conformationally selective inhibitors targeting this key immune checkpoint.

### Defining the Cryptic Nature of the WYF Pocket

The WYF pocket fits the classical definition of a cryptic site whereby a binding pocket that is absent or occluded in the unbound protein structure becomes apparent upon conformational changes, which can be induced or stabilized by a ligand[29]. Analysis of the available structural data for SIRPα confirms this classification. The pocket is not observed in previously solved structures of apo SIRPαV2 (PDB entry 2UV3) or in complex with its natural ligand CD47 (e.g., PDB entry 2JJS). In contrast, the pocket is present in the crystal structures described in this work, where it is occupied by various fragment hits like x0408, L-Tryptophan (Trp), and 5-hydroxy-tryptophan (5-HTP), as well as in its apo form in our MDS trajectory though at very low frequency. The discovery of this novel site was serendipitous, reflecting a recurring pattern in the identification of functionally relevant cryptic pockets, which are often revealed only when a ligand-bound structure is solved[30]. This finding underscores the utility of fragment-based methods not just for hit identification, but for druggable site discovery, effectively expanding the targetable landscape of proteins previously considered intractable[29]. Significantly, this pocket-bearing conformation is not unique to the fragment-bound SIRPαV2 variant. It is also observed in the structure of the SIRPαV1 variant in complex with an antagonistic antibody Fab fragment (PDB entry 6NMR), where the antibody HCD3 loop inserts a tryptophan residue into the exact same WYF pocket. The superposition of the fragment hits with this particular tryptophan shows a remarkable conservation of the key edge-to-face π-stacking interactions with Tyr50 and Phe74, validating the pharmacophoric features of this site across different SIRPα variants and binding modalities (**Figure 8A**). The opening of the WYF pocket requires substantial conformational rearrangement of the C’D and DE loops and the Gln52 side chain, yielding a conformation that is incompatible with CD47 binding, but allows ligand binding.

**Figure 8.**
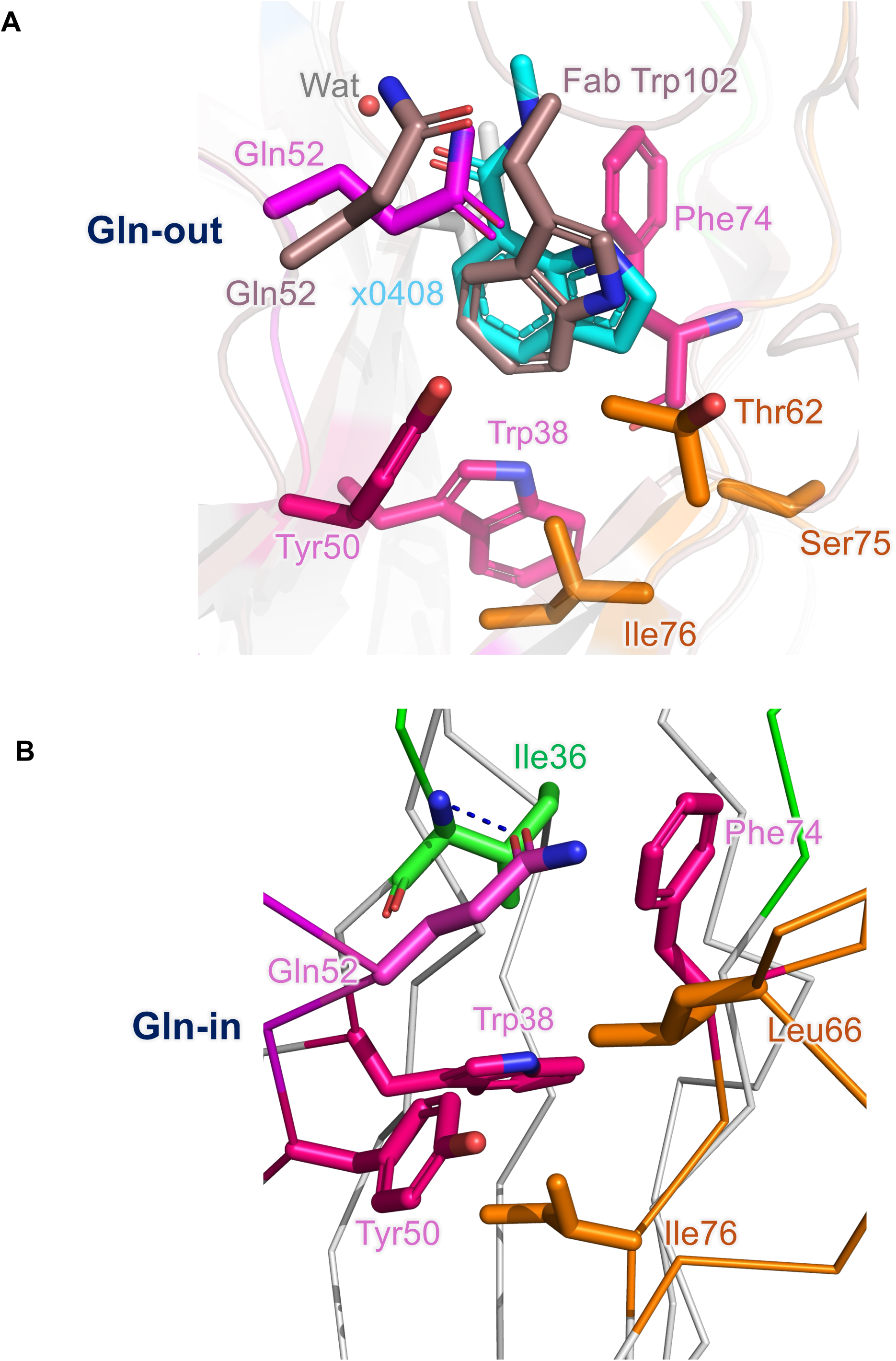
The cryptic nature of the WYF pocket. **(A)** The binding mode of fragment x0408 (cyan stick) in the WYF pocket of SIRPαV2 is similar to that of Fab Trp102 (brown stick) side chain bound to SIRPαV1 (PDB entry 6NMR). Both structures show the Gln52-out conformation (magenta and brown). (B) With Fab fragments that do not bind near the CD47 interface (PDB entry 6NMS), SIRPαV1 Gln52 is in the Gln-in conformation, and DE loop residues Leu66 stabilize a closed conformation that prohibits access to the WYF pocket.

The findings presented herein establish a new conceptual framework for targeting the SIRPα-CD47 checkpoint with small molecules. While the overall interaction surface between SIRPαV1 and the Fab fragment (PDB entry 6NMR, chains I and F, respectively) that induces the WYF-open conformation spans 779.0 Å^2^, we show that the WYF-open conformation can be induced and stabilized using fragment-sized ligands (TPSA 99.3 Å^2^) that are albeit much weaker but represent the size of a single amino acid. This allows us to propose a small-molecule inhibition strategy based on structure-based drug design using tools like ChemoDOTS[31] to elaborate the initial fragment hits into higher affinity lead molecules.

Rather than attempting the difficult task of designing an orthosteric competitor to block a large and flat PPI interface, this work validates a conformation-trapping inhibition strategy. The mechanism relies on stabilizing the intrinsic low-population WYF-open conformation of SIRPα that is inherently incompetent for CD47 binding. This presents a diversified approach for drug discovery, as conformation-trapping modulators can offer greater selectivity and novel pharmacological profiles compared to traditional orthosteric inhibitors[29]. While the identification and validation of such cryptic sites often remain a major challenge, this study provides a successful blueprint for achieving this goal on a therapeutically relevant PPI target.

### Protein Dynamics as the Origin of the Cryptic Pocket

A critical distinction in cryptic site discovery is whether a pocket is purely ligand-induced or if it represents a pre-existing, transiently sampled state[32]. Our comprehensive dynamics data strongly support the latter. NMR relaxation measurements reveal significant motion on the ps-to-ns timescale in the C’D and DE loops that form the WYF pocket lid. This flexibility is a prerequisite for large-scale movements required to open and close the pocket. Furthermore, the all-atom molecular dynamics simulations (MDS), initiated from both the closed and open (Fab fragment-bound) states, demonstrate that even in the absence of a ligand, the protein spontaneously samples conformations where the pocket is accessible, albeit with low frequency. This suggests a mechanism that likely involves both conformational selection and induced fit; the intrinsic flexibility of SIRPα allows it to transiently breathe and expose the pocket (conformational selection), which can then be captured and stabilized by a ligand (induced fit)[32].

This dynamic view also provides a rationale for the initial focus on the SIRPαV2 variant. While this choice was first made for practical reasons related to crystallization, subsequent NMR dynamics data revealed it to be advantageous. We show that the SIRPαV2 DE loop (around residue Ser66) is significantly more flexible than the corresponding SIRPαV1 loop. This is in agreement with structures of SIRPαV1 in complex with antibody Fab fragments that do not bind near the CD47 interface (PDB entries 6NMS, 6NMU, 6NMV, 6NMT) and that can be used as surrogates for apo SIRPαV1 structures. In these structures, DE loop residue Leu66 and C’D loop residue Gln52 both face the protein core and stabilize a closed conformation of the WYF pocket (**Figure 8B**). In contrast, our NMR and MDS data reveal a different configuration in SIRPαV2, whereby the WYF pocket entrance is closed by Gln52 but the pocket remains more accessible due to the high flexibility of the DE loop. This also likely manifests in the lower affinity of Trp and 5-HTP ligand binding to SIRPαV1 compared to SIRPαV2 in our SPR and HSQC study. As the DE loop acts as the primary lid covering the WYF pocket, its enhanced flexibility in SIRPαV2 lowers the energetic barrier for pocket opening and makes this variant an inherently more receptive target for a cryptic-site-directed drug discovery campaign. Gln52 can also hydrogen bond to DE loop residue S66 (as observed in PDB entry 2UV3) and limit pocket accessibility. Removal of the side chain in the Q52A mutant could provide greater accessibility of the WYF pocket, though we did not observe additional flexibility of the DE loop in our NMR dynamics studies.

### Redefining the Gatekeeper Concept for a Dynamic System

The combined mutagenesis, structural, and dynamic data presented in this study identify Gln52 as the key modulator of the WYF pocket conformational state. The term gatekeeper has been widely adopted in structural biology, primarily from studies of protein kinases. In that context, the gatekeeper is typically a single residue (often a threonine) in the ATP-binding pocket whose side chain sterically controls access to an adjacent hydrophobic pocket, thereby determining sensitivity to a wide range of small-molecule inhibitors. Mutations that increase the size of the gatekeeper (e.g., the T315I mutation in Abl kinase) confer drug resistance by physically blocking the inhibitor from binding[33]. More recently, structural studies highlighted the key role of third-shell isozyme-specific residues allowing conformational changes of a flexible conserved Gln gatekeeper residue to open a cryptic ligand pocket in the Nitric Oxide Synthase enzyme[34].

We propose that SIRPα Gln52 represents a distinct and more dynamic type of gatekeeper, which does not simply function as a steric block. Instead, it acts as a conformational switch, mediated by the rotameric state of its flexible side chain (**Suppl. Fig. S21**). In the Gln52-in state, its side chain points inward, forms a hydrogen bond with Ile36 main chain and effectively seals the pocket. In the Gln52-out state, the side chain swings outward to hydrogen bond with the Tyr50 hydroxyl group, a movement that is coupled to the large-scale opening of the entire DE loop pocket lid. Therefore, Gln52 is not merely a gatekeeper of inhibitor shape complementarity; it is a gatekeeper of conformational accessibility that orchestrates a concerted structural rearrangement to toggle the protein between its closed and open functional states.

We propose a mechanism linking the Gln52 rotameric state, WYF pocket conformation, protein stability, and ligand binding (**Figure 9**). We tested and confirmed this hypothesis with rationally designed mutations at position 52 to favor the WYF-open or WYF-closed pocket conformation.

**Figure 9.**
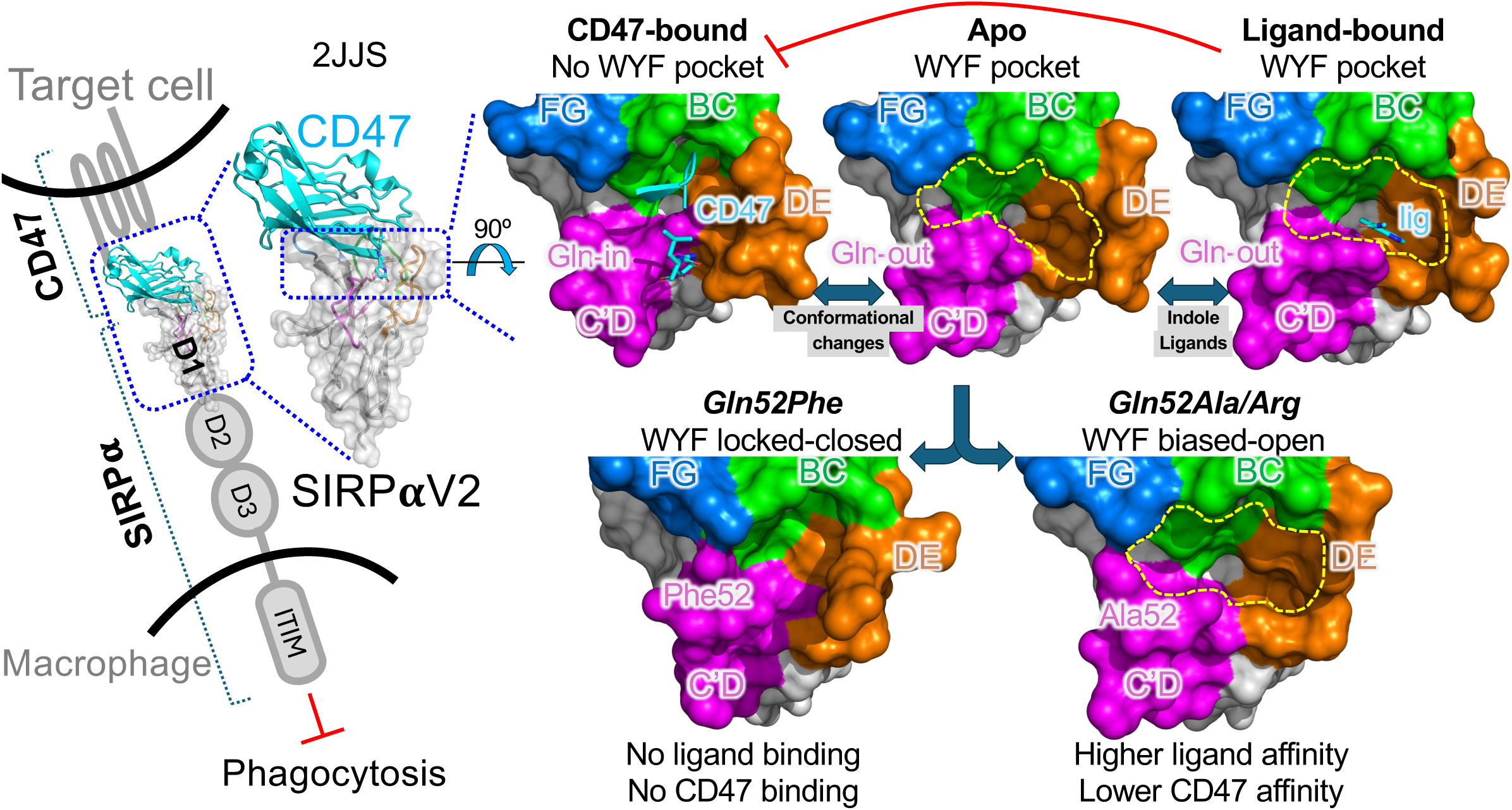
Proposed model for SIRPα conformational selection. Conformational plasticity of the CD47 binding domain of SIRPα reveals a cryptic WYF pocket gated by Gln52 and that binds indole ligands. Gln52 mutants favor specific conformations and modulate CD47 and ligand binding.

### The ‘Locked-Closed’ State

The Q52F mutant was designed to mimic a permanently closed state by introducing a bulky hydrophobic phenylalanine residue predicted to favor an inward-facing orientation of the C’D loop. We confirmed this prediction with combined biophysical, structural, and dynamics studies. First, the Q52F mutant exhibits a significant increase in thermal stability, with a melting temperature (Tm) approximately 8°C higher than the WT protein, indicating a more rigid and stable fold. Second, the crystal structure of Q52F SIRPαV2 reveals that the Phe52 side chain is indeed oriented into the WYF pocket, where it stacks between Phe74 and Tyr50, physically occupying and occluding the site. Third, NMR data show that the DE loop is less flexible in this mutant, consistent with it being locked down. Functionally, this locked-closed conformation abolishes detectable binding to the natural ligand CD47 while also preventing ligand binding to the cryptic pocket. Since the native CD47-bound state is also somewhat closed, this result highlights that closed is not synonymous with binding-competent. The loss of affinity for CD47 in Q52F is likely twofold: first, the Q52F mutation eliminates the hydrogen bond interaction to CD47 Glu104 amide. Second, the bulky phenylalanine side chain, while filling the hydrophobic WYF pocket, induces subtle distortions in the C’D loop that prevent the precise surface complementarity required for CD47 engagement.

### The ‘Biased-Open’ State

Conversely, the Q52A, Q52R, and Q52K mutants were designed to favor the open state by removing the key intramolecular interactions mediated by the Gln52 side chain. Our data show that these mutants are destabilized compared to WT SIRPα, with Tm values decreasing by 1 to 4°C, consistent with disruption of favorable native contacts. The x-ray structure of the Q52A mutant reveals a more open WYF pocket conformation even in the absence of a bound ligand, as confirmed by NMR data. The functional consequences of this biased-open state reveal the dichotomy that lies at the heart of this study therapeutic premise. While biasing the conformation toward the open state, these mutations have opposing effects on the two classes of ligands. On one hand, the affinity for CD47 is dramatically reduced in these mutants, with the dissociation constant (K_D_) increasing by 8-to 180-fold compared to WT SIRPα. This demonstrates that the high-affinity interaction with the natural ligand requires a Gln-in-like conformation. Both Q52R and Q52K were designed to favor the open state by eliminating the stabilizing Gln-in hydrogen bond to Ile36. However, while Q52R retains some CD47 binding, Q52K interaction is nearly undetectable. Since both residues are positively charged and of similar size, this suggests that the local electrostatic environment or specific packing interactions are critical for maintaining a residual binding-competent conformation. This discrepancy underscores that the open state is likely an ensemble of conformations rather than a single rigid geometry.

On the other hand, the affinity for the cryptic pocket ligand 5-HTP is improved. The K_D_ for 5-HTP binding to the Q52A, Q52R and Q52K mutants decreases by 1.9-to 4.6-fold, respectively. These results confirm that the WYF pocket is more accessible in the open state and that stabilizing this conformation facilitates ligand engagement.

Our integrative studies reveal the role of Gln52 as a conformational gatekeeper. The inverse correlation between CD47 affinity and cryptic pocket ligand affinity, mapped to the conformational state controlled by Gln52, provides a solid mechanistic foundation for a novel conformation-trapping inhibitor strategy.

### Engineering SIRPα Flexibility to Alter its Function

The insights into SIRPα conformational control gain further significance when contrasted with previous protein engineering efforts on this target. This comparison reveals the novel “flexibility-for-inhibition versus rigidity-for-activation” biophysical paradigm that governs SIRPα function.

We show that increasing conformational dynamics of the SIRPα binding site, as seen for the Q52A and Q52R mutants, disrupts the canonical high-affinity interaction with CD47 while simultaneously unmasking a druggable allosteric pocket. The therapeutic goal is therefore to develop a small molecule that mimics this effect, effectively acting as a molecular wedge to favor the flexible non-binding open state. Directed evolution has been used previously to create high-affinity SIRPα variants (e.g., CV1, FD6) acting as CD47-blocking biologics[35]. These engineered proteins exhibit up to 50,000-fold increase in affinity for CD47. Structural analysis of these variants revealed that the affinity enhancement is achieved through a series of mutations that stabilize and rigidify binding loops, particularly the C’D loop. As a result, the entropic penalty of binding is minimized by pre-organizing the binding interface into an optimal high-affinity conformation for engaging CD47.

Synthesizing these two opposing strategies reveals that SIRPα function is a direct readout of the conformational state and dynamics of its D1 domain, which can be tuned in opposite directions for different therapeutic outcomes. To create a potent SIRPα-based antagonist of CD47 (a super-agonist SIRPα), one must engineer rigidity into the interface. In contrast, we show that exploiting SIRPα intrinsic flexibility allows for the discovery of small-molecule inhibitors that bind SIRPα and prevent CD47 interaction by favoring the opening of the WYF pocket.

### Evolutionary and Biological Context for SIRPα Plasticity

The question then arises: why does SIRPα possess this inherent conformational plasticity? The evolutionary pressure on immune receptors provides a possible explanation. The paired receptor theory, proposed by Barclay and Hatherley, suggests that inhibitory receptors like SIRPα, which recognize self-ligands to maintain homeostasis, are prime targets for pathogens seeking to co-opt these pathways to suppress the host immune response[36]. To counteract this, the host evolves activating receptor counterparts (e.g., SIRPβ) with highly similar extracellular domains but opposing signaling functions. This sets up an evolutionary arms race that drives rapid evolution, polymorphism, and conformational flexibility in the receptor families as the host attempts to evade pathogen binding while maintaining self-recognition[36]. The intrinsic flexibility of the SIRPα-D1 domain, the observed differences between the V1 and V2 allelic variants, and the existence of the cryptic WYF pocket can be rationalized as potential consequences of this continuous evolutionary pressure. The cryptic pocket may be an “evolutionary spandrel”, that is a non-selected structural feature that arose as a byproduct of the selection for overall conformational plasticity.

In addition, the biological context of SIRPα is more complex than its interaction with CD47 alone. It is known to bind other ligands, such as the collectin Surfactant Protein D (SP-D), a soluble pattern recognition molecule involved in pulmonary innate immunity[37]. Notably, SP-D binds to the membrane-proximal D3 domain of SIRPα, a site completely distinct from the D1 domain that binds CD47 and contains the WYF pocket. This establishes SIRPα as a multi-functional signaling hub. The conformational dynamics observed in the D1 domain may therefore have unappreciated roles in allosterically modulating signaling from the D3 domain or in integrating signals from multiple simultaneously bound ligands.

### Conservation of WYF-Pocket and Gln gatekeeper motif in SIRP proteins across species

The SIRP family of proteins, including SIRPα, is considered to have appeared around the same time as the emergence of the adaptive immune system in vertebrates, approximately 500 million years ago. While pinpointing the single most primitive species is challenging, the evolutionary evidence points to the presence of SIRPα and its related family members in early vertebrates including mammals, birds and fish[38–41]. The SIRP family, and by extension SIRPα, is thought to be evolutionarily related to a primordial antigen receptor that predates the development of the rearranging antigen receptors (like T-cell and B-cell receptors) that are characteristic of the adaptive immune system[28,42]. The structural similarities between SIRPα immunoglobulin domains and those involved in vertebrate antigen recognition support this evolutionary link. The high degree of conservation in the SIRPα-CD47 binding interface underscores the importance of this regulatory mechanism. While human CD47 does not effectively interact with mouse SIRPα (aside from the NOD mouse strain[43]), highlighting some species-specificity, the fundamental mechanism is conserved. The structural conservation of the extracellular domains ensures the proper presentation of the CD47 binding site and the overall architecture necessary for cell-cell communication. The juxtaposition of the highly conserved WYF structural motif (90% conservation) with the evolutionarily divergent gatekeeper residue (Gln, Phe, etc.) suggests a finely tuned functional interplay. We propose that while the WYF aromatic core is selectively conserved to maintain the IgV domain structural integrity, the gatekeeper position functions as an evolutionary rheostat. By varying the residue at position 52 (e.g., Gln vs. Phe), evolution modulates the conformational dynamics of the binding interface, thus balancing the rigidity required for high-affinity CD47 recognition against the plasticity needed to evade pathogen engagement. Consequently, the cryptic pocket identified here is likely an intrinsic structural feature of this dynamic trade-off and presents an attractive vulnerability for therapeutic targeting.

### Conclusions and Roadmap for Inhibitor Development

Our comprehensive multi-layered investigation into the conformational dynamics of the SIRPα D1 domain leads to three major advances. First, it reports the discovery and validation of the novel druggable WYF cryptic pocket that partially overlaps with the CD47 binding site. Second, it identifies and rigorously characterizes Gln52 as a dynamic, conformational gatekeeper whose rotameric state allosterically controls the equilibrium between a closed state and an open, pocket-bearing, non-binding state. The CD47-binding competent state lies in between these two extreme conformations. Third, by demonstrating that small-molecule fragments can bind this pocket and functionally inhibit the SIRPα-CD47 interaction, it establishes and paves the way for a new therapeutic strategy based on the allosteric stabilization of a CD47 non-binding protein conformation.

This detailed mechanistic understanding provides a clear roadmap for a rational structure-guided drug discovery program: the immediate next steps will involve chemical optimization of the identified indole-based fragments to develop lead compounds with nanomolar potency and favorable pharmacological properties. Perhaps most powerfully, the work described herein has generated not only a hypothesis but also tools needed to accelerate its translation. Screening for ligands that bind to transiently populated cryptic pockets is inherently inefficient. The engineered open-biased Q52A and Q52R mutants, which present a stabilized and more accessible version of the WYF pocket, can now be employed as superior screening reagents in future HTS or biophysical screening campaigns. By lowering the energetic and kinetic barriers for hit identification, these mutants will dramatically accelerate the discovery of new, diverse, and more potent ligands targeting this allosteric site. This transforms the conclusion of the present study into the starting point for the next phase of inhibitor development. Ultimately, this mechanistic dissection of SIRPα dynamics has unlocked a new and highly promising avenue for the development of a first-in-class small-molecule immunotherapy for cancer.

## MATERIALS AND METHODS

### SIRPα and CD47 protein expression and purification

The CD47-CD4-6His construct encompasses the extracellular domain (Gln19-Pro139) of human CD47 (Uniprot Q08722-1) fused to the rat CD4 domains 3 and 4, and a C-terminal 6xHis tag. It was expressed and purified as described previously[14]. The SIRPαV1-biotin and SIRPαV2-biotin proteins were expressed and purified as described previously[14]. The codon-optimized shorter WT SIRPαV2 construct (residues 31-119, Uniprot P78324) with an N-terminal 6xHis-Thioredoxin-SUMO tag was synthesized by Biobasic, Inc, and inserted into the multi cloning site 1 (MCS1) of the expression vector pCDF-Duet between the NcoI and EcoRI restriction sites. The protein was expressed and purified similarly to the longer SIRPαV2-Avitag construct as described previously[14], without the final biotinylation reaction. The Gln52 mutant constructs were created using the Quick-Change mutagenesis kit (Agilent, Q52F and Q52A; **Suppl. Table S3**) or synthesized as gene fragments (Q52K and Q52R, GeneCust) and expressed and purified like the WT protein.

### X-ray crystallography

#### 1. Crystallization

Crystal trials for WT SIRPαV2-D1 were setup with an ArtRobbins Phenix robot with sitting drops containing 0.1 µL of protein (36 mg/mL) and 0.1 µL reservoir solution at room temperature. Large parallelepiped crystals were obtained in 2 days with 0.01 M zinc sulfate heptahydrate, 0.1M MES monohydrate pH 6.5, and 25% (v/v) PEGMME 550. Crystals were cryoprotected in 20% (v/v) ethylene glycol, flash-cooled in liquid nitrogen, and stored until data collection. Crystals of SIRPαV2 Q52A, Q52F, and Q52R were obtained manually in the same conditions.

#### 2. Soaking and crystal harvesting for fragment screening

The DSI-poised fragment collection was screened as described[17,44,45]. Crystals were soaked with 20 mM of each fragment (20% DMSO) for 3 hours at RT. The iSCB fragment collection was screened by soaking compounds at 20 mM (10% DMSO) for 18h-72h at 19°C.

#### 3. X-ray data collection

Stanford Synchrotron Radiation Laboratory (SSRL, Palo Alto CA, USA): x-ray data for WT SIRPαV2 were collected on beamline 9-2 at a wavelength of 0.98 Å, 100K, with a crystal-detector distance of 350 mm, oscillation range of 0.15°, and exposure of 0.2 sec. X-ray data were processed with HKL3000[46] and reduced with CCP4[47]. The structure was solved by molecular replacement with Phaser[48] using one SIRPαV2 monomer (PDB entry 2UV3) as a search model to find the three molecules in the asymmetric unit. Refinement was performed with Phenix[49]. Anomalous difference maps were calculated with Phenix. Data collection and refinement statistics are provided in **Table S1**.

Diamond Light Source XChem: x-ray data were collected at the beamline I04-1 at 100 K and processed with the fully automated pipelines at Diamond[50–52], which variously combine XDS[53], xia2[51], autoPROC and DIALS[51], and select resolution limits algorithmically; no manual curation of processing parameters was applied. Further analysis was performed through XChemExplorer[54]: for each dataset, the version of processed data was selected by the default XChemExplorer score, and electron density maps were generated with Dimple[55]. Ligand-binding events were identified using PanDDA[20], and ligands were modelled into PanDDA-calculated event maps using Coot[56]. Refinement was performed with Phenix[49].

Soleil/ESRF: x-ray data for WT SIRPαV2 with Trp and 5-HTP ligands and for the Q52A/F/R mutants were collected on beamlines Proxima1 and Proxima2a at the Synchrotron Soleil, Gif-sur-Yvette and beamline MASSIF at the European Synchrotron Radiation Source, Grenoble. X-ray data were processed with XDS[53]. The structures were solved by molecular replacement with Phaser[48] or Molrep[47] using WT SIRPαV2 as search model. Refinement was performed with Phenix[49] or Refmac[47]. Data collection and refinement statistics are provided in **Table S1**.

### MD simulations

#### 1. System Preparation for Molecular Dynamics Simulations

Initial structure coordinates were taken from the PDB: PDB entry 6NMR (chain I) for SIRPαV1 and PDB entry 2JJS (chain A) for SIRPαV2. Structural preparation prior to molecular dynamics was performed using the Molecular Operating Environment (MOE, Chemical Computing Group), where all non-essential solvent molecules (water molecules and ions) were removed. The target protein was embedded in an explicit solvent environment comprising a cubic water box with a physiological ionic strength of 0.1 M NaCl using the CHARMM-GUI web server (https://www.charmm-gui.org)[57–59]. The N- and C-termini were capped, and a disulfide bridge between Cys25 and Cys91 was explicitly defined.

#### 2. Molecular Dynamics Simulation Protocol

All-atom molecular dynamics (MD) simulations were carried out in explicit solvent with periodic conditions using the AMBER package version 2022[60,61]. Input files for MD simulations were generated using CHARMM-GUI[57–59], and the simulations were executed in three sequential phases: energy minimization, equilibration, and production. Hydrogen mass repartitioning (HMR) was applied to allow for an increased integration time step of 4 fs[62]. All simulations were executed on GPU-accelerated hardware using the CUDA-enabled PMEMD module of AMBER.

#### 3. Energy Minimization

The energy of the system was first minimized using 2500 steps of steepest descent to eliminate large steric clashes, followed by 2500 steps of conjugate gradient minimization to further relax the system. A harmonic positional restraint of 10 kcal/mol/Å² was applied to all heavy atoms during this step to maintain structural integrity while relieving steric clashes and unfavorable contacts. The simulation used a nonbonded cutoff of 12.0 Å and a force-based switching function starting at 10.0 Å. Periodic boundary conditions and long-range electrostatics were handled using particle mesh Ewald (PME). This step ensured proper relaxation of the solvent environment and minimized unfavorable contacts before proceeding to the equilibration stages.

#### 4. Equilibration

Equilibration was performed in three successive stages (250 ps each, totaling 750 ps) under NPT conditions. Each stage employed Langevin dynamics for temperature control and a Monte Carlo barostat for isotropic pressure control at 1.0 bar. The target temperature was set to 303.15 K. To progressively relax the system, a positional harmonic restraint was applied to heavy atoms during each stage with decreasing force constants of 1.0 kcal/mol/Å², 0.3 kcal/mol/Å², 0.0 kcal/mol/Å² (i.e., unrestrained). A 1-fs integration time step was used throughout equilibration. Nonbonded interactions were truncated at 12.0 Å with a force-based switching function applied from 10.0 Å. Long-range electrostatic interactions were treated using PME summation. Atomic coordinates were wrapped to the unit cell and saved periodically saved every 500 steps.

#### 5. Production

Following equilibration, production MD was carried out for a total of 2 µs using a 4-fs integration time step enabled by HMR and SHAKE constraints on all bonds involving hydrogen[62]. Simulations were performed in the NPT ensemble with the same temperature and pressure parameters as the equilibration phase. Positional harmonic constraints were removed for this phase. System coordinates were saved every 1000 steps. All simulations were conducted using the CUDA-enabled PMEMD module of AMBER on an in-house high-performance computing cluster. Each trajectory was run in parallel on a 24-core compute node equipped with GPUs, ensuring efficient performance for long timescale simulations.

#### 6. MD Analyses

MD trajectories were analyzed using a combination of in-house python and tcl scripts and tools from the AMBERTools22 package. Key structural and dynamical properties, including root mean square fluctuation (RMSF), root mean square deviation (RMSD), interatomic distances, and dihedral angles variations, were calculated using the CPPTRAJ module from Amber22 package.

#### 7. Pocket detection with FPocket and MDpocket

MDpocket was used to detect and analyze pockets along the MD trajectories[23]. A first run was performed using the default parameters to identify all pockets present at the surface of SIRPα. From this analysis, a PDB file containing the grid point spheres defining the binding site pocket of interest was generated. A second MDpocket run was then performed, restricting the search to this predefined pocket. To further characterize the binding site pockets, Fpocket was used to identify pocket on individual frames and to estimate the volume of each subpocket[63,64].

### NMR experiments

NMR experiments were recorded on Bruker Avance Neo 900 MHz or Avance III HD 600 MHz spectrometers (Bruker BioSpin) equipped with cryogenically cooled triple resonance ^1^H{^13^C/^15^N} TCI probes. Data were acquired with Topspin versions 3.2 to 4.0.7 (Bruker BioSpin), processed with Topspin and NMRPipe[65], and analyzed with CCPNMR analysis 2.5.2[66].

#### 1. Resonance assignment

Experiments were initially performed on the 600 MHz spectrometer at 30°C on a ^15^N/^13^C-labeled SIRPαV1 sample (260 µM) prepared in 10 mM HEPES pH 7.4, 150 mM NaCl, 10% D_2_O and on the 900 MHz spectrometer at 27°C on a ^15^N/^13^C-labeled SIRPαV2 sample (260 µM) prepared in the same buffer. Backbone ^1^H/^15^N/^13^C assignment was obtained by standard methods from ^1^H-^15^N HSQC and 3D HNCO, HN(CA)CO, HN(CO)CACB, HNCACB, HNCA, HN(CO)CA from Topspin library for SIRPαV1 or as implemented in NMRLib 2.0[67] using BEST version of the 3D for SIRPαV2. For both isoforms, 99% of non-proline residues were assigned (only Asn51 was missing). For the Q52A and Q52F mutants, only a few residues could not be unambiguously assigned (Lys11, Asn51, Arg69, Ala84 and Ser98 for Q52A; and Lys11 and Asn51 for Q52F).

#### 2. Predictions of secondary structures and order parameters

(S^2,pred^) for SIRPαV1 and SIRPαV2 were obtained from H^N^, N, C^α^, C^β^, CO chemical shifts using the TALOS-N software[22].

#### 3. Dynamics

For apo SIRPαV1, SIRPαV2, SIRPαV2-Q52F, SIRPαV2-Q52A and holo SIRPαV2/5-HTP, ^15^N relaxation measurements were performed on the 600 MHz spectrometer at 25°C on ^15^N-labeled samples at 200 µM protein concentration and a 5-fold molar excess of 5-HTP compound for the complex. Amide (^1^H, ^15^N) resonance assignments of SIRPαV2-Q52F, SIRPαV2-Q52A and SIRPαV2/5-HTP complexes were previously obtained by analogy to SIRPαV2 and careful inspection of the behavior of the neighboring residues in case of ambiguities. Note that for 5-HTP, the theoretical portion of complex formed is only 11% at V2:ligand of 200uM:1000mM with K_D_ of ∼8 mM (estimated from the quadratic binding equation assuming a 1:1 binding model). This is in agreement with CSP intensities when comparing WT SIRPαV2 bound to 5-HTP to SIRPαV2-Q52F, where we have 100% pocket occupancy by Phe52.

The ^15^N relaxation times (T_1_ and T_2_) and {^1^H}-^15^N heteronuclear NOE were recorded by standard methods implemented in NMRLib 2.0, in an interleaved manner with a recycling time of 2 s and with eight relaxation delays for T_1_ (20, 100, 200, 400, 600, 900, 1300, 1800 ms) and for T_2_ (0, 17, 34, 51, 68, 102, 136, 204 ms). The heteronuclear NOE were recorded in the presence and absence of a 3-s ^1^H saturation sequence (120° ^1^H pulse train). The relaxation parameters were analyzed with the program TENSOR2[68] to infer global and internal motions. Due to the non-spherical shape of SIRPα domains, a fully anisotropic model was used to describe the global reorientation of the domains. The rotational diffusion tensor [Dx, Dy, Dz (1e^7^ s^-1^)] calculated from the T_1_/T_2_ ratios of non-flexible residues was: [1.68 ± 0.01, 1.72 ± 0.02, 2.58 ± 0.02] for SIRPαV1; [1.95 ± 0.02, 1.99 ± 0.02, 2.73 ± 0.03] for SIRPαV2; [1.91 ± 0.02, 2.02 ± 0.02, 2.78 ± 0.03] for SIRPαV2-Q52F; [2.12 ± 0.02, 2.16 ± 0.02, 2.71 ± 0.02] for SIRPαV2-Q52A and [2.00 ± 0.02, 2.03 ± 0.01, 2.79 ± 0.02] for SIRPαV2/5-HTP. The relaxation parameters were then analyzed using the Lipari and Szabo formalism[69] to extract internal dynamical parameters (order parameter S^2^: internal correlation time τ_e_ on the ps-ns timescale; exchange parameter R_ex_ on the μs-ms timescale).

#### 4. Chemical Shift Perturbations (CSP)

Chemical shift differences (Δδ) between SIRPαV1 and SIRPαV2 were calculated as the weighted average (^1^H, ^15^N) chemical shift differences between the two isoforms as follows: Δδ = [(Δδ(^1^H))^2^+(Δδ(^15^N)×0.159)^2^]^1/2^. To validate and characterize initial fragment hits binding to SIRPαV2, HSQC spectra were recorded on ^15^N-labeled wild-type protein (50 µM) in the presence of 10-50-fold molar excess of fragments from DMSO stocks (resulting in 0.2-1.2% final DMSO concentrations). CSP on SIRPαV2 induced by the Q52F or Q52A mutation were obtained from HSQC spectra recorded on ^15^N-labeled wild-type and mutant SIRPaV2 samples at 200 µM concentrations. CSP were calculated as the weighted average (^1^H, ^15^N) chemical shift differences between wild-type and mutant as follows: CSP = [(Δδ(^1^H))^2^+(Δδ(^15^N)×0.159)^2^]^1/2^. CSP on SIRPαV2 induced by 5-HTP were obtained using an ^15^N-labeled SIRPαV2 sample at 200 µM in the presence of a 5-fold molar excess of 5-HTP.

### Isothermal Titration Calorimetry (ITC)

All ITC experiments were carried out with a MicroCal ITC200 instrument at a temperature of 25°C, a reference power of 5μcal/sec, an initial delay of 60sec, and a constant stirring speed of 750rpm. For all experiments, protein samples were diluted in gel filtration buffer (10mM HEPES pH 7.5, 0.15M NaCl). Titrations were carried out in 23 injections of 1.5μL from the syringe to the analytical cell.

To study the SIRPα-CD47 interaction, CD47 was placed in the analytical cell at a concentration of 22μM while SIRPα (WT and mutants) protein was loaded in the syringe at a concentration of 250μM.

To study the binding of small molecules to SIRPα, the protein (WT or mutants) was placed in the analytical cell at a concentration of 200μM and the small molecules were loaded in the syringe at a concentration of 50mM in identical sample buffer as the protein (10 mM HEPES pH 7.5, 0.15M NaCl). Raw data were scaled after setting the zero to the titration saturation heat value. Integrated raw ITC data were fitted to one site nonlinear least squares fit model using MicroCal Origin 9.1 (Origin Lab).

Each experiment was performed in triplicates and data are presented as the mean +/-SD.

### Nano-Differential Scanning Fluorimetry (nanoDSF)

We measured nanoDSF with a Prometheus NT.48 instrument (Nanotemper Technologies GmBH). High-sensitivity capillaries were filled with 10 µL of protein at 22 µM and placed in the sample holder. All samples were diluted in gel filtration buffer (10 mM HEPES pH 7.5, 0.15 M NaCl). All measurements were done in triplicates. The temperature was increased from 20 to 95 °C with a ramp of 0.5°C/min and intrinsic fluorescence (350 nm and 330 nm) and backreflection (for aggregation) were measured using the PR.Thermcontrol software. Data were analyzed using the PR.StabilityAnalysis software (Nanotemper Tech.) by fitting each curve within a region of interest to derive precise Tm values.

### Surface Plasmon Resonance (SPR) Spectroscopy

SPR experiments were conducted using a Biacore T200 system. Streptavidin (SAV) sensor chips (SAS, Cytiva) were used for immobilization. All experiments were performed at 25°C. The running buffer consisted of a solution suitable for protein-ligand binding (PBS + 0.005% IGEPAL CA-630). SIRPα-biotin was immobilized onto the flow cells of the SAV sensor chip. The chip surface was conditioned with regeneration solution (3x 1min; 1M NaCl, 50mM NaOH) and SIRPα-biotin was injected at a low flow rate (10 µL/min; 200nM; 7min) to achieve an immobilization level of ∼2500-3500 Response Units (RU). One flow cell was left blank (immobilization buffer only) to serve as a reference surface for signal subtraction. Biotin was subsequently injected to saturate binding surfaces (40µg/ml; 10µl/min; 100µl). The binding partner (analyte), either CD47-CD4-6His or the small molecule ligands (e.g., Trp, 5-HTP) was diluted in running buffer to create a concentration series. For protein analytes (CD47), a range of concentrations (e.g., 6.25 nM to 200 nM) of the analyte was injected over both the SIRPα-immobilized surface and the reference surface at a flow rate of 30–50 µL/min. Association time was typically 120– 180 seconds, followed by a dissociation time of 300–600 seconds. For small molecule analytes, compounds were tested using a single-cycle kinetics or steady-state affinity approach. A series of increasing concentrations was injected sequentially without regeneration in between. The sensor surface was regenerated between cycles with a regeneration solution (10 mM Glycine pH 2.0) to completely dissociate bound analytes, ensuring the reproducibility of the measurements. The raw sensorgram data were double referenced by subtracting the signal from the reference flow cell and then subtracting the signal from a buffer blank injection. The resulting binding curves were analyzed using the Biacore evaluation software.

Kinetic Analysis. The binding data were fit to a 1:1 Langmuir binding model to determine the association rate constant (k_a_) and the dissociation rate constant (k_d_).

Affinity Constant. The equilibrium dissociation constant (K_D_) was calculated from steady-state analysis. The equilibrium response (R_eq_) was plotted against the analyte concentration and fit to a steady-state affinity model to determine the K_D_.

### Quantification and statistical analysis

The number of biological and technical replicates are indicated in relevant figure legends. Details on statistical tests, parameters and significance cutoffs are listed in relevant figure legends. Statistical analyses were performed using GraphPad Prism software, version 9.5.0 (GraphPad). Significance was defined as follows: ∗, p < 0.05; ∗∗, p < 0.01; ∗∗∗, p < 0.001; ∗∗∗∗, p < 0.0001.

## Acknowledgements

The authors would like to acknowledge the following:

The Frederick National Laboratory for Cancer Research (FNLCR) Protein Expression Laboratory, the Biomass & Protein Engineering platform at the Mediterranean Microbiology Institute, the NMR platform at the Mediterranean Microbiology Institute, the NIH S10 Grant: S10OD030350 and University of Maryland Biomolecular NMR Facility, the IR-RMN infrastructure (FR3050 CNRS) for access to NMR facilities and financial support and FX Canterelle for his expert technical assistance, the Datacentre IT and Scientific Computing platform of the CRCM for providing computational resources.

The Diamond Light Source for access to the fragment screening facility XChem, for usage of DSi-Poised library and for beamtime on beamline I04-1 under proposal SW21399, the DLS XChem Industrial Liaison Service, SOLEIL for provision of synchrotron radiation facilities using beamlines PROXIMA 1 and PROXIMA 2A under proposal 20231038, the European Synchrotron Radiation Facility (ESRF) for provision of synchrotron radiation facilities using beamlines ID30A1 and ID30B under proposal number mx2700. We would like to thank the staff of the ESRF and EMBL Grenoble for assistance and support in using the above beamline(s) under proposal number MX2490. Use of the Stanford Synchrotron Radiation Lightsource, SLAC National Accelerator Laboratory, is supported by the U.S. Department of Energy, Office of Science, Office of Basic Energy Sciences under Contract No. DE-AC02-76SF00515. The SSRL Structural Molecular Biology Program is supported by the DOE Office of Biological and Environmental Research, and by the National Institutes of Health, National Institute of General Medical Sciences (P30GM133894). The contents of this publication are solely the responsibility of the authors and do not necessarily represent the official views of NIGMS or NIH.

We also acknowledge the help of the CRCM bioinformatics platform (Cibi) in particular Ghislain Bidaut, who performed some data analysis.

Finally, the authors acknowledge the contributions of Catherine Farrell and Teresa Burgess that were critical to the inception of this research program, its funding, and intellectual advancement.

## Data and materials availability

Coordinates and structure factors for wild-type apo SIRPαV2, wild-type SIRPαV2 with Trp, wild-type SIRPαV2 with 5-HTP, Q52A SIRPαV2, Q52F SIRPαV2 have been deposited in the PDB with entry codes 9TF5, 9SIA, 9T7F, 9SIC, and 9SID, respectively.

