## Supplementary-Material for "Engineering SIRPα conformational plasticity to reveal a cryptic pocket suitable for structure-based drug design"

**Affiliations**
[1]CRCM, CNRS, Inserm, Institut Paoli-Calmettes, Aix-Marseille Univ, Marseille, France
[2]Biological NMR & HDX-MS Technological Platform, Institut Pasteur, Université Paris Cité, CNRS UMR 3528, Paris, France.
[3]Structural Bioinformatics Unit, Institut Pasteur, Université Paris Cité, CNRS UMR 3528, Paris, France.
[4]Institute of Molecular Physical Science, Swiss Federal Institute of Technology Zürich (ETHZ), Vladimir-Prelog-Weg 2, 8093 Zürich, Switzerland
[5]NexMR AG, Wiesenstrasse 10A, Schlieren, Switzerland
[6]Paradigm Shift Therapeutics LLC, Sonoma, California, United States of America
[7]Department of Chemistry and Biochemistry, University of Maryland Baltimore County, Baltimore, Maryland, USA

[†]**Current address:** [a]Institute for Integrative Biology of the Cell (I2BC, UMR9198), Computer Science Laboratory (LIX, UMR7161), Université Paris-Saclay, CNRS, Gif-sur-Yvette, France
[b]Architecture et Fonction des Macromolécules Biologiques (AFMB), Unité Mixte de Recherche (UMR) 7257, Aix-Marseille Univ, Marseille, France
[c]Institut de Chimie radicalaire (ICR) UMR7273, Équipe Pharmaco-chimie radicalaire, CNRS, Aix-Marseille Université, Marseille, France,
[d]CRCM, CNRS, Inserm, Institut Paoli-Calmettes, Aix-Marseille Univ, Marseille, France

*These authors contributed equally to this work

**Acknowledgements:** The authors would like to acknowledge the following:

The Frederick National Laboratory for Cancer Research (FNLCR) Protein Expression Laboratory, the Biomass & Protein Engineering platform at the Mediterranean Microbiology Institute, the NMR platform at the Mediterranean Microbiology Institute, the NIH S10 Grant: S10OD030350 and University of Maryland Biomolecular NMR Facility, the IR-RMN infrastructure (FR3050 CNRS) for access to NMR facilities and financial support and FX Canterelle for his expert technical assistance, the Datacentre IT and Scientific Computing platform of the CRCM for providing computational resources.
The Diamond Light Source for access to the fragment screening facility XChem, for usage of DSi-Poised library and for beamtime on beamline I04-1 under proposal SW21399, the DLS XChem Industrial Liaison Service, SOLEIL for provision of synchrotron radiation facilities using beamlines PROXIMA 1 and PROXIMA 2A under proposal 20231038, the European Synchrotron Radiation Facility (ESRF) for provision of synchrotron radiation facilities using beamlines ID30A1 and ID30B under proposal number mx2700. We would like to thank the staff of the ESRF and EMBL Grenoble for assistance and support in using the above beamline(s) under proposal number MX2490. Use of the Stanford Synchrotron Radiation Lightsource, SLAC National Accelerator Laboratory, is supported by the U.S. Department of Energy, Office of Science, Office of Basic Energy Sciences under Contract No.

DE-AC02-76SF00515. The SSRL Structural Molecular Biology Program is supported by the DOE Office of Biological and Environmental Research, and by the National Institutes of Health, National Institute of General Medical Sciences (P30GM133894). The contents of this publication are solely the responsibility of the authors and do not necessarily represent the official views of NIGMS or NIH.
We also acknowledge the help of the CRCM bioinformatics platform (Cibi) in particular Ghislain Bidaut, who performed some data analysis.
Finally, the authors acknowledge the contributions of Catherine Farrell and Teresa Burgess that were critical to the inception of this research program, its funding, and intellectual advancement.

**Funding :** This work was supported in part by the National Cancer Institute (R37CA218259; TWM), French National Research Agency ANR-22-CE18-0023 (XM), Maryland Industrial Partnerships (MIPS; #5914, EDG, TWM), Aix Marseille University Institute of Cancer and Immunology (ICI, TWM), La Ligue Contre le Cancer (MS). This work received support from the French government under the France 2030 investment plan, as part of the Initiative d'Excellence d'Aix-Marseille Université - A*MIDEX (AMX-18-ACE-004 to EDG).

**Data and materials availability.**

Coordinates and structure factors for wild-type apo SIRPαV2, wild-type SIRPαV2 with Trp, wild-type SIRPαV2 with 5-HTP, Q52A SIRPαV2, Q52F SIRPαV2 have been deposited in the PDB with entry codes 9TF5, 9SIA, 9T7F, 9SIC, and 9SID, respectively.

**Supplementary Materials.**

**Supplementary Table S1**. X-ray data collection and refinement statistics for SIRPαV2 proteins.

| Data set | WT (apo) | Q52A | Q52F | WT + Trp | WT + 5-HTP |
|---|---|---|---|---|---|
| PDB Code | 9TF5 | 9SIC | 9SID | 9SIA | 9T7F |
| Beamline | 9-2 (SSRL) | PX1 (SOLEIL) | PX2 (SOLEIL) | MASSIF-1 (ESRF) | PX1 (SOLEIL) |
| Wavelength (Å)° | 0.9795 | 0.9786 | 0.9801 | 0.9655 | 0.9786 |
| Resolution range (Å) | 43.36-1.63 (1.65-1.63) | 40.51-1.50 (1.59-1.50) | 43.80-1.69 (1.80-1.69) | 43.32-1.83 (1.89-1.83) | 49.70-1.50 (1.59-1.50) |
| Spacegroup | P41212 | P41212 | P212121 | P41212 | P41212 |
| Unit cell parameters | 57.26 57.26 199.2 90 90 90 | 57.29 57.29 200.54 90 90 90 | 57.53 58.05 66.73 90 90 90 | 57.35 57.35 198.29 90 90 90 | 57.43 57.43 198.04 90 90 90 |
| Unique reflections | 41772 (1230) | 54492 | 25431 | 29981 | 54272 |
| Completeness | 97.5 (69.3) | 100 | 99.76 | 98.68 | 100 |
| Redundancy | 8.9 (3.5) | 27.7 | 12.3 | 7.3 | 25.0 |
| Mean I/sigmaI | 8.0 (2.9) | 32.64 (5.42) | 9.95 (0.89) | 14,00 (1.98) | 19.23 (1.50) |
| Wilson Bfactor | 23.2 | 16.8 | 25.4 | 23.6 | 21.7 |
| CC1/2 | 0.99 (0.55) | 0.999 (0.984) | 0.997 (0.285) | 0.998 (0.596) | 1 (0.666) |
| Reflections used in refinement | 41772 (1230) | 51767 (3729) | 25416 (2603) | 29893 (2793) | 51557 (3717) |
| Reflections used for Rfree | 2000 (59) | 2725 (196) | 1272 (138) | 1453 (148) | 2714 (196) |
| R-work | 0.19 (0.35) | 0.18 | 0.24 | 0.22 | 0.23 |
| R-free | 0.22 (0.44) | 0.22 | 0.27 | 0.25 | 0.25 |
| Number of non-hydrogen atoms | 3041 | 2959 | 1839 | 2783 | 2833 |
| Macromolecules | 2701 | 2638 | 1733 | 2637 | 2605 |
| Ligands | 17 | 5 | 29 | 19 | 22 |
| Solvent | 323 | 316 | 90 | 142 | 206 |
| Protein residues | 354 | 344 | 227 | 341 | 340 |
| RMS (bonds) | 0.008 | 0.009 | 0.013 | 0.009 | 0.012 |
| RMS (angles) | 0.85 | 1.32 | 1.0 | 1.02 | 1.29 |
| Ramachandran favored (%) | 99.1 | 99.4 | 99.5 | 99.1 | 100 |
| Ramachandran allowed (%) | 0.9 | 0.6 | 0.5 | 0.9 | 0 |
| Ramachandran outliers (%) | 0.0 | 0 | 0 | 0 | 0 |
| Rotamer outliers (%) | 0.67 | 0 | 0 | 0 | 0 |
| Clashscore | 1.49 | 1.71 | 3.97 | 3.42 | 1 |
| Average Bfactor | 29.6 | 22.4 | 28.13 | 26.5 | 26 |
| Macromolecules | 28.8 | 21.3 | 28 | 26.4 | 24 |
| Ligands | 59.9 | 22.3 | 35 | 26.2 | 28.7 |
| Solvent | 34.7 | 32.2 | 30.5 | 27.1 | 27.5 |
| RMSZ (bonds) | 0.53 | 0.54 | 0.33 | 0.38 | 0.57 |
| RSMZ (angles) | 0.60 | 0.85 | 0.55 | 0.57 | 0.87 |

**Supplementary Table S2**. Activity of fragment hits and HSQC NMR validation. (pIC50 measured by HTRF for compounds noted by #).

| Compound ID | SMILES | Structure 2D | pIC50 AlphaScreen SIRPaV2 | HSQC SIRPαV2 |
|---|---|---|---|---|
| x0220 | Oc1ccc(CCNC(=O)c2ccccn2)cc1 | 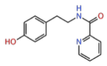 | -3.5 | negative |
| x0408 | CNC(=O)c1cccc2cc[nH]c12 | 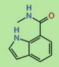 | -3.3 | positive |
| x0592 | CC(=O)Nc1ccccc1O | 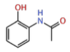 | -2.9 | negative |
| x0229 | O=S1(=O)CCN(Cc2ccco2)CC1 | 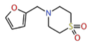 | -2.0 | negative |
| x0537 | COC(=O)N1CCN(CC1)c1ccc(F)cc1 | 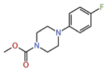 | -2.8 | negative |
| 5-hydroxy-L-Tryptophan (5-HT) | N[C@@H](Cc1c[nH]c2ccc(O)cc12)C(=O)O | 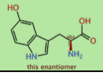 | -2.7 | positive |
| x0098 | CC(=O)NC1CNc2ccccc2C1 | 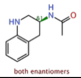 | -2.3 | negative |
| x0530 | Fc1ccc(OCC(=O)N2CCCC2)cc1 | 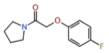 | -2.3 | negative |
| L-Tryptophan (Trp) | N[C@@H](Cc1c[nH]c2ccccc12)C(=O)O | 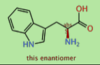 | -2.2 | positive |
| x0284 | Cl.N[C@H]1CCN(C1)S(=O)(=O)c1ccccc1 | 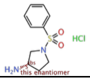 | -2.1 | negative |
| x0330 | CN1CCC(Oc2cccc(F)c2)C1=O | 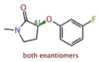 | -2.1 | negative |
| x0576 | Cl.CNC(C)c1ccc(F)cc1 | 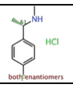 | -2.1 | negative |
| x0230 | O=C(COc1ccccc1)N1CCNC(=O)C1 | 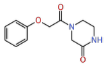 | -2.0 | negative |
| x0104 | CS(=O)(=O)NCCc1ccc(F)cc1 | 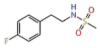 | inactive | negative |
| x0109 | NC(=O)C1CCN(CC1)C(=O)Nc1ccccc1 | 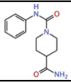 | inactive | negative |
| x0563 | CC(Oc1ccc(cc1)C#N)C(N)=O | 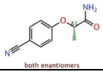 | inactive | negative |
| x0422 | Fc1cccc(CN2CCCS2(=O)=O)c1 | 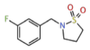 | inactive | negative |
| x0234 | Cc1c(Cl)cccc1NC(=O)[C@@H]1CCCO1 | 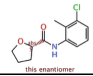 | inactive | negative |
| x0250 | Fc1ccc(NC(=O)CN2CCCC2)cc1 | 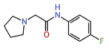 | -2.3 (#) | negative |
| x0083 | NCc1cc(no1)-c1ccccc1 | 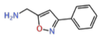 | -2.5 (#) | negative |
| x0130 | CC(=O)NC1CNC=2C=CC=CC2C1 | 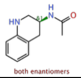 | not tested | negative |
| x0195 | CCN1C=C(C=N1)C(=O)NCC=2C=CC(F)=CC2 | 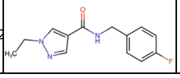 | not tested | not tested |

**Supplementary Table S3. Primers used to design mutant proteins.**

| Primer Name | Primer Sequence (5' to 3') |
|---|---|
| **SIRPaV2Q52F_For** | 5'-cacgtgggaaatgaccttctttaaagttatagatcagttcacgagcc-3' |
| **SIRPaV2Q52F_Rev** | 5'-ggctcgtgaactgatctataactttaaagaaggtcatttcccacgtg-3' |
| **SIRPaV2Q52A_For** | 5'-gtgggaaatgaccttctttcgcgttatagatcagttcacgag-3' |
| **SIRPaV2Q52A_Rev** | 5'-ctcgtgaactgatctataacgcgaaagaaggtcatttcccac-3' |

**Supplementary Figure S1: Positive fragment hits and event maps.**
The Pandda event map is shown as a light blue mesh contoured at 0.8 sigma. X083-X098: high occupancy; X0537-X0592: high occupancy for one moiety of the compound; X0109->X0422: low occupancy; X0229: site 1 (low occupancy) + site 2 (low occupancy); All structures are shown in the same orientation as **Figure 2**.

**Supplementary Figure S2: Fragment hits inhibitory activity of the SIRPα-CD47 interaction.**
**(A)** Dose-response curves assay showing the inhibitory activity of compounds x0408, L-Tryptophan (Trp), and 5-hydroxy-tryptophan (5-HTP) on SIRPαV2-CD47 interaction (AlphaLISA, normalized activity, n=3). **(B)** Dose-response curves assay showing the inhibitory activity of weak hits (AlphaLISA, except HTRF for x0083* and x0250*).

**Suppl. Figure S3: NMR assignments and chemical shift differences between SIRPαV1 and SIRPαV2. (A, B)** $^{1}H$–$^{15}N$ HSQC spectra of uniformly $^{15}N$/$^{13}C$-labeled SIRPαV1 **(A)** and SIRPαV2 **(B)**, recorded at 260 µM concentration on 600 MHz and 950 MHz spectrometers, respectively. Backbone amide assignments are indicated in grey (99% of non-Proline residues, only Asn51 was missing in both isoforms). **(C)** Backbone amide chemical shift differences (Dd) between SIRPαV1 and SIRPαV2. Amino acid differences between the two variants are marked with open red circles. Secondary structure elements and loops are indicated at the top. **(D)** Mapping of Dd values on our SIRPαV2 structure (PDB code 9TF5), with color ranging from blue to red with increasing Dd values, as shown in panel **C**, and grey for missing data. Gln52 and Ser66 side chains are shown as sticks. Residues that differ between SIRPαV1 and SIRPαV2 are shown as red spheres.

**Suppl. Figure S4: NMR validation of indole hits.** HSQC spectrum for SIRPαV2 without and with 10-50-fold molar excess of **(A)** x0408, **(B)** Trp, or **(C)** 5-HTP. Chemical shifts for peaks within or near the WYF pocket are magnified as insets.

**Suppl. Figure S5: Affinity of ligands (Trp and 5-HTP) for SIRPαV1 and SIRPαV2.** Ligand affinity was measured for (A) Trp with SIRPαV2, (B) Trp with SIRPαV1, (C) 5-HTP with SIRPαV2, and (D) 5-HTP with SIRPαV1, by SPR at various concentrations (500-8000 µM) using the steady state approximation (n=3).

**Suppl. Figure S6: Affinity of ligands for SIRPαV2 WT and Q52 mutants.** Ligand affinity was measured for (**A**) Trp and SIRPαV2 proteins (n=1 for Q52R and Q52K mutants), and (**B**) 5-HTP with SIRPαV2 WT proteins using ITC (n=3).

**Suppl. Figure S7: $^{15}N$ relaxation parameters of SIRPαV1 and SIRPαV2 and predicted dynamics.** **(A)** $^{15}N$ relaxation times $T_1$ (top) and $T_2$ (middle) and {$^{1}H$}-$^{15}N$ NOE (bottom) for SIRPαV1 (black) and SIRPαV2 (red). Amino acid differences between the two variants are marked with open red circles. Secondary structure elements and loops are indicated at the top. **(B, C)** Backbone dynamics parameters of SIRPαV1 (black) and SIRPαV2 (red), extracted from $^{15}N$ relaxation data using the Lipari-Szabo model-free approach with an anisotropic global reorientation model: **(B)** order parameter $S^2$ probing motions on the ps-ns timescale and **(C)** exchange contribution $R_{ex}$ probing motions on the ms-

Secondary structure elements and the four loops are indicated at the top. **(D, E)** Fast motions ($S^2$) mapped on the structures of SIRPαV1 (PDB code 6NMU) and SIRPαV2 (our structure, PDB code 9TF5), with the color coding as indicated on panel **B**, i.e., from blue to red for increasing ps-ns dynamics (grey for missing data). Side chains of Gln52 and Leu/Ser66 are highlighted as sticks. **(F)** Predicted order parameters ($S^{2,pred}$) derived from backbone $^1H$, $^{15}N$ and $^{13}C$ chemical shifts using TALOS-N (Shen & Bax 2013) for SIRPαV1 (black) and SIRPαV2 (red). Amino acid differences between the two variants are marked with open red circles. Secondary structure elements and loops are indicated at the top. **(G, H)** Mapping of predicted $S^{2,pred}$ values onto the structures of SIRPαV1 (PDB code 6NMU) and SIRPαV2 (our structure, PDB code 9TF5) with the color-coding from blue (high $S^{2,pred}$, rigid) to red (small $S^{2,pred}$, flexible) as shown in panel **F**. Gln52 and Leu/Ser66 are highlighted as sticks.

**Suppl. Figure S8: $^{15}$N relaxation parameters of SIRPαV2 and 5-HTP-bound SIRPαV2.** $^{15}N$ relaxation times $T_1$ (top) and $T_2$ (middle) and $\{^1H\}$-$^{15}N$ NOE (bottom) for SIRPαV2 apo (black) and in presence of a 5-fold excess of 5-HTP ligand (cyan). Secondary structure elements and loops are indicated at the top.

**Suppl. Figure S9: Root-mean-square deviation (RMSD) of protein backbone atoms during 2-µs molecular dynamics simulations.** RMSD values, calculated using all Cα atoms relative to the equilibrated structure, are plotted as a function of simulation time for variants SIRPαV1 (**A**) and SIRPαV2 (**B**). The overall average fluctuation is indicated for each variant.

**Suppl. Figure S10: Root-mean-square fluctuation (RMSF) of protein backbone atoms during 2-µs molecular dynamics simulations.** RMSF values, calculated using all Cα atoms relative to the average structure, are plotted as a function of the protein sequence for variants SIRPαV1 (**A**) and SIRPαV2 (**B**). The most flexible loops are highlighted, and the 10 most representative conformations observed during the simulation are shown. Loop colors match those in Figure **1**.

**Suppl. Figure S11: Clustering of the representative conformations observed during 2-µs molecular dynamics simulations.** Cartoon representations of the 10 most representative conformations observed during the MD simulation for variants SIRPαV1 (**A**) and SIRPαV2 (**B**) are shown. The percentage indicates the proportion of simulation frames corresponding to each conformation. Loop colors match those in Figure **1**.

**Suppl. Figure S12: Binding site opening during 2-µs molecular dynamics simulations.** Interatomic distances between Cα atoms of Glu54-Thr67 and Gln52-Phe74 are plotted for variants SIRPαV1 (**A**) and SIRPαV2 (**B**), representing the relative conformational sampling of loops C'D and DE. The distances sampled during the MD simulations encompass the range observed in all known x-ray structures (indicated by PDB entry numbers).

**Suppl. Figure S13: Binding site volume during 2-µs molecular dynamics simulations.** Distributions of the binding site volumes for variants SIRPαV1 (**A**) and SIRPαV2 (**B**) are shown. The measured volume includes both the entrance pocket and the deep WYF pocket. The average volumes are 425.1 ± 160.4 $Å^3$ for SIRPαV1 and 433.2 ± 165.2 $Å^3$ for SIRPαV2, indicating similar overall sizes. Boxplot representations of binding site volumes (**C**) and number of pockets (**D**) are also provided.

**Suppl. Figure S14: Conformation sampling of Phe74 during 2-µs molecular dynamics simulations.** Two-dimensional plots of χ1 (x-axis) versus χ2 (y-axis) dihedral angles for variants SIRPαV1 (**A**) and SIRPαV2 (**B**) reveal two predominant side chain conformations of Phe74. The most frequent one, with Phe74 positioned outside the WYF pocket, is observed in all reported x-ray structures (indicated by PDB entry numbers). The alternative conformation, in which Phe74 is buried within the

WYF pocket, is observed in approximately 30 percent of the MD-sampled structures and has never been detected in any known x-ray structure to date.

**Suppl. Figure S15: WYF pocket conservation among SIRPα isoforms and paralogs**
The SIRPαV1 sequence of the D1 domain was used for a BLAST search using Clustered NR database, and limited to boreoeutheria and birds. We retrieved 1410 sequences with E-values better than $10^{-20}$. These sequences were aligned with Clustal Omega, and displayed with Alignment Viewer (alignmentviewer.org). A few sequences are displayed above, with the sequence logo above the reference SIRPαV1 sequence. Arrows show amino acids Trp38, Tyr50, and Phe74 forming the WYF pocket, and Gln52 from the C'D loop in SIRPαV2.

**Suppl. Figure S16: NMR backbone assignments of SIRPαV2 Q52F and Q52A mutants. (A)** $^{1}H-^{15}N$ HSQC spectra of $^{15}N$-labeled SIRPαV2 WT (black) and Q52F (red) and **(B)** SIRPαV2 WT (black) and Q52A (blue), recorded at 200 µM concentration on 600 MHz spectrometer. Backbone amide assignments of the mutants are indicated in red and blue, respectively.
Note that a second conformation can be detected for some resonances (e.g. indole $Trp38^{NHe1}$) and is consistently observed on $^{1}H-^{15}N$ HSQC spectra at 600 MHz for all SIRPα constructs, including SIRPαV1, SIRPαV2 WT, the Q52F and Q52A mutants, as well as 5-HTP-bound SIRPαV2. Detailed analysis of these additional peaks, including assignment, intensity profiles, chemical shift differences relative to the major conformation and relaxation parameters $T_1$, $T_2$ (data not shown), indicates that they correspond to residues located in spatial proximity to Trp38 and that this second conformation displays similar ps-ns and ms-ms timescale dynamics as the main conformation and accounts for approximatively 20-40% of SIRPα depending on the construct.

**Suppl. Figure S17: $^{15}N$ relaxation parameters of SIRPαV2 Q52F and Q52A mutants. (A, B)** $^{15}N$ relaxation times $T_1$ (top) and $T_2$ (middle) and $\{^{1}H\}$-$^{15}N$ NOE (bottom) for SIRPαV2 WT (black) and SIRPαV2-Q52F (red in panel A) and SIRPαV2-Q52A (blue in panel B). Secondary structure elements and loops are indicated at the top, and Q52F and Q52A mutations by the red and blue stars.

**Suppl. Figure S18. Effect of Gln52 mutations on WYF pocket volume during molecular dynamics simulations.** Gln52 was mutated *in silico* in SIRPαV2 to either alanine (Q52A) or phenylalanine (Q52F), and 50-ns MD simulations were performed to analyze the impact of these substitutions. For each construct (wild type (WT), Q52A, and Q52F), the distribution of binding pocket volumes is shown on the left, while representative snapshots of pockets detected by MDPocket are displayed on the right. The average pocket volume increases in Q52A, where the pocket is not occupied by the Gln52 side chain. The Q52F average volume is similar to WT because, although the WYF pocket is closed and blocked by Phe52, the enlarged entrance pocket keeps the total volume comparable.

**Suppl. Figure S19. Biased-open Gln52 mutants impair the high-affinity interaction with CD47.**
**(A)** Isothermal Titration Calorimetry (ITC) analysis of the SIRPα-CD47 interaction. Representative ITC thermograms show the binding of wild-type (WT) SIRPαV2 and its mutants (Q52A, Q52R, Q52F) to CD47. **(B)** Thermodynamic profile of the complex formation between CD47 and SIRPαV2 WT and mutants. (**C-E**) HTRF competition binding assay that measures the ability of non-tagged SIRPα variants (competitors) to disrupt the interaction between biotin-tagged SIRPα and CD47. (**F-H**) AlphaLISA competition binding assay that measures the ability of non-tagged SIRPα variants (competitors) to disrupt the interaction between biotin-tagged SIRPα and CD47.

**Supplementary Figure S20. Enhanced binding of 5-HTP for SIRPα Q52 mutants observed by CIDNP NMR.**
The figure shows the binding curve for 5-HTP with WT SIRPαV2 and Q52A and Q52R SIRPαV2 mutants. The scattered data points are represented instead of the fitted curve when the fitting was not

possible. In each case 100 μM of protein was titrated with increasing amount of ligand from 190 μM to 10 mM. The sample also contained either 100 μM of Fluorescein or 20 μM of AT-12, 200 nM GOCAT, 2.5 mM Glucose. The samples were irradiated for 2 s.

**Supplementary Figure S21. Gln52 dihedral angle analysis of SIRPαV1 and SIRPαV2 variants during MD simulations.**
(**A, B**) Backbone φ/ψ dihedral distributions of Gln52 sampled over the MD trajectories for SIRPαV1 (**A**) and SIRPαV2 (**B**). Density plots illustrate the conformational space explored during the MD simulations. φ/ψ values derived from available X-ray structures (indicated by PDB entry numbers) are overlaid in brown, showing that the simulations for both variants sample the conformations observed experimentally. In SIRPαV2, an additional minor conformation (~4%) not present in any known X-ray structure is observed, whereas this state is not sampled in SIRPαV1. The φ/ψ values corresponding to the most populated conformational basin in each trajectory are indicated.
(**C, D**) Sidechain χ1/χ2 dihedral angle distributions for residue Gln52 in SIRPαV1 (**C**) and SIRPαV2 (**D**). The nine major χ1/χ2 rotamers identified along the trajectories are shown, with the percentage occupancy of each rotameric state indicated. For both variants, the overall rotameric landscape is similar between V1 and V2 and the sampled rotamers collectively span the conformations observed in available X-ray structures.
Bound and Ground indicate the x-ray structures for SIRPαV2 without (ground) and with fragment x0408 (bound).

#### Figure S1

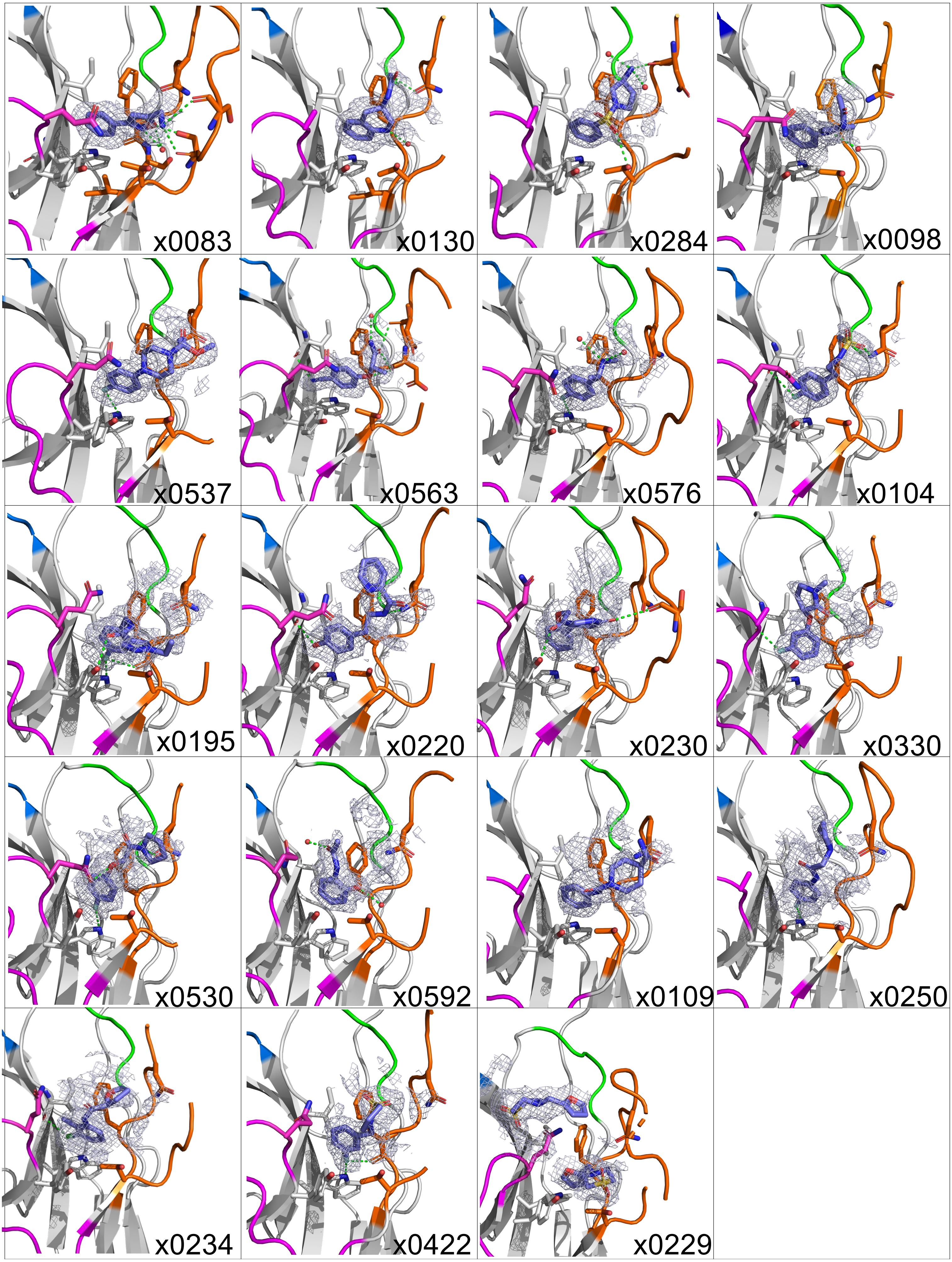

x0083
x0130
x0284
x0098
x0537
x0563
x0576
x0104
x0195
x0220
x0230
x0330
x0530
x0592
x0109
x0250
x0234
x0422
x0229

### Figure S2

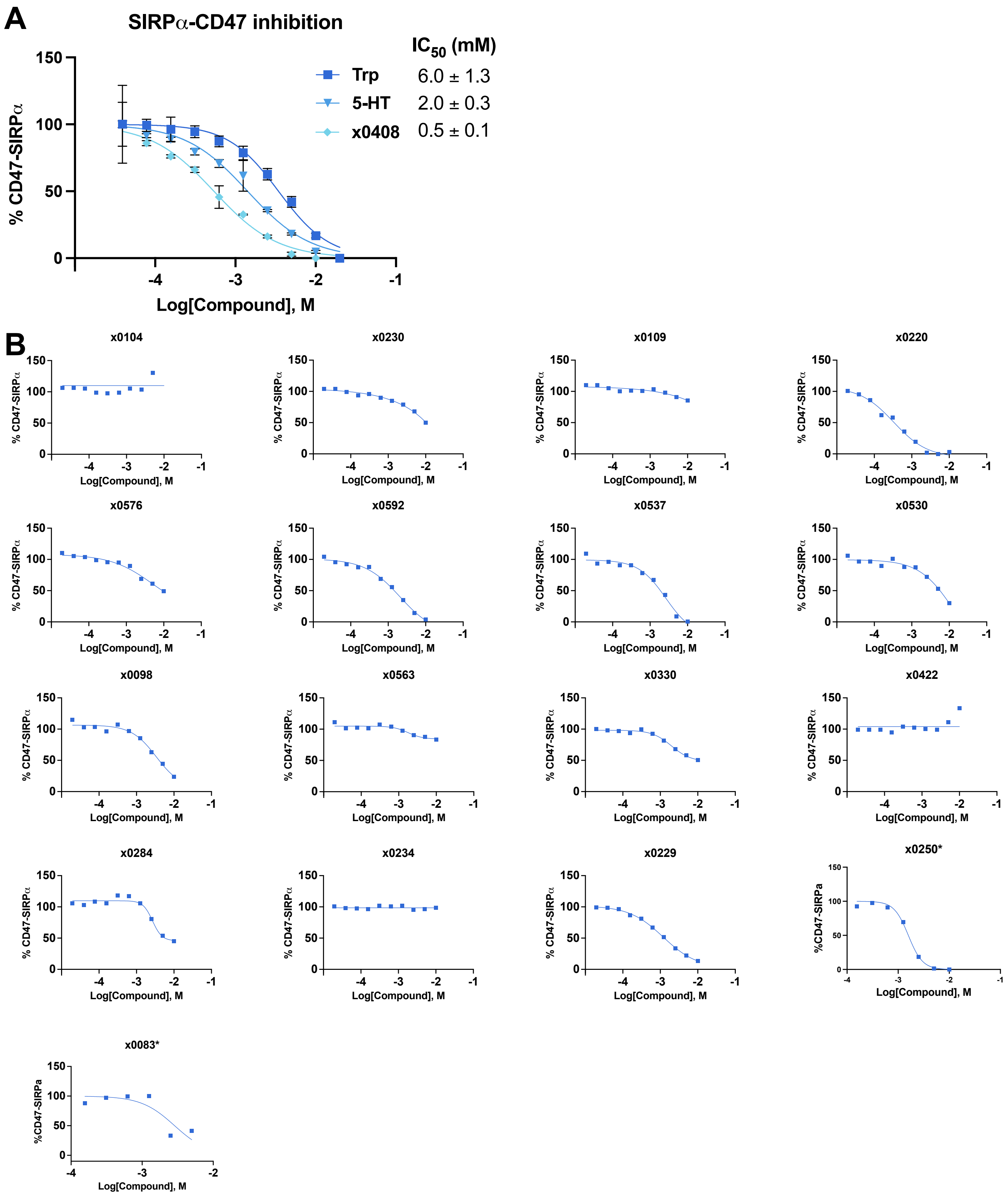
A
SIRPα-CD47 inhibition
IC50 (mM)
Trp 6.0 ± 1.3
5-HT 2.0 ± 0.3
x0408 0.5 ± 0.1
% CD47-SIRPα
Log[Compound], M
B
x0104
x0230
x0109
x0220
x0576
x0592
x0537
x0530
x0098
x0563
x0330
x0422
x0284
x0234
x0229
x0250*
x0083*
%CD47-SIRPa
Log[Compound], M

### Figure S3

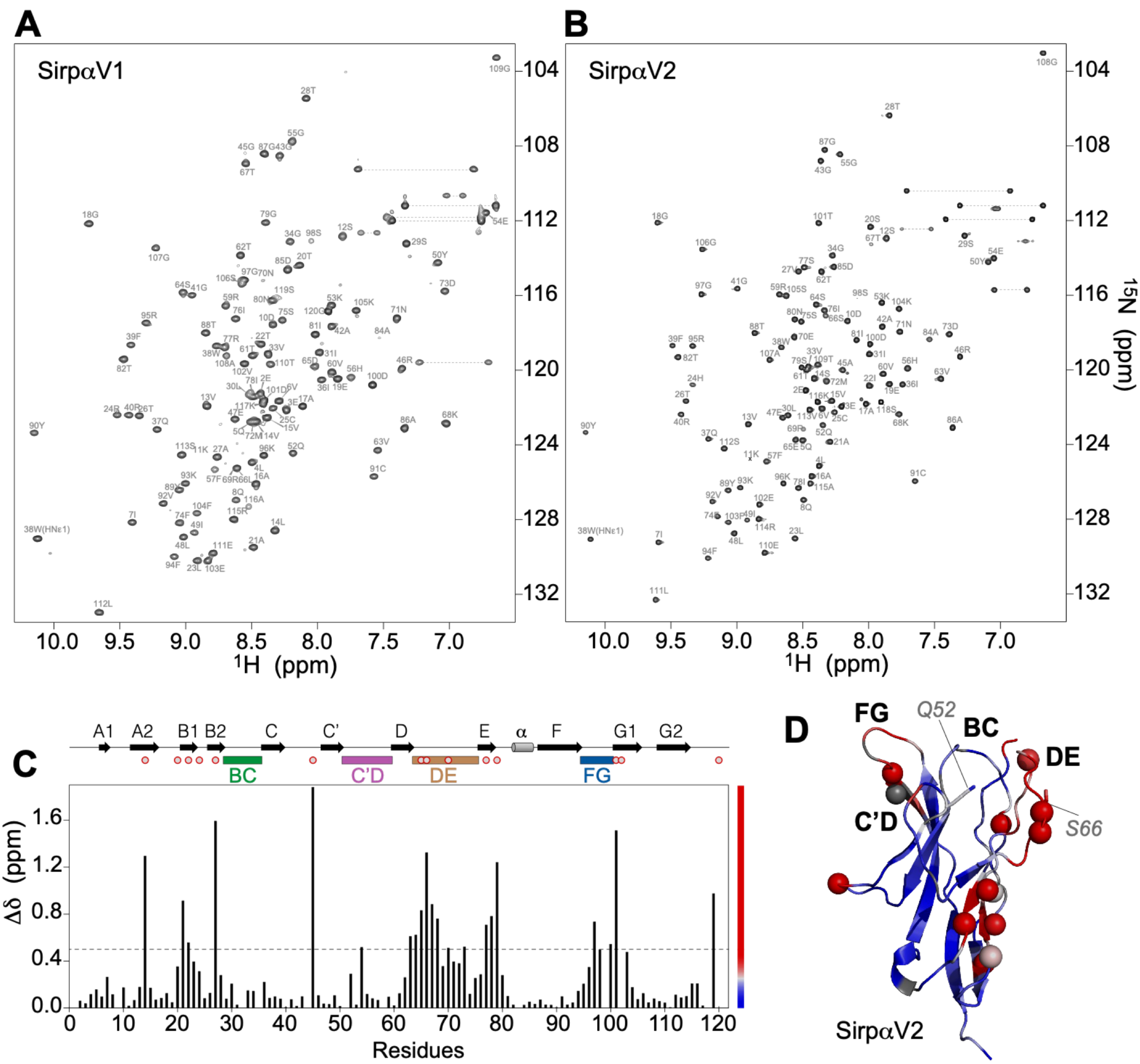
A
SirpαV1
B
SirpαV2
$^1$H (ppm)
$^{15}$N (ppm)
C
Δδ (ppm)
Residues
A1
A2
B1
B2
C
C'
D
E
α
F
G1
G2
BC
C'D
DE
FG
D
FG
Q52
BC
DE
C'D
S66
SirpαV2

### Figure S4

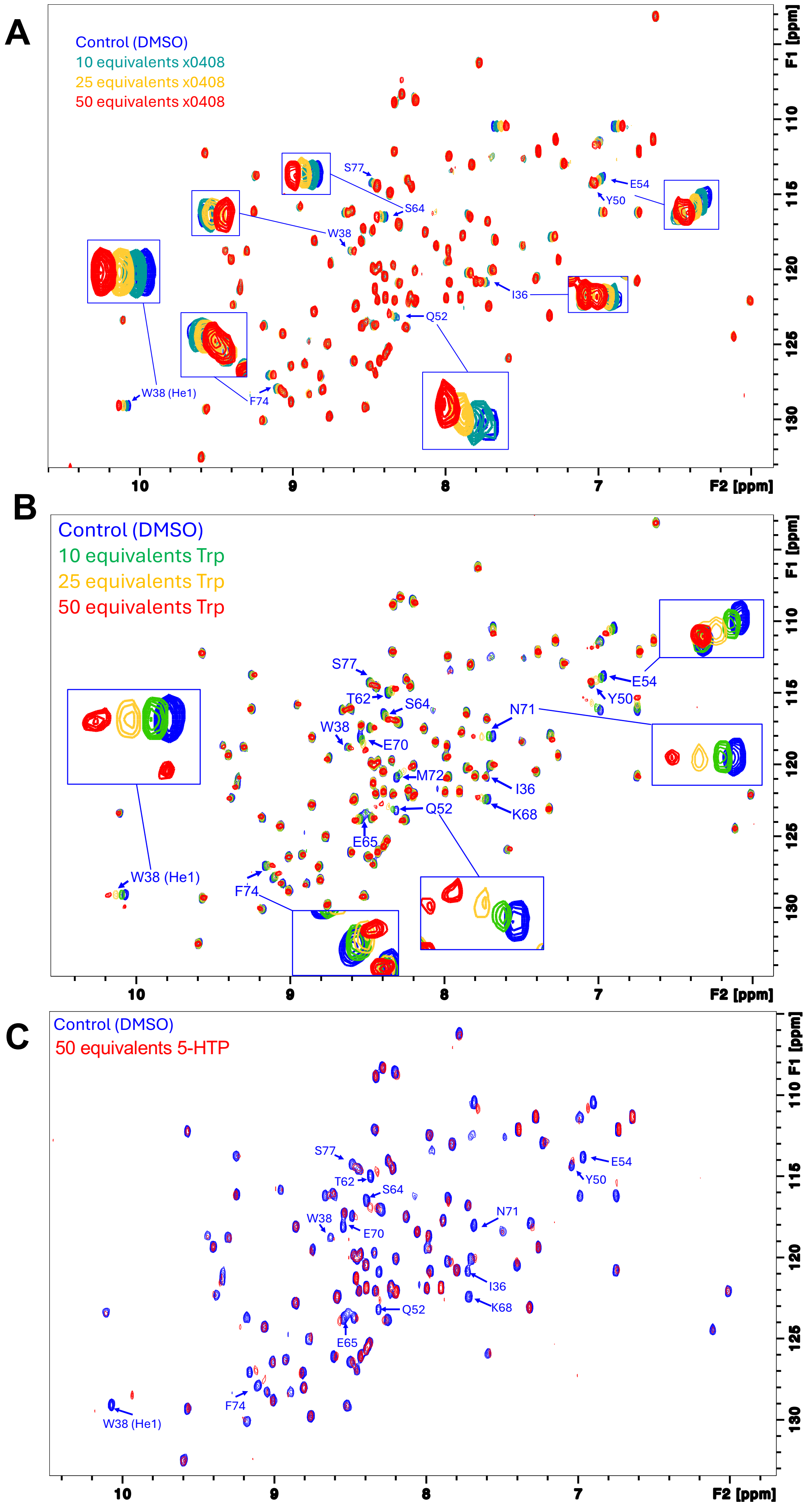
A
Control (DMSO)
10 equivalents x0408
25 equivalents x0408
50 equivalents x0408
S77
S64
W38
E54
Y50
I36
Q52
W38 (He1)
F74
F1 [ppm]
F2 [ppm]
B
Control (DMSO)
10 equivalents Trp
25 equivalents Trp
50 equivalents Trp
S77
T62
S64
W38
E70
N71
M72
I36
Q52
K68
E65
E54
Y50
W38 (He1)
F74
C
Control (DMSO)
50 equivalents 5-HTP
S77
T62
S64
W38
E70
N71
I36
Q52
K68
E65
E54
Y50
F74
W38 (He1)

### Figure S5

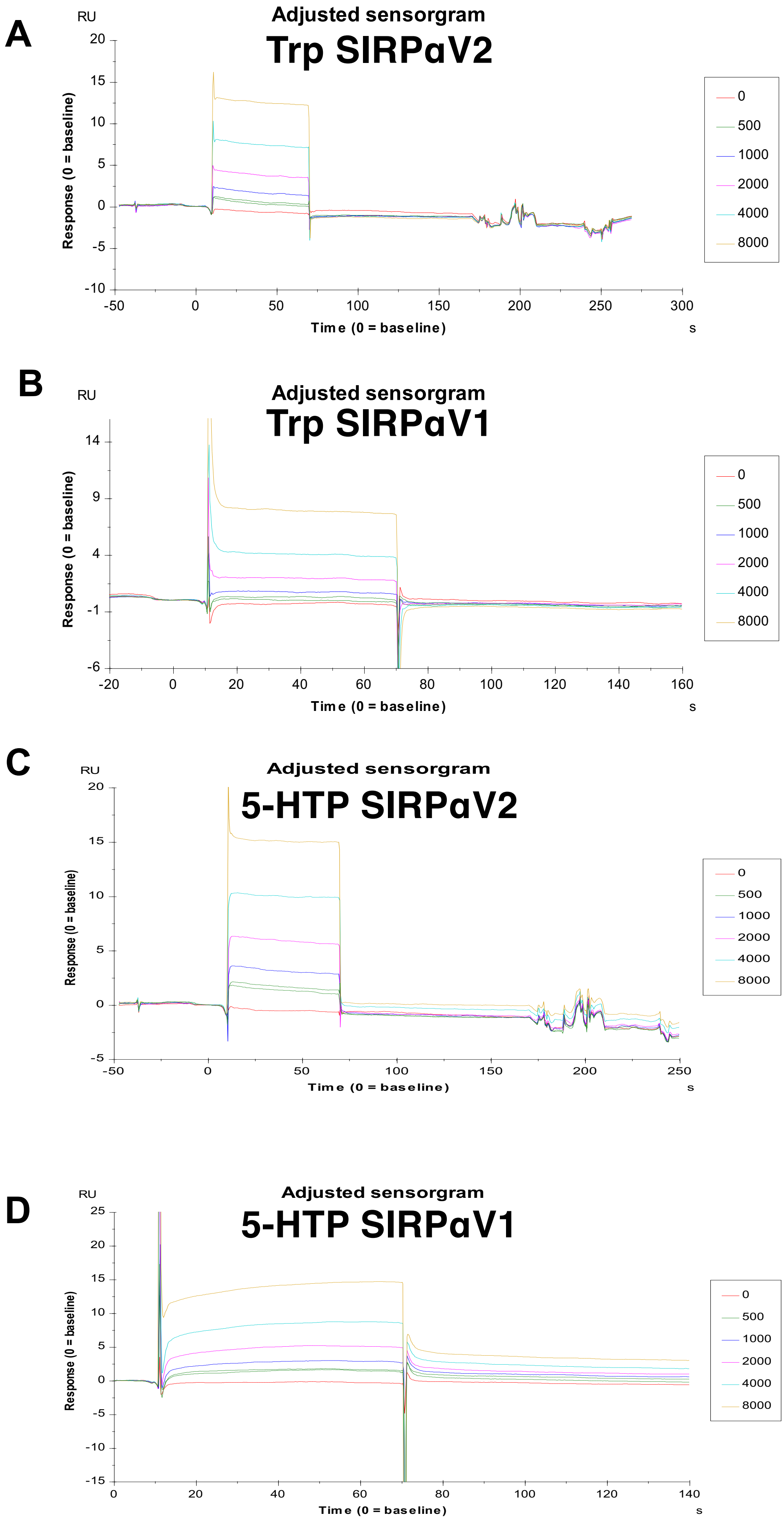
A
Adjusted sensorgram
Trp SIRPαV2
RU
Response (0 = baseline)
20
15
10
5
0
-5
-10
-50
0
50
100
150
200
250
300
Time (0 = baseline)
s
0
500
1000
2000
4000
8000
B
Adjusted sensorgram
Trp SIRPαV1
RU
Response (0 = baseline)
14
9
4
-1
-6
-20
0
20
40
60
80
100
120
140
160
Time (0 = baseline)
s
0
500
1000
2000
4000
8000
C
Adjusted sensorgram
5-HTP SIRPαV2
RU
Response (0 = baseline)
20
15
10
5
0
-5
-50
0
50
100
150
200
250
Time (0 = baseline)
s
0
500
1000
2000
4000
8000
D
Adjusted sensorgram
5-HTP SIRPαV1
RU
Response (0 = baseline)
25
20
15
10
5
0
-5
-10
-15
0
20
40
60
80
100
120
140
Time (0 = baseline)
s
0
500
1000
2000
4000
8000

### Figure S6

**A**

#### SIRPαV2-TRP

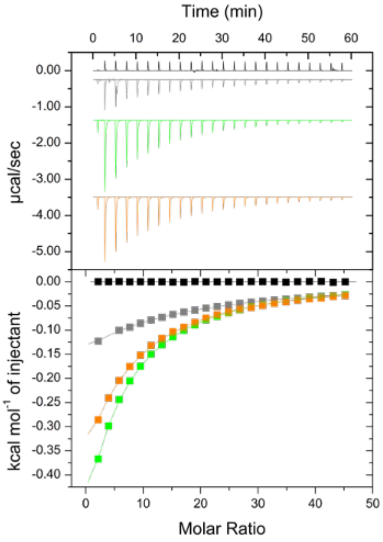


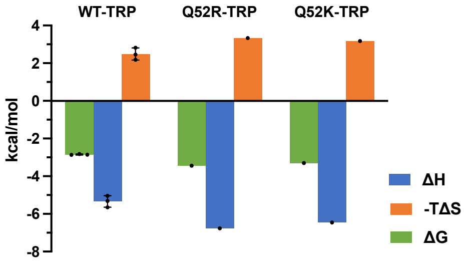


| SIRPαV2 | N +/- SD | $K_D$ (mM) |
|---|---|---|
| WT | **1 (fixed)** | **8,12 +/- 0,32** |
| Q52R | 1 (fixed) | 2,99 |
| Q52K | 1 (fixed) | 3,82 |

**B**

#### SIRPαV2-5HTP

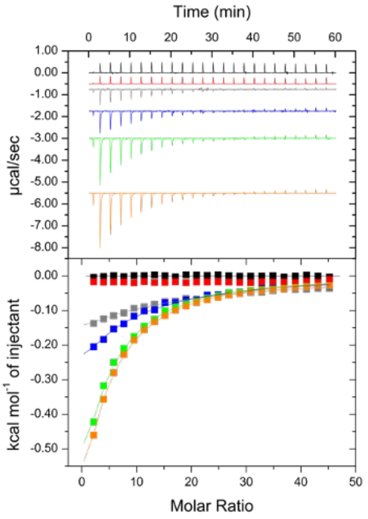


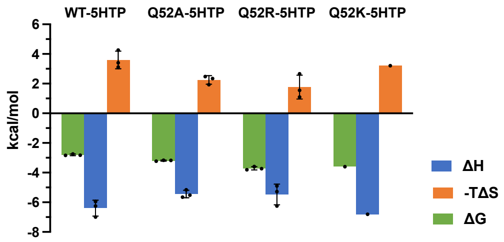


| SIRPαV2 | N +/- SD | $K_D$ (mM) |
|---|---|---|
| WT | **1 (fixed)** | **8,80 +/- 0,87** |
| Q52F | / | / |
| Q52A | 1 (fixed) | 4,55 +/- 0,22 |
| Q52R | 1 (fixed) | 1,91 +/- 0,37 |
| Q52K | 1 (fixed) | 2,27 |

### Figure S7

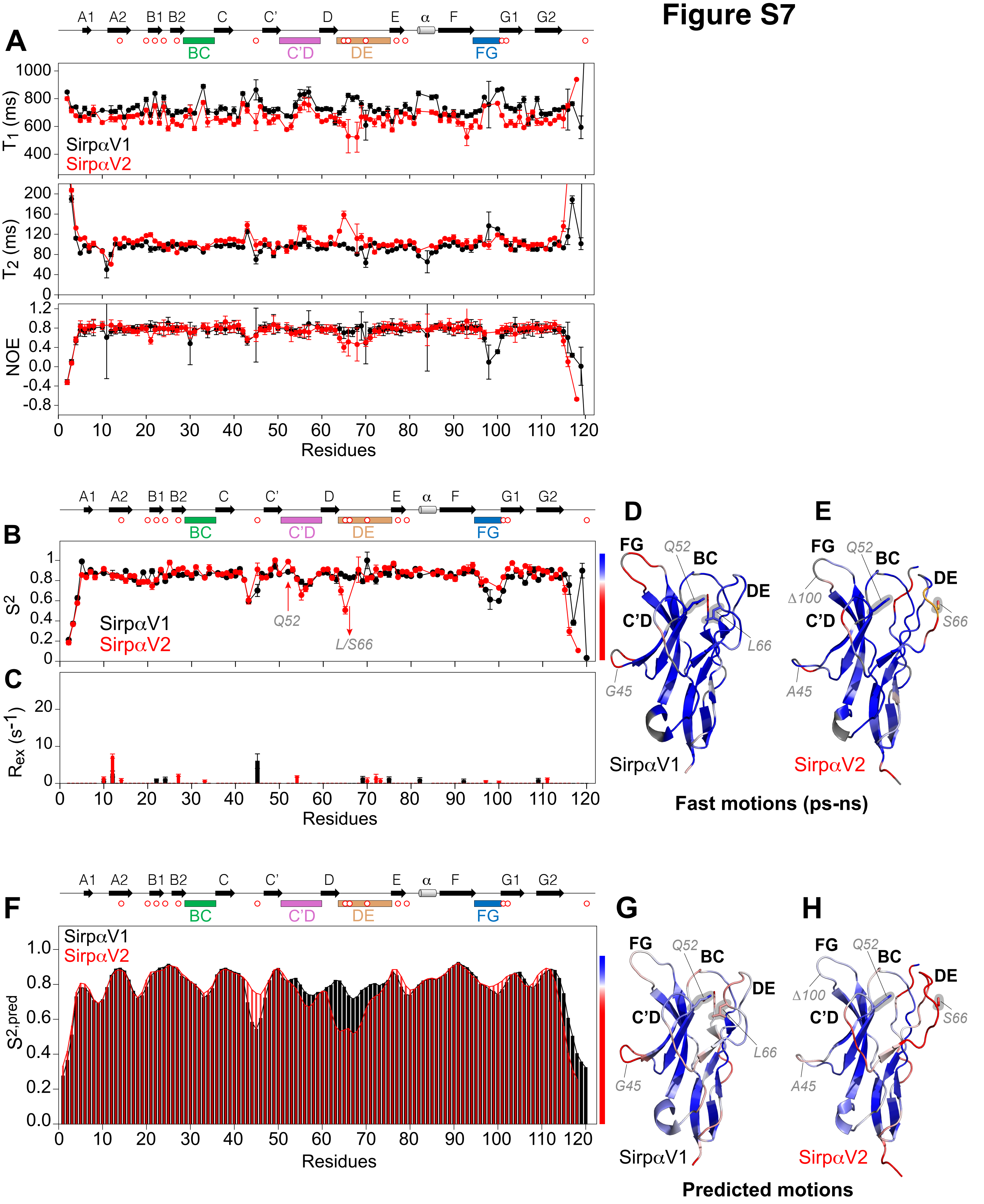

A
A1 A2 B1 B2 C C' D E α F G1 G2
BC
C'D
DE
FG
$T_1$ (ms)
SirpαV1
SirpαV2
$T_2$ (ms)
NOE
Residues
B
$S^2$
Q52
L/S66
C
$R_{ex}$ ($s^{-1}$)
D
E
Q52
G45
L66
Δ100
A45
S66
Fast motions (ps-ns)
F
$S^{2,pred}$
G
H
Predicted motions

### Figure S8

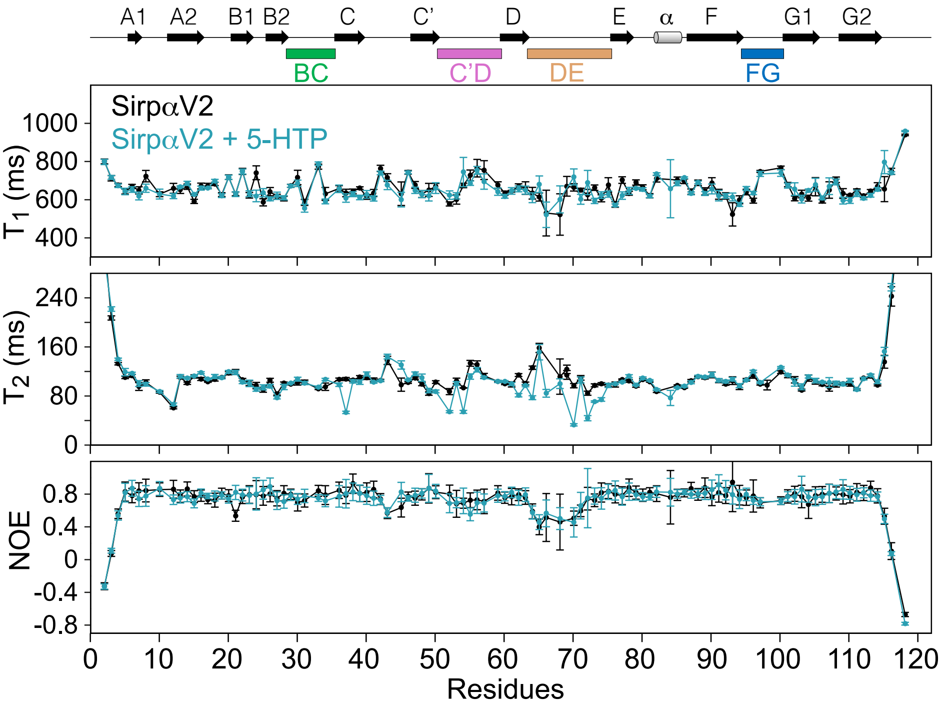

A1
A2
B1
B2
C
C'
D
E
α
F
G1
G2
BC
C'D
DE
FG
SirpαV2
SirpαV2 + 5-HTP
T1 (ms)
1000
800
600
400
T2 (ms)
240
160
80
0
NOE
0.8
0.4
0
-0.4
-0.8
0
10
20
30
40
50
60
70
80
90
100
110
120
Residues

#### Figure S9

RMSD variation (CA atoms) during trajectories

A

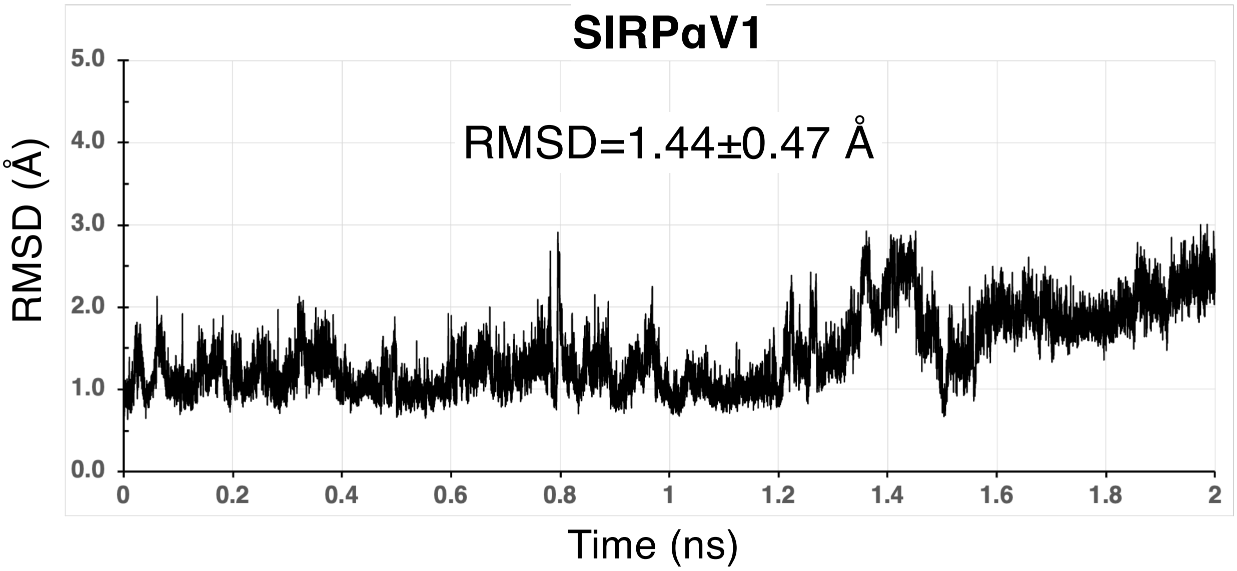


B

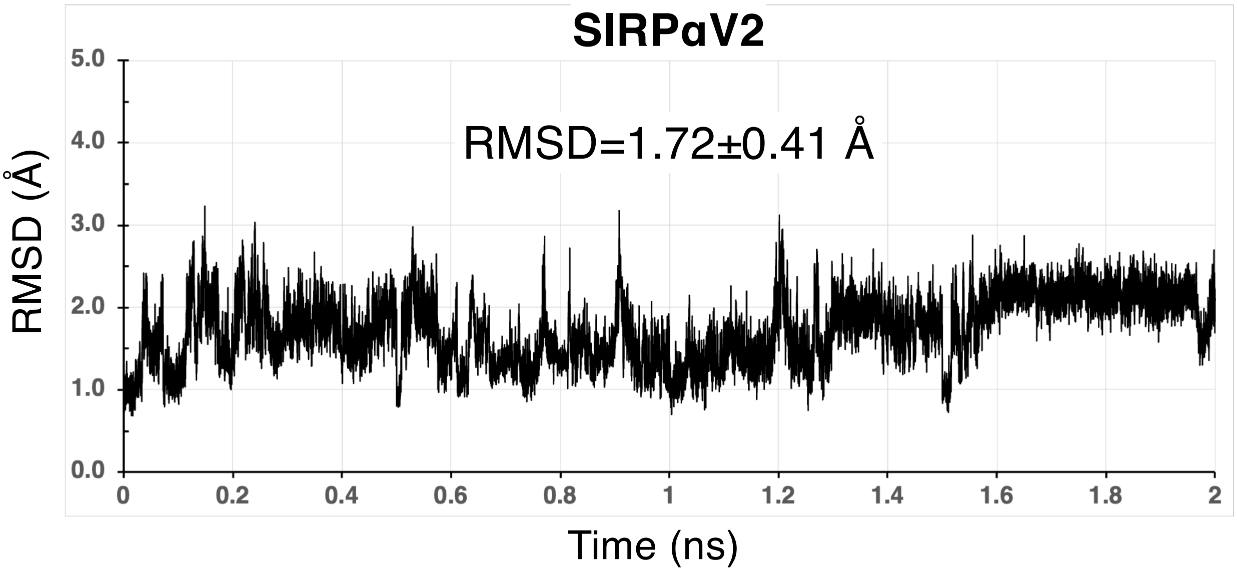

#### Figure S10

RMSF variation (CA atoms) during trajectories

**A**

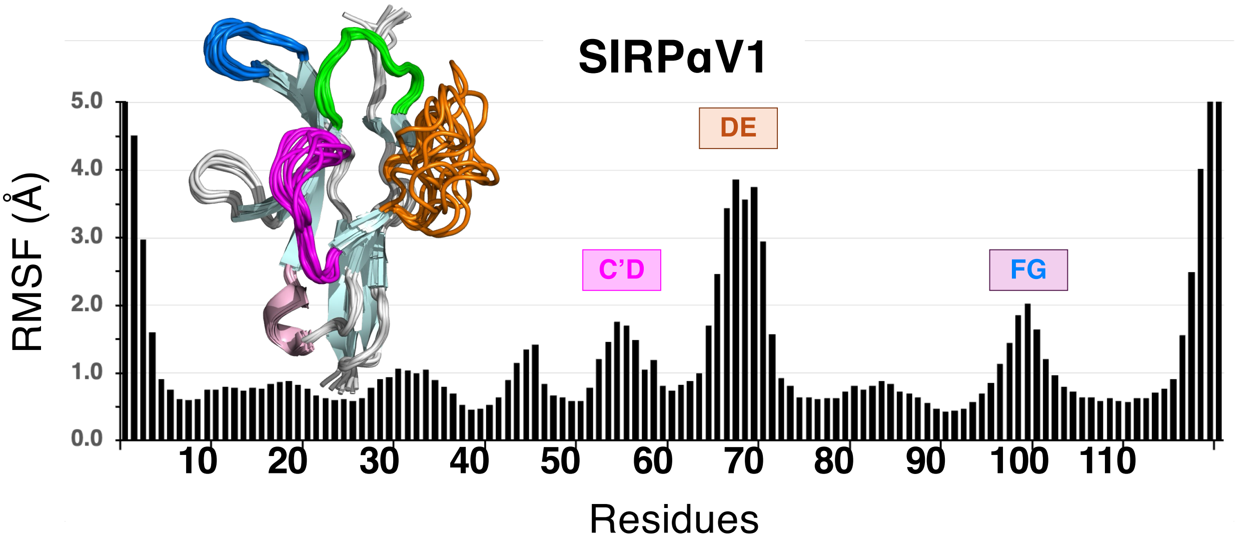


**B**

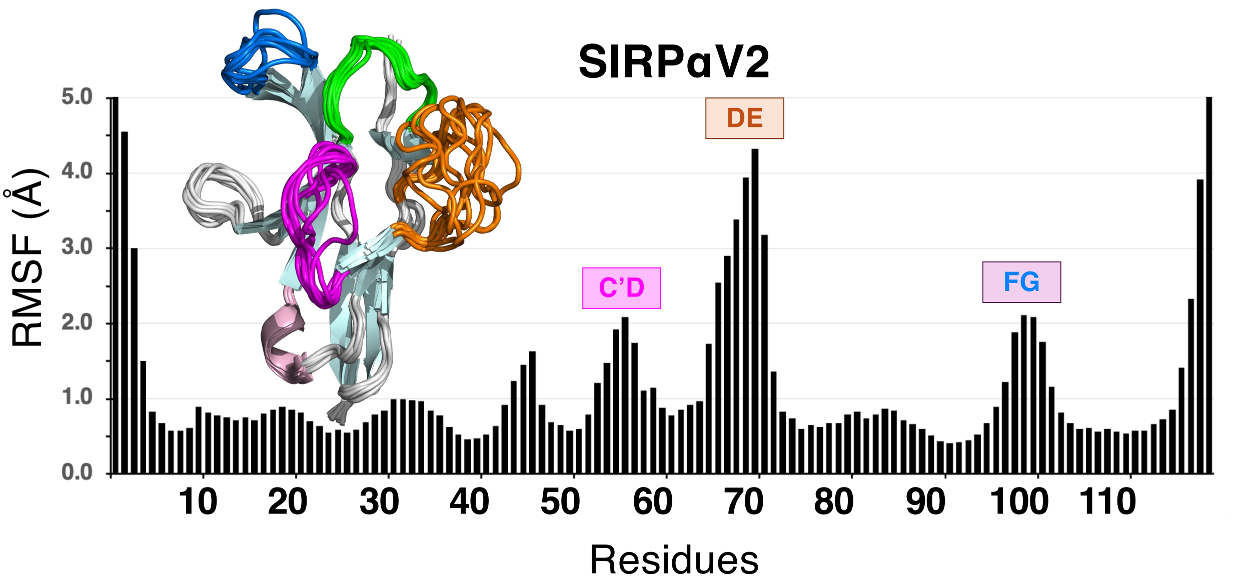

### Figure S11

Conformation clustering

**A**

#### SIRPαV1

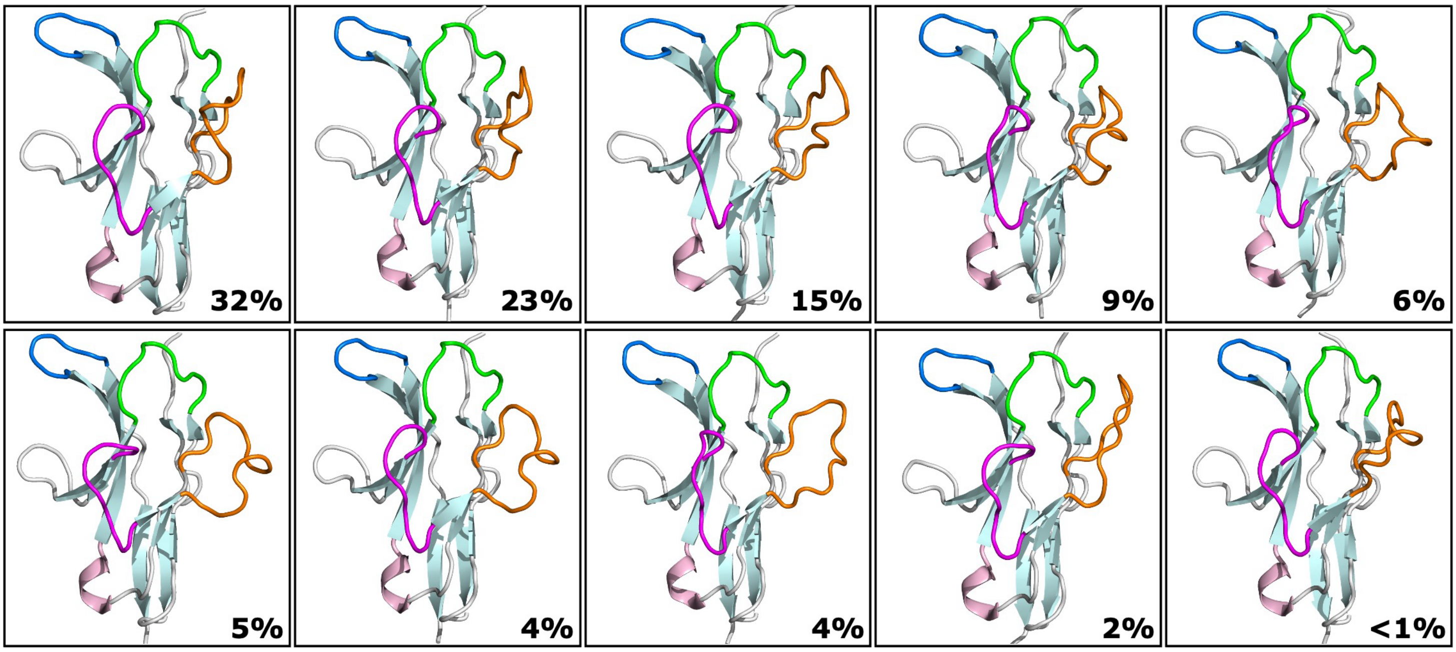


**B**

#### SIRPαV2

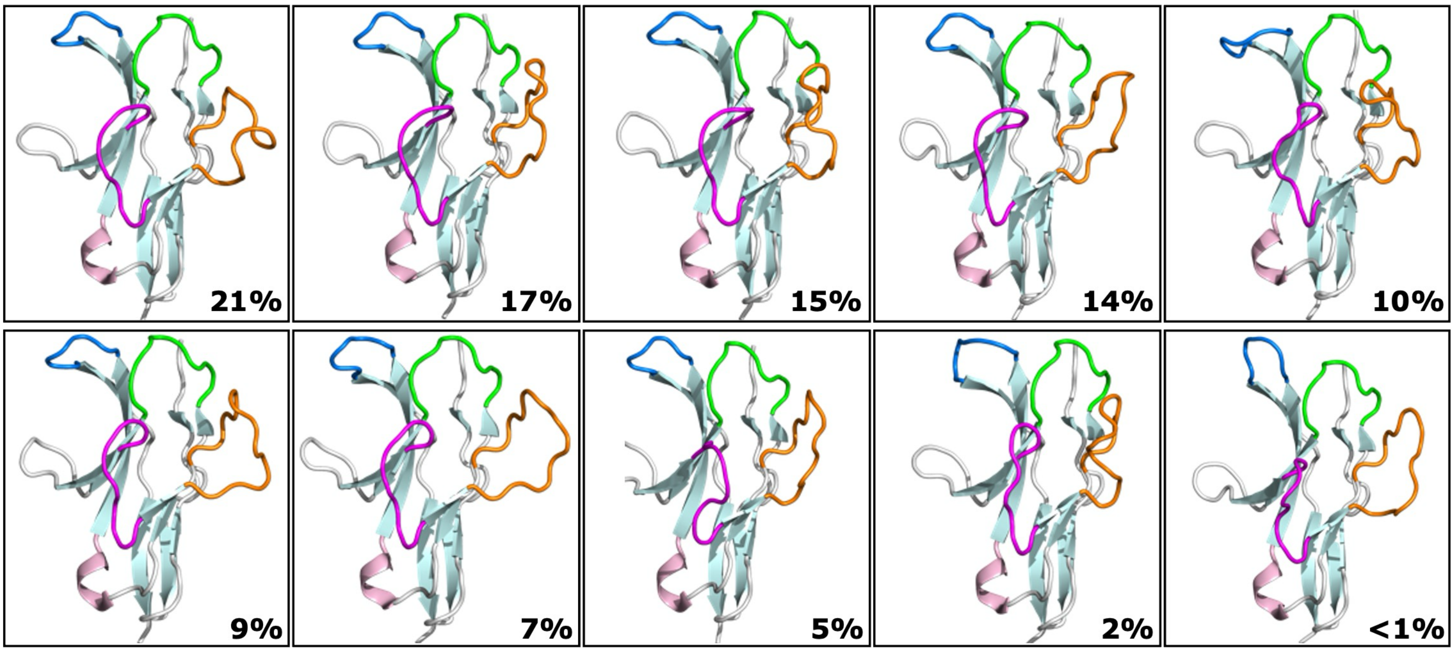

### Figure S12

Binding site opening (Loop distances)

A

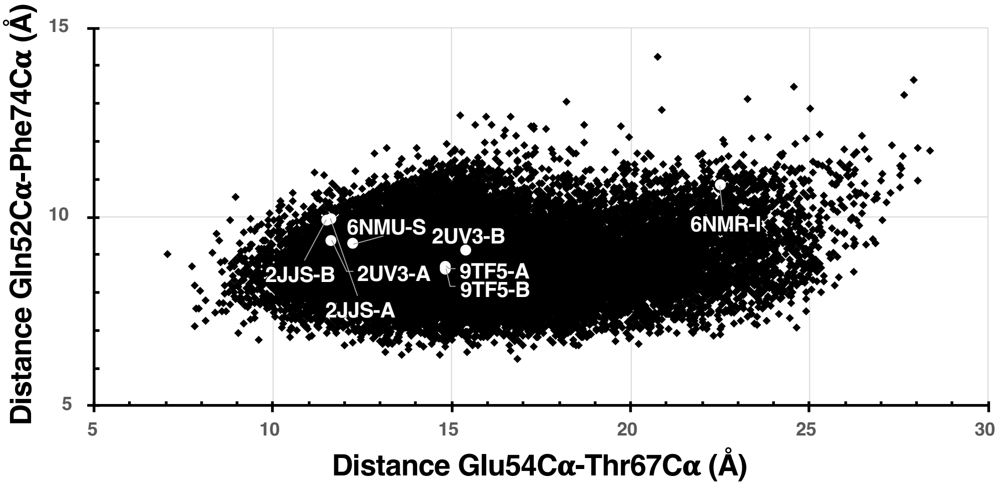


B

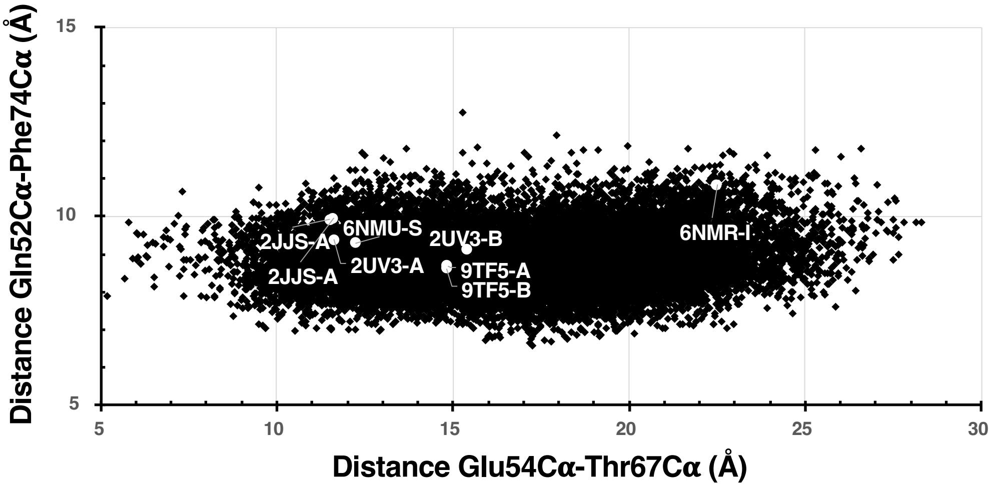

### Figure S13

Binding site pocket

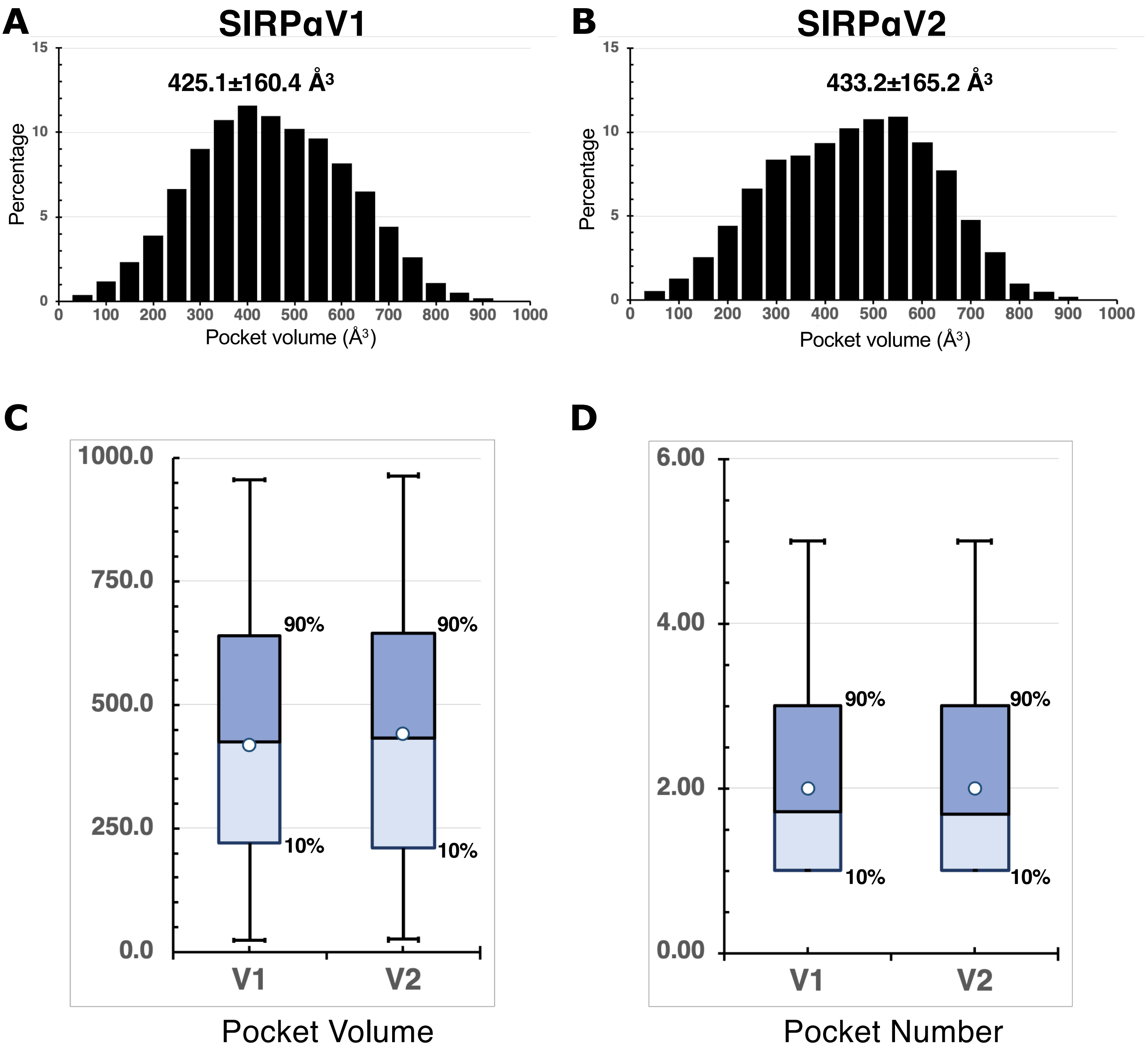

### Figure S14

Phe74 Dihedral angles

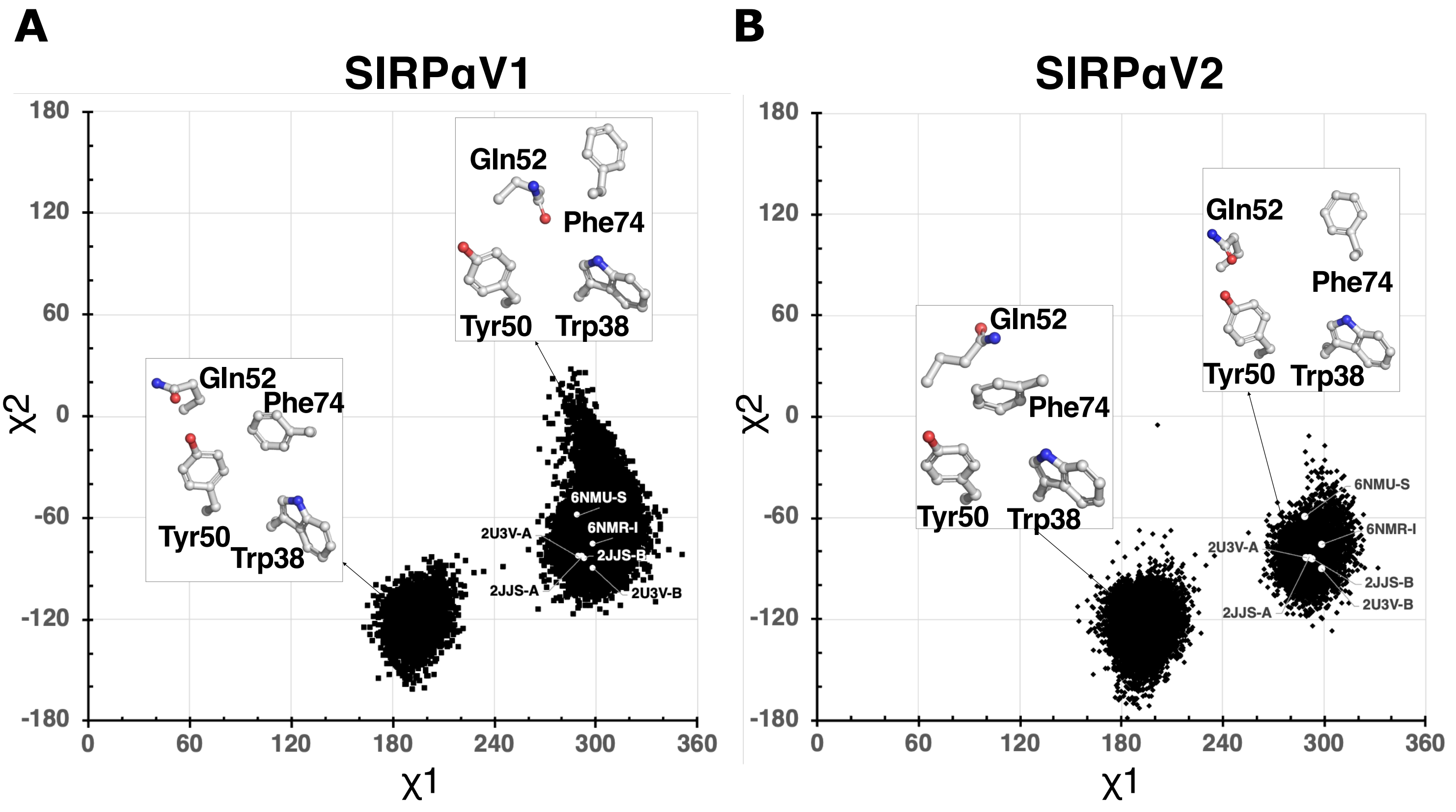

### Figure S15

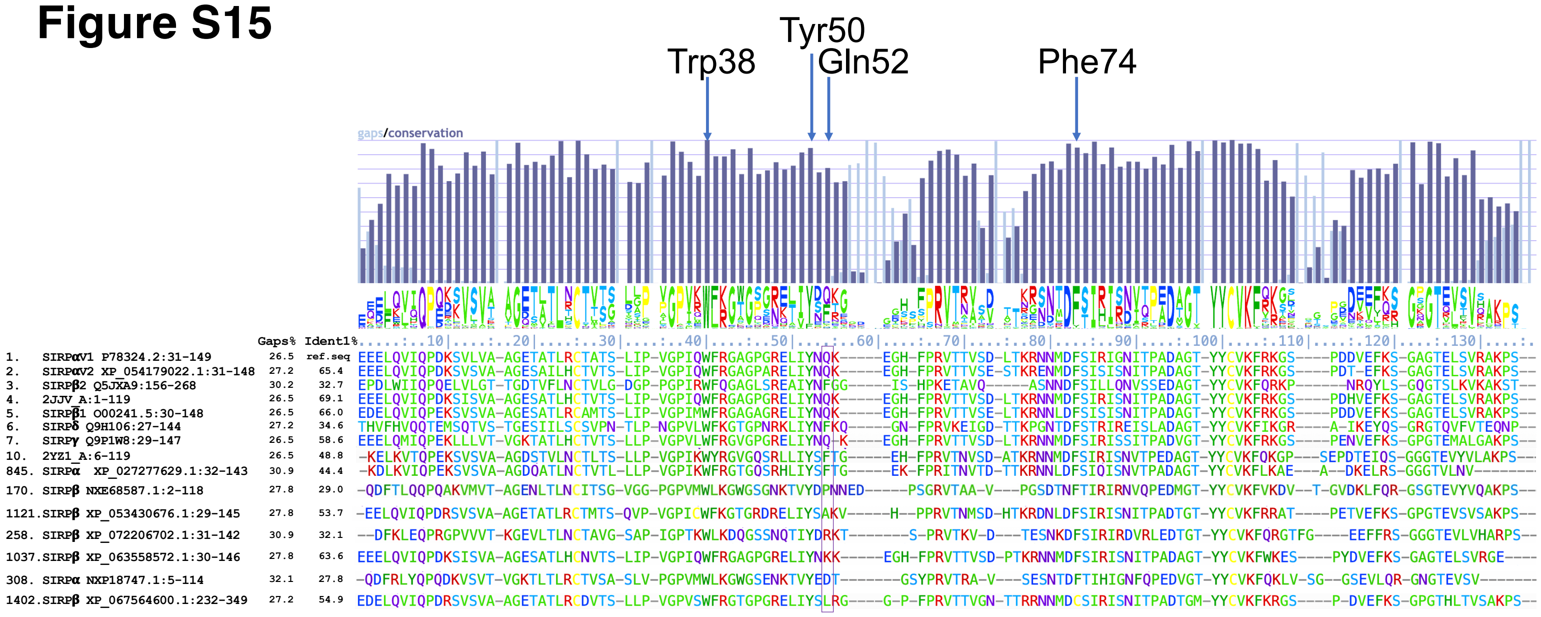

Tyr50
Trp38
Gln52
Phe74
gaps/conservation
Gaps% Ident1% ....:..10|....:..20|....:..30|....:..40|....:..50|....:..60|....:..70|....:..80|....:..90|....:.100|....:.110|....:.120|....:.130|....:.
1. SIRPαV1 P78324.2:31-149 26.5 ref.seq EEELQVIQPDKSVLVA-AGETATLRCTATS-LIP-VGPIQWFRGAGPGRELIYNQK-----EGH-FPRVTTVSD-LTKRNNMDFSIRIGNITPADAGT-YYCVKFRKGS----PDDVEFKS-GAGTELSVRAKPS--
2. SIRPαV2 XP_054179022.1:31-148 27.2 65.4 EEELQVIQPDKSVSVA-AGESAILHCTVTS-LIP-VGPIQWFRGAGPARELIYNQK-----EGH-FPRVTTVSE-STKRENMDFSISISNITPADAGT-YYCVKFRKGS----PDT-EFKS-GAGTELSVRAKPS--
3. SIRPβ2 Q5JXA9:156-268 30.2 32.7 EPDLWIIQPQELVLGT-TGDTVFLNCTVLG-DGP-PGPIRWFQGAGLSREAIYNFGG-----IS-HPKETAVQ-----ASNNDFSILLQNVSSEDAGT-YYCVKFQRKP------NRQYLS-GQGTSLKVKAKST--
4. 2JJV_A:1-119 26.5 69.1 EEELQVIQPDKSISVA-AGESATLHCTVTS-LIP-VGPIQWFRGAGPGRELIYNQK-----EGH-FPRVTTVSD-LTKRNNMDFSIRISNITPADAGT-YYCVKFRKGS----PDHVEFKS-GAGTELSVRAKPS--
5. SIRPβ1 O00241.5:30-148 26.5 66.0 EDELQVIQPEKSVSVA-AGESATLRCAMTS-LIP-VGPIMWFRGAGAGRELIYNQK-----EGH-FPRVTTVSE-LTKRNNLDFSISISNITPADAGT-YYCVKFRKGS----PDDVEFKS-GAGTELSVRAKPS--
6. SIRPδ Q9H106:27-144 27.2 34.6 THVFHVQQTEMSQTVS-TGESIILSCSVPN-TLP-NGPVLWFKGTGPNRKLIYNFKQ-----GN-FPRVKEIGD-TTKPGNTDFSTRIREISLADAGT-YYCVKFIKGR----A-IKEYQS-GRGTQVFVTEQNP--
7. SIRPγ Q9P1W8:29-147 26.5 58.6 EEELQMIQPEKLLLVT-VGKTATLHCTVTS-LLP-VGPVLWFRGVGPGRELIYNQ-K----EGH-FPRVTTVSD-LTKRNNMDFSIRISSITPADVGT-YYCVKFRKGS----PENVEFKS-GPGTEMALGAKPS--
10. 2YZ1_A:6-119 26.5 48.8 -KELKVTQPEKSVSVA-AGDSTVLNCTLTS-LLP-VGPIKWYRGVGQSRLLIYSFTG-----EH-FPRVTNVSD-ATKRNNMDFSIRISNVTPEDAGT-YYCVKFQKGP---SEPDTEIQS-GGGTEVYVLAKPS--
845. SIRPα XP_027277629.1:32-143 30.9 44.4 -KDLKVIQPEKSVSVA-AGDQATLNCTVTL-LLP-VGPIKWFRGTGQSRHLIYSFTG-----EK-FPRITNVTD-TTKRNNLDFSIQISNVTPADAGT-YYCVKFLKAE---A--DKELRS-GGGTVLNV-------
170. SIRPβ NXE68587.1:2-118 27.8 29.0 -QDFTLQQPQAKVMVT-AGENLTLNCITSG-VGG-PGPVMWLKGWGSGNKTVYDPNNED-----PSGRVTAA-V---PGSDTNFTIRIRNVQPEDMGT-YYCVKFVKDV--T-GVDKLFQR-GSGTEVYVQAKPS--
1121.SIRPβ XP_053430676.1:29-145 27.8 53.7 -EELQVIQPDRSVSVA-AGETATLRCTMTS-QVP-VGPICWFKGTGRDRELIYSAKV-----H--PPRVTNMSD-HTKRDNLDFSIRISNITPADTGT-YYCVKFRRAT----PETVEFKS-GPGTEVSVSAKPS--
258. SIRPβ XP_072206702.1:31-142 30.9 32.1 --DFKLEQPRGPVVVT-KGEVLTLNCTAVG-SAP-IGPTKWLKDQGSSNQTIYDRKT-------S-PRVTKV-D---TESNKDFSIRIRDVRLEDTGT-YYCVKFQRGTFG----EEFFRS-GGGTEVLVHARPS--
1037.SIRPβ XP_063558572.1:30-146 27.8 63.6 EEELQVIQPDKSISVA-AGESATLHCNVTS-LIP-VGPIQWFRGAGPGRELIYNKK-----EGH-FPRVTTVSD-PTKRNNMDFSIRISNITPADAGT-YYCVKFWKES----PYDVEFKS-GAGTELSVRGE----
308. SIRPα NXP18747.1:5-114 32.1 27.8 -QDFRLYQPQDKVSVT-VGKTLTLRCTVSA-SLV-PGPVMWLKGWGSENKTVYEDT-------GSYPRVTRA-V---SESNTDFTIHIGNFQPEDVGT-YYCVKFQKLV-SG--GSEVLQR-GNGTEVSV-------
1402.SIRPβ XP_067564600.1:232-349 27.2 54.9 EDELQVIQPDRSVSVA-AGETATLRCDVTS-LLP-VGPVSWFRGTGPGRELIYSLRG----G-P-FPRVTTVGN-TTRRNNMDCSIRISNITPADTGM-YYCVKFKRGS----P-DVEFKS-GPGTHLTVSAKPS--

### Figure S16

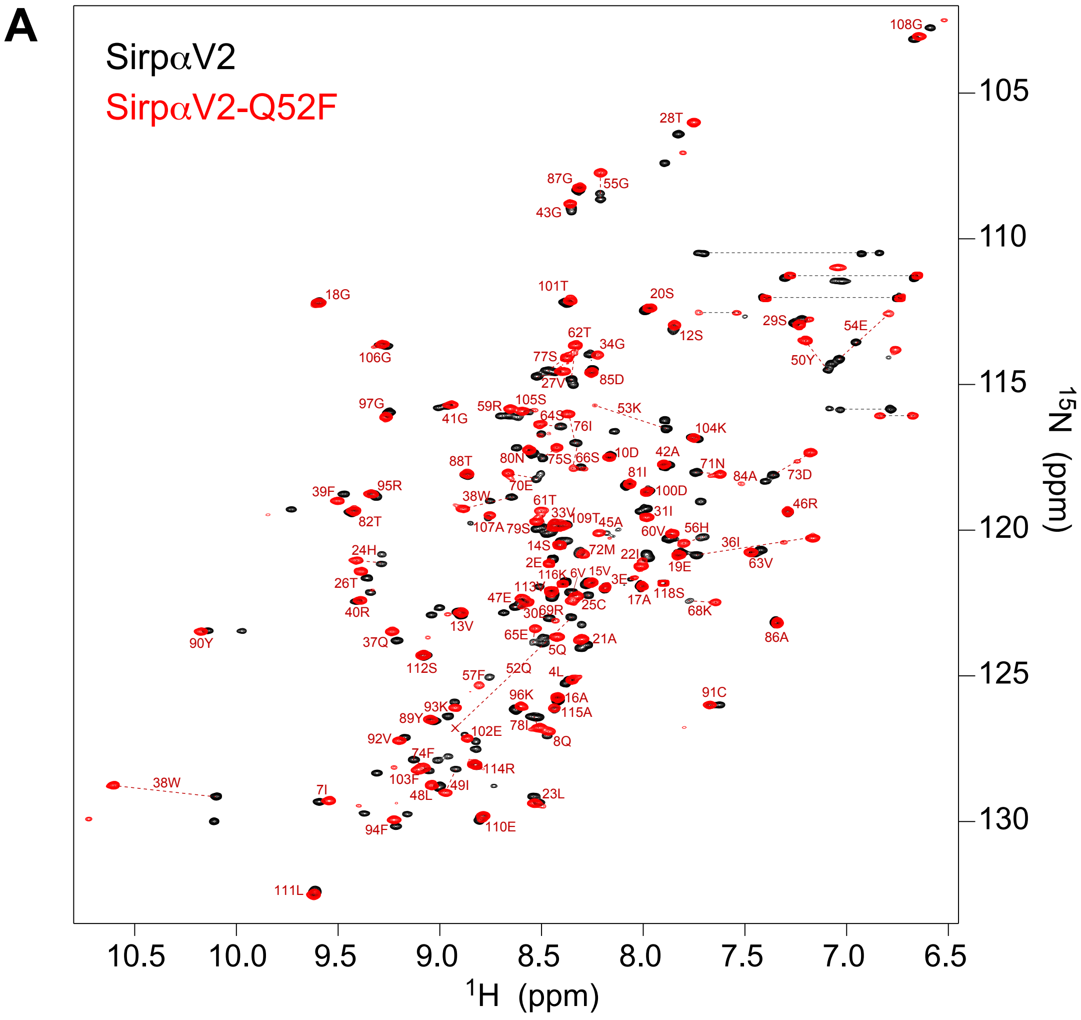
A
SirpαV2
SirpαV2-Q52F
105
110
115
120
125
130
15N (ppm)
10.5
10.0
9.5
9.0
8.5
8.0
7.5
7.0
6.5
1H (ppm)

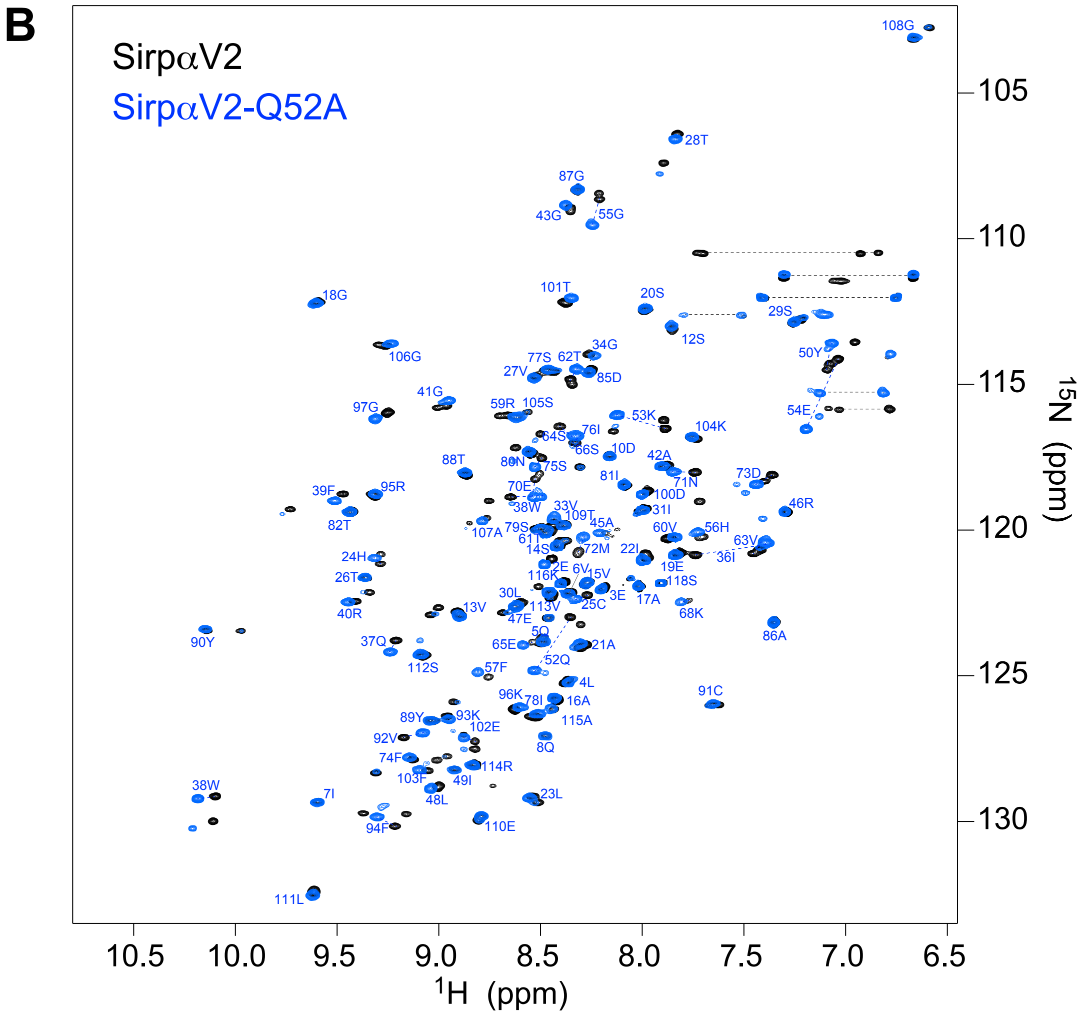
B
SirpαV2
SirpαV2-Q52A
105
110
115
120
125
130
15N (ppm)
10.5
10.0
9.5
9.0
8.5
8.0
7.5
7.0
6.5
1H (ppm)

### Figure S17

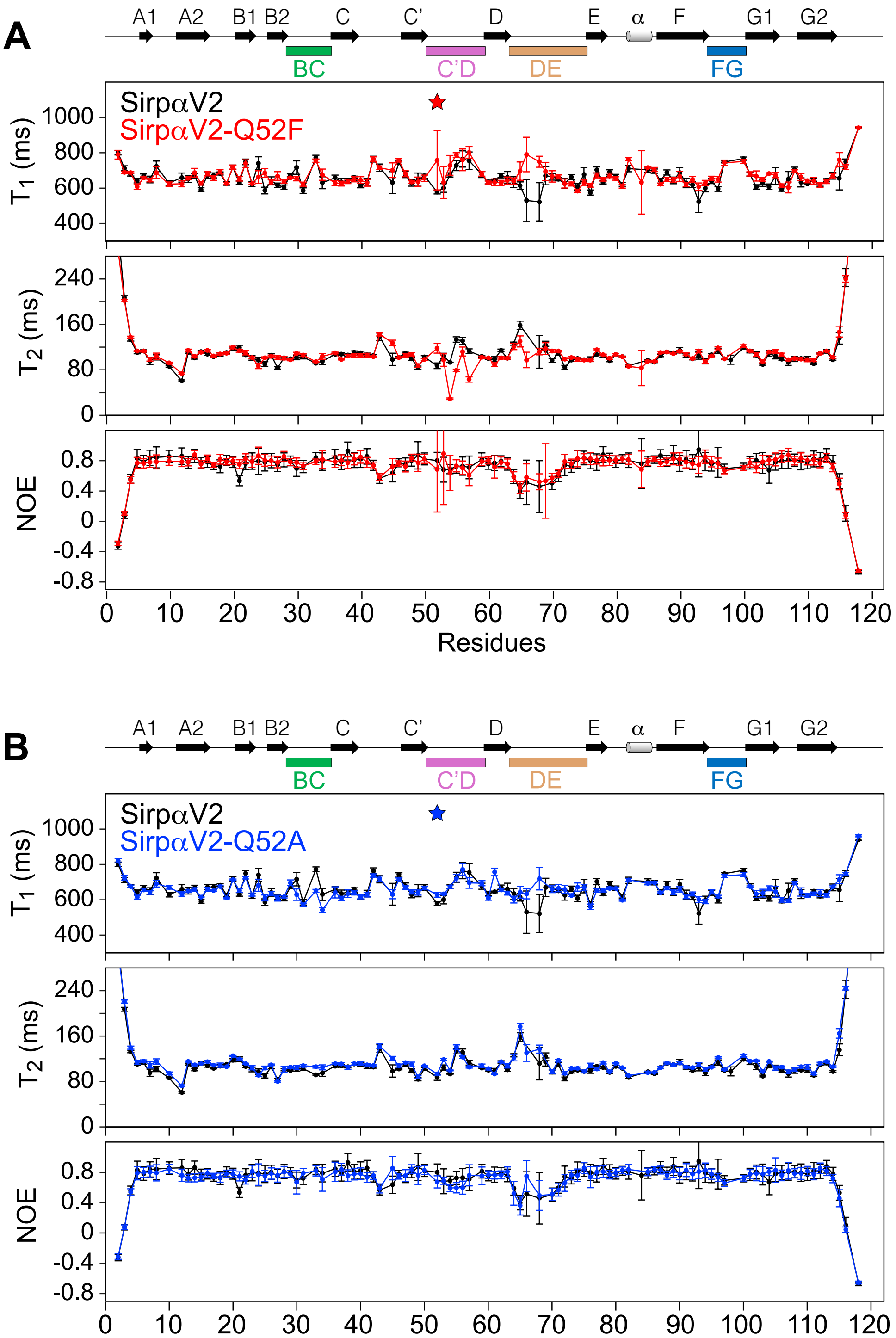
A
A1
A2
B1
B2
C
C'
D
E
α
F
G1
G2
BC
C'D
DE
FG
SirpαV2
SirpαV2-Q52F
T1 (ms)
T2 (ms)
NOE
Residues
B
SirpαV2
SirpαV2-Q52A

**Figure S18**
(Gln52 mutants)

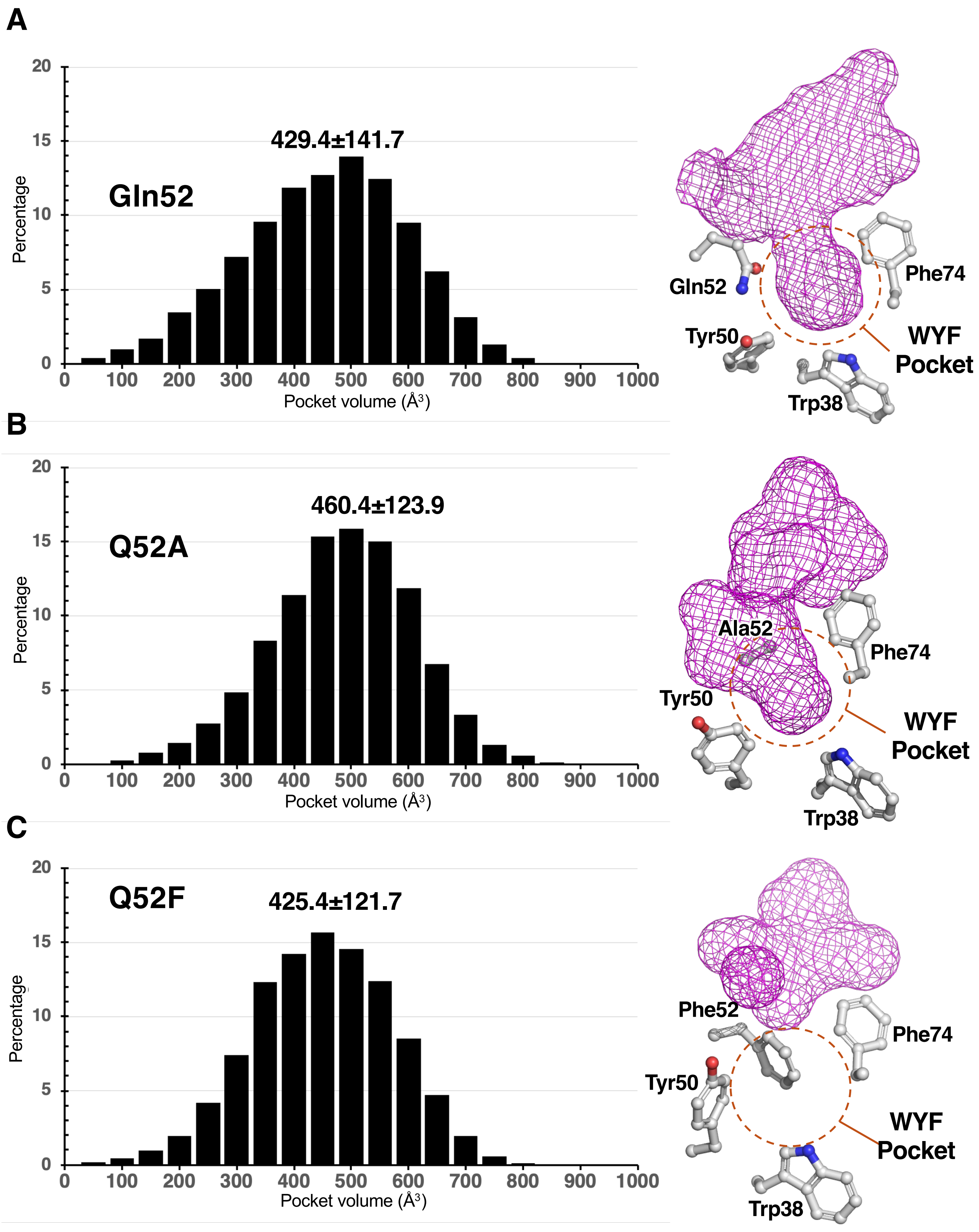

### Figure S19

A B

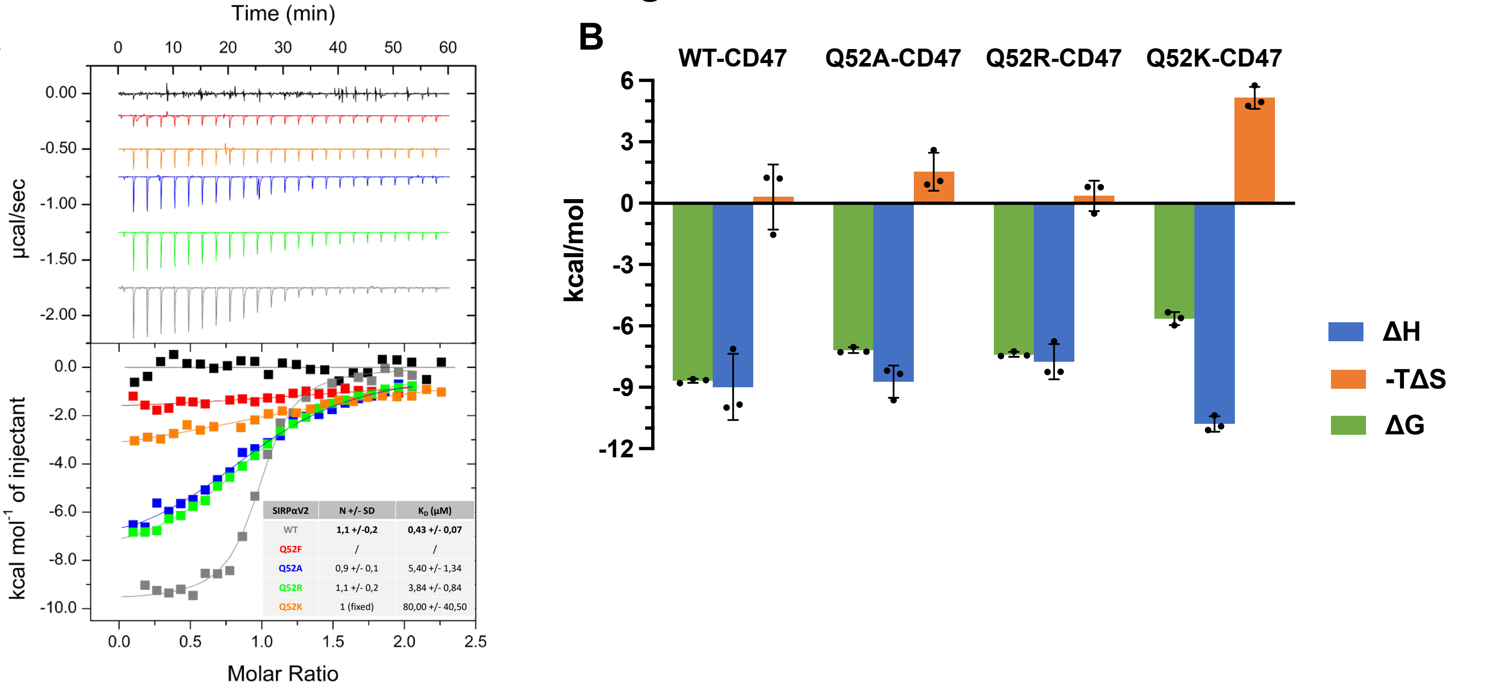


C

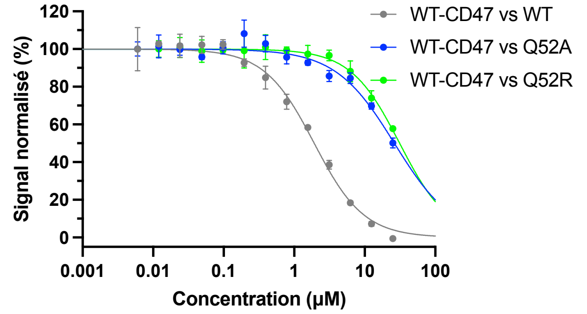


| | WT-CD47 vs WT | WT-CD47 vs Q52A | WT-CD47 vs Q52R |
|---|---|---|---|
| IC50 | 1.921 | 26.30 | 31.24 |
| Hillslope | 1.230 | 1.038 | 1.246 |

D

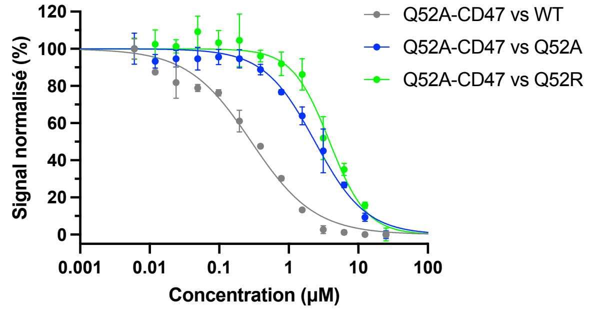


| | Q52A-CD47 vs WT | Q52A-CD47 vs Q52A | Q52A-CD47 vs Q52R |
|---|---|---|---|
| IC50 | 0.2885 | 2.378 | 3.861 |
| Hillslope | 0.9519 | 1.181 | 1.616 |

E

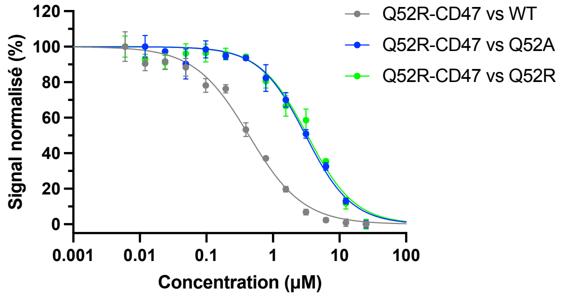


| | Q52R-CD47 vs WT | Q52R-CD47 vs Q52A | Q52R-CD47 vs Q52R |
|---|---|---|---|
| IC50 | 0.4358 | 3.093 | 3.304 |
| Hillslope | 1.066 | 1.256 | 1.223 |

F

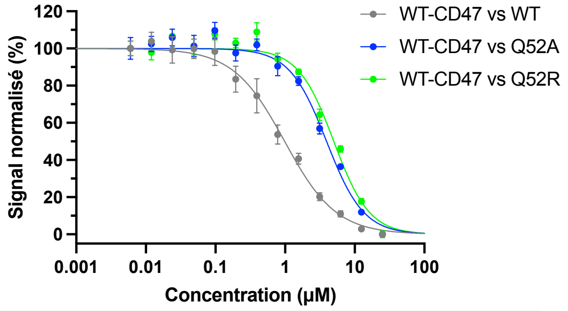


| | WT-CD47 vs WT | WT-CD47 vs Q52A | WT-CD47 vs Q52R |
|---|---|---|---|
| IC50 | 0.9849 | 3.940 | 5.004 |
| Hillslope | 1.175 | 1.628 | 1.702 |

G

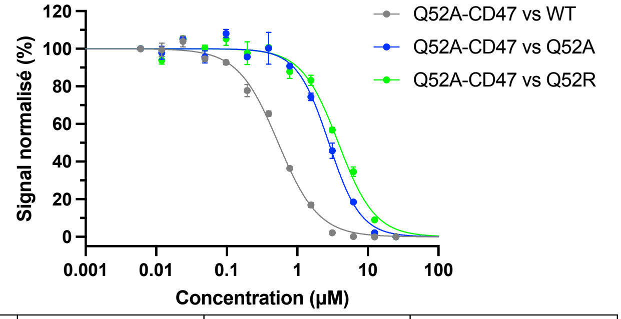


| | Q52A-CD47 vs WT | Q52A-CD47 vs Q52A | Q52A-CD47 vs Q52R |
|---|---|---|---|
| IC50 | 0.5363 | 2.797 | 3.817 |
| Hillslope | 1.524 | 1.926 | 1.673 |

H

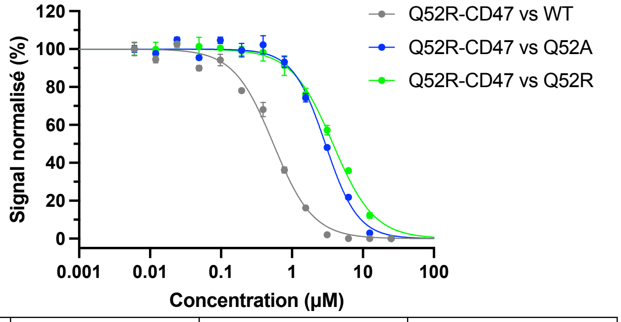


| | Q52R-CD47 vs WT | Q52R-CD47 vs Q52A | Q52R-CD47 vs Q52R |
|---|---|---|---|
| IC50 | 0.5470 | 2.954 | 3.783 |
| Hillslope | 1.550 | 1.894 | 1.496 |

### Figure S20

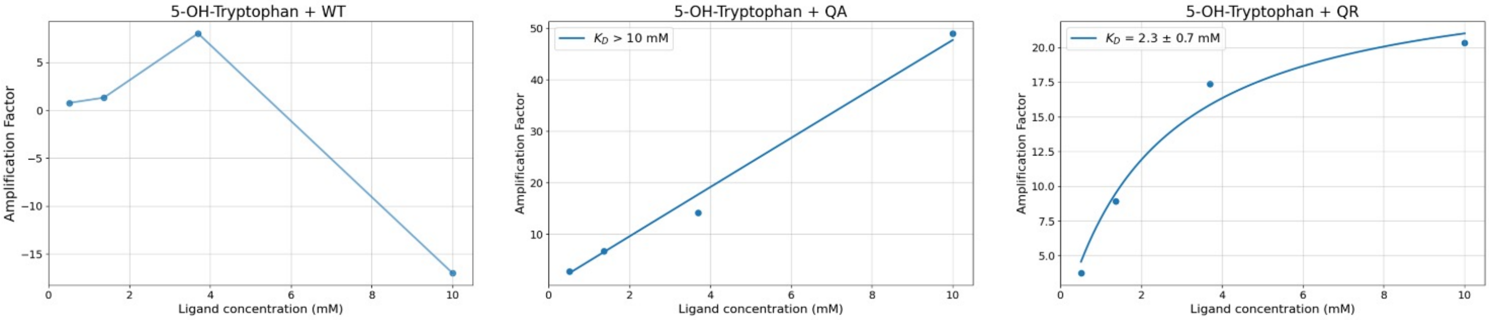

5-OH-Tryptophan + WT
Amplification Factor
Ligand concentration (mM)
5-OH-Tryptophan + QA
$K_D$ > 10 mM
Amplification Factor
Ligand concentration (mM)
5-OH-Tryptophan + QR
$K_D$ = 2.3 ± 0.7 mM
Amplification Factor
Ligand concentration (mM)

### Figure S21

Gln52 Dihedral angles

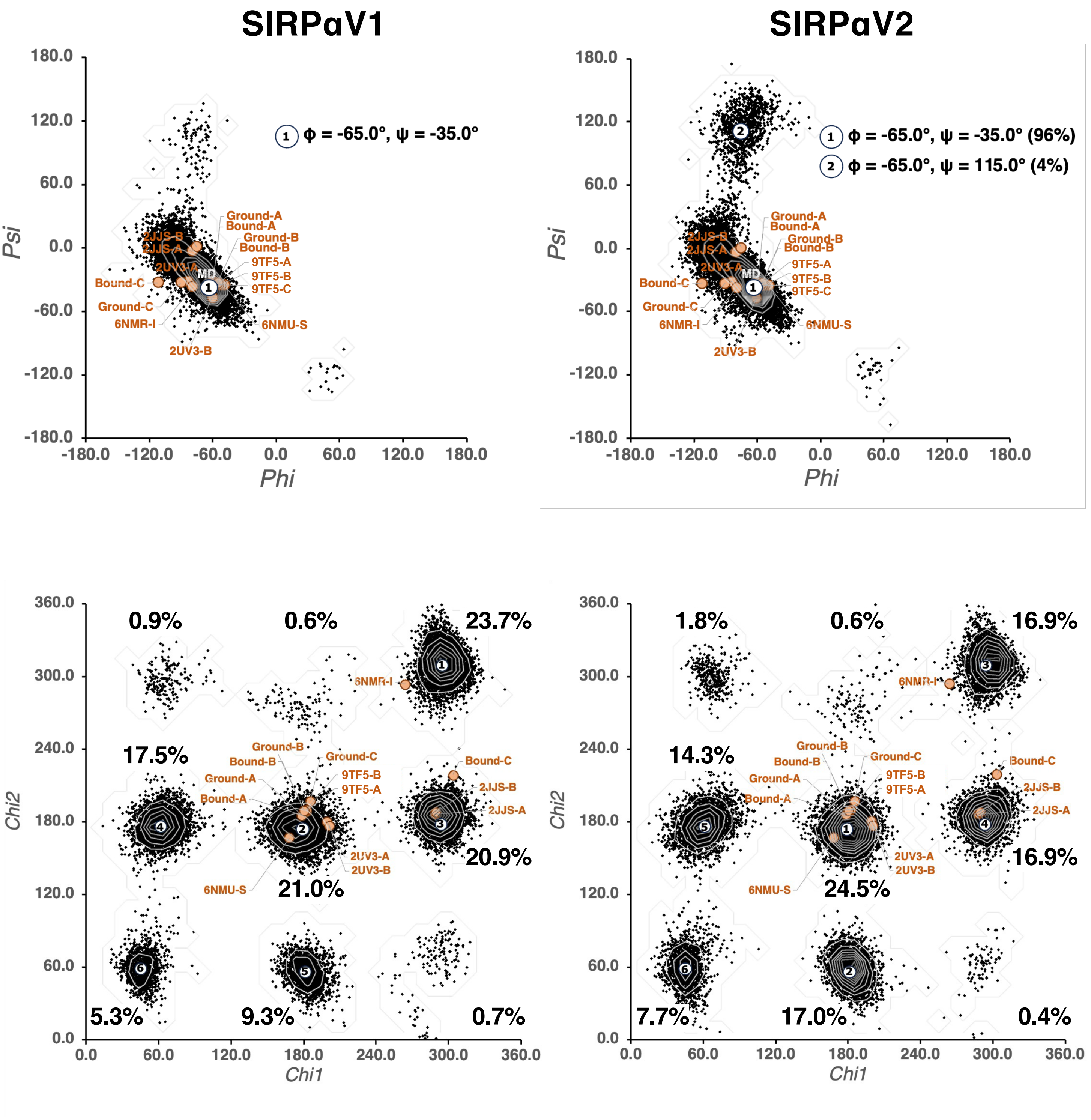
